# Force field parameters for Fe^2+^_4_S^2-^_4_ clusters of dihydropyrimidine dehydrogenase, the 5-fluorouracil cancer drug deactivation protein: a step towards *in silico* pharmacogenomics studies

**DOI:** 10.1101/2021.04.20.440516

**Authors:** Maureen Bilinga Tendwa, Lorna Chebon-Bore, Kevin Lobb, Thommas Mutemi Musyoka, Özlem Tastan Bishop

## Abstract

The dimeric dihydropyrimidine dehydrogenase (DPD) metalloenzyme, an adjunct anti-cancer drug target contains highly specialized 4 × Fe^2+^_4_S^2-^_4_ clusters per chain. These clusters facilitate the catalysis of the rate-limiting step in the pyrimidine degradation pathway through a harmonized electron transfer cascade that triggers a redox catabolic reaction. In the process, majority of administered 5-fluorouracil (5-FU) cancer drug is inactivated while a small proportion is activated to nucleic acid antimetabolites. The occurrence of missense mutations in DPD protein within the general population, including those of African descent, has adverse toxicity effects due to altered 5-FU metabolism. Thus, deciphering mutation effects on protein structure and function is vital, especially for precision medicine purposes. We previously proposed combined molecular dynamics (MD) and dynamic residue network (DRN) analysis to decipher the molecular mechanisms of missense mutations in other proteins. However, the presence of Fe^2+^_4_S^2-^_4_ clusters in DPD poses a challenge for such *in silico* studies. The existing AMBER force field parameters cannot accurately describe the Fe^2+^ center coordination exhibited by this enzyme. Therefore, this study aimed to derive AMBER force field parameters for DPD enzyme Fe^2+^ centers, using the original Seminario method and collation features Visual Force Field Derivation Toolkit as a supportive approach. All-atom MD simulations were performed to validate the results. Both approaches generated similar force field parameters which accurately described the human DPD protein Fe^2+^_4_S^2-^_4_ clusters architecture. This information is crucial and opens new avenues for *in silico* cancer pharmacogenomics and drug discovery related research on 5-FU drug efficacy and toxicity issues.

## 1. Introduction

Dihydropyrimidine dehydrogenase (DPD; EC 1.3.1.2) is the initial rate-limiting enzyme in the triple-step pyrimidine-based catabolic pathway [1, 2]. The enzyme is involved in the degradation of pyrimidine bases (thymine and uracil) via a NADPH-dependent reaction to 5,6-dihydrothymine and 5,6 dihydrouracil, respectively [1]. Besides its biological nucleotide catabolizing function, the enzyme is an adjunct anti-cancer drug target [3]. It is solely responsible for phase 1 metabolism of 5-fluorouracil (5-FU), a commonly prescribed pyrimidine-like anti-cancer drug. During 5-FU metabolism, majority (80-85 %) of the administered dose is rapidly degraded to dihydroflourouracil (DHFU), the inactive form. Additionally, a small proportion (1-3 %) of the administered drug is activated to fluorodeoxyuridine monophosphate (FdUMP) and fluorouridine triphosphate (FUTP), leading to inhibition of the DNA synthesis and RNA processing. The remaining 12-19 % unmetabolized 5-FU form is excreted through urine [4]. As such, deficiency or reduced levels of DPD enzyme as well as sequence variation due to mutations have been reported to cause fluoropyrimidine associated toxicity effects [5, 6]. Thus, understanding the implication of mutations on the catalytic mechanism of DPD can improve treatment approaches for oncology patients. Although there is a growing interest in molecular investigations on DPD enzyme, especially via computational approaches such as molecular dynamics (MD) simulations [7], a significant hindrance against the implementation of such studies is the presence of four iron-sulfur (Fe^2+^_4_S^2-^_4_) clusters in the homodimeric form of DPD structure, which require additional force field parameters.

### 1.1. DPD structure and mechanism of action

In this study, our interest is in the human DPD protein. However, since we used the crystal structure of the pig homolog to build the 3D human model, we will define the structural features from the template structure.

The 222 kDa homodimeric structure of a pig DPD (PDB ID: 1H7X) [2, 8] enzyme consists of one ligand (5-FU represented as URF), a cofactor (nicotinamide adenine dinucleotide phosphate [NADPH]), two protein-bound organic cofactors (flavin adenine dinucleotide [FAD] and flavin mononucleotide [FMN]) and four inorganic Fe^2+^_4_S^2-^_4_ clusters. Each 1020 residues monomers has five domains; domain one (residues 27 - 173, 2 × Fe^2+^_4_S^2-^_4_ clusters); domain two (residues 174 - 286 and 442 - 524) and three (residues 287 - 441) are the NADPH- and FAD-binding domains respectively; domain four (FMN, URF residue 535 - 847; active - site loop residues 675 -679) and domain five (residues 1 - 26 and 848 - 1017; 2 × Fe^2+^_4_S^2-^_4_) (Figure 1)[1, 2, 9, 10]. Additionally, the two Fe^2+^_4_S^2-^_4_ (hetero atoms 1028 and 1029) clusters in domain four of the same chain are in very close proximity to domain five Fe^2+^_4_S^2-^_4_ (hetero atoms 1026 and 1027) clusters of the opposite chain [11, 12]. The FMN/pyrimidine binding domain of each chain is closely positioned to the corresponding C terminal domain (2 × Fe^2+^_4_S^2-^_4_ clusters) [1, 2, 9]. This arrangement is crucial for the electron transfer pathway from the NADPH donor molecule to pyrimidine binding sites [1, 2, 9]. However, the exact mechanism of how these redox cofactors participate in the reaction is largely unknown [10]. Previous studies have indicated that the Fe^2+^_4_S^2-^_4_ clusters tend to form a bridge between FMN and FAD cofactors for electron transport to the active site [9, 12-14].

**Figure 1.**
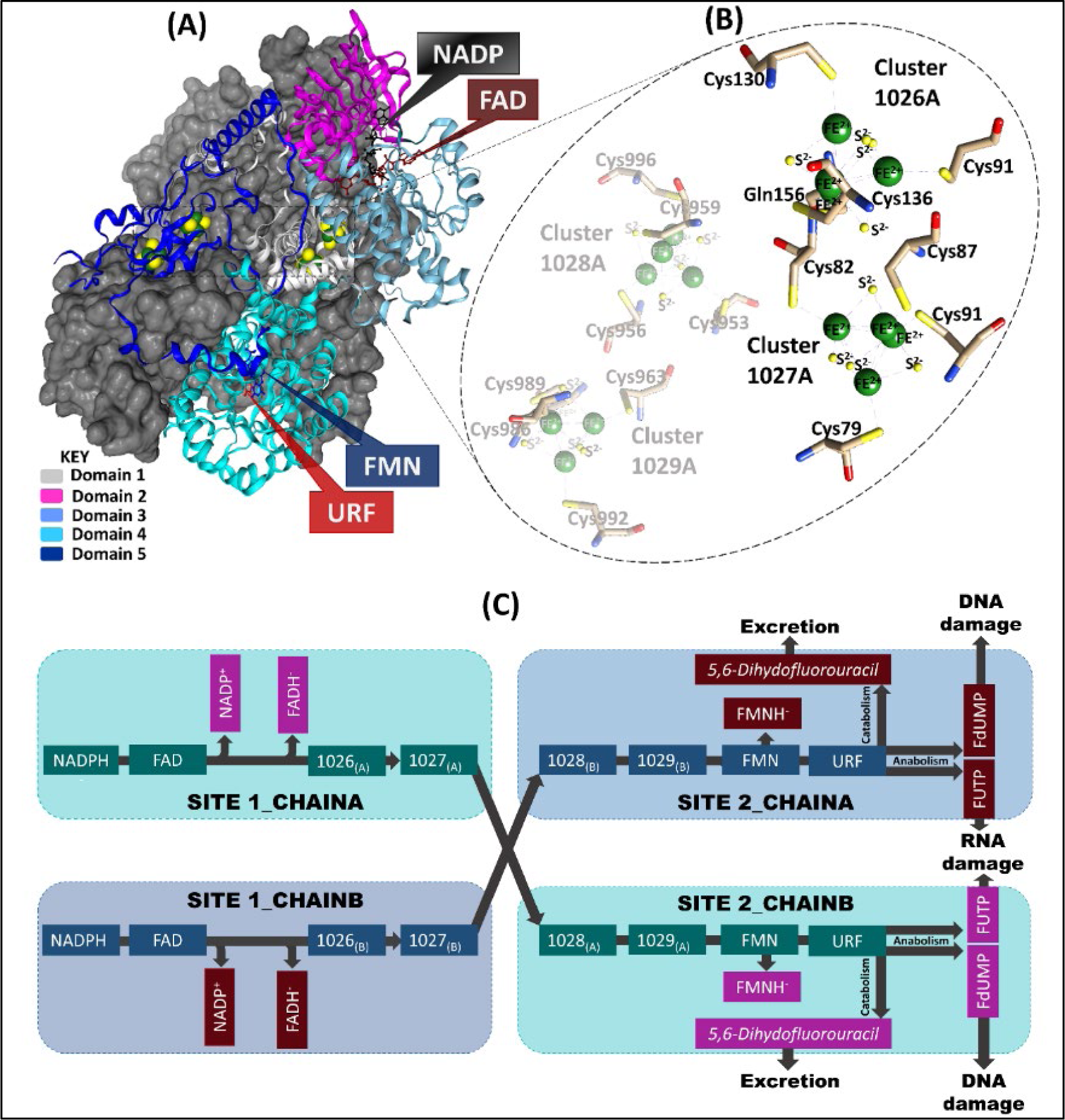
Comprehensive representation of chain-A and chain-B of the pig DPD template (PDB ID: 1H7X) crystal structure. **A)** Chain-A domains (1-5) are colored (teal, magenta, light grey, light and dark blue respectively) and the cofactors are represented as sticks. The grey surface represents chain-B which is a mirror image of chain-A. **B)** Highlights the Fe^2+^_4_S^2-^_4_ clusters coordinating environment of chain-A. **C)** The electron transport process in which 2 electrons are lost from nicotinamide-adenine-dinucleotide phosphate (NADPH) via flavin adenine dinucleotide (FAD) and Fe^2+^_4_S^2-^_4_ (1026 and 1027) clusters in site 1 of both chains, for the reduction of URF (5-FU) in site 2 of the opposite chain via Fe^2+^_4_S^2-^_4_ (1028 and 1029) clusters and flavin mononucleotide (FMN).

The Fe^2+^_4_S^2-^_4_ clusters manifest a distorted tetrahedron cubane-like geometry [1, 2, 15]. Each Fe^2+^ in three of the clusters (1027, 1028, and 1029) is coordinated by cysteine residues and are connected by disulfide bridges ([Fe^2+^_4_S^2-^_4_ (S-Cys)^4^]). Although cluster 1026 depicts unique coordination in which four Fe^2+^ atoms are inter-connected by disulfide bridges, three of which are bound to the protein backbone by cysteine residue side chain while the fourth one is bound via glutamine residue ([Fe^2+^_4_S^2-^_4_ (S-Cys)_3_(S-Gln)]) side chain [1, 2, 9, 10, 15].

### 1.2. The study

Although there is an increased interest in the protein metal interactions prompted by the essential physiological roles played by metal ions [16-18], the Fe–S (Gln) coordination in cluster 1026 is yet to be reported in other Fe^2+^_4_S^2-^_4_ cluster containing proteins [1, 2]. Metal ions such as iron (Fe^2+^) are crucial components of a protein’s electron transportation as they trigger the activation process in the catalytic subunit. Additionally, they play important stabilization and homeostatic functions in a protein [19]. As a result, these metal ions form a highly organized geometric arrangement with specific highly conserved residues [20].

We can gain insights into metal coordinating environments through computational studies, especially via molecular dynamics (MD) simulations. However, MD calculations are highly dependent on force fields derived through quantum mechanics (QM), and molecular mechanics (MM) approaches [21, 22]. MM methods employ classical type models to predict the amount of energy in a molecule based on its conformation [23]. These methods are less complicated and highly efficient. Unfortunately, most of the MM force field cannot accurately describe the metal environment since they do not consider the explicit electronic degree of freedom [24]. As a mitigating factor, QM calculations have been employed to adequately account for precise electron structure of atoms [25-27]. Hitherto, various methods such as polarization model, non-bonded, semi-bonded, and bonded have been implemented to characterize metalloproteins. The non-bonded model uses non-covalent (van der Waals and electrostatic forces) interaction to define metal centers [28, 29] whereas, semi-bonded [30, 31] models put dummy atoms around metals to resemble electron orbitals. However, these two methods are unable to take into account charge transfer and polarization effects [32]. These shortcomings have been solved by incorporating the charge transfer and polarization effects in potential energy function models [33, 34]. Contrastingly, the bonded model utilizes defined harmonic energy terms, which have been introduced into possible energy function to account for the bond formation between atoms and metal centers [28, 34, 35]. The approaches mentioned above have extensively been used in studies to characterize Fe^2+^ centers in a range of metalloproteins [36- 39]. Among other Fe^2+^ clusters, Carvalho and colleagues [36] satisfactorily generated AMBER force field parameters for Fe^2+^_4_S^2-^_4_ clusters coordinated by cysteine residues. However, none of these parameters featured glutamine residue coordination to Fe^2+^ center or developed parameters for structures of composite multiple clusters, besides applying two approaches. To the best of our knowledge, this is the first study to determine the human DPD protein metal parameters.

Collectively, the current study integrates MM with QM techniques to determine accurate force field parameters for 8 × Fe^2+^_4_S^2-^_4_ cluster complexes of the modeled human DPD proteins. We utilized the bonded method of QM and Seminario techniques in our calculations [40]. More specifically, the density functional theory (DFT) of the QM approach was used to derive Fe^2+^ center AMBER parameters for two models using different Seminario methods. The first method (*viz*. Model 1) used the original Seminario [41] method [42] whereas the second method (*viz*. Model 2) used collation features Visual Force Field Derivation Toolkit (VFFDT) Seminario [43]. A comparison of the parameters from the two methods was performed and their reliability evaluated via all atom MD simulations. For the first time, the current study reports novel force field parameters for multiple Fe^2+^_4_S^2-^_4_ clusters, coordinated to both cysteine and glutamine residues. Furthermore, the reliability of the two parameter generation approaches was also evaluated and found to be comparable. The newly derived force field parameters can be adopted by other systems depicting a similar Fe^2+^ coordinating environment. More importantly, establishment of these parameters creates an avenue for further molecular studies to fully understand the functional mechanism of the human DPD protein, and to decipher the effects of missense mutations on drug metabolism and cancer drug toxicity issues. As part of our ongoing investigations about the effects of known variants in human DPD especially on its structure and stability, the reliability of the current parameters has been confirmed and the findings will be published as a follow-up study. Furthermore, different methods such as identification of new mutants coupled with the structural analysis and clinical studies, i.e. phenotyping of DPD, has a great impact on understanding the structural and functional effects of these mutations [6]. Together, these results would be crucial not only towards understanding how mutations lead to 5-FU toxicities but also inform better the implementation of precision medicine in cancer treatment.

## 2. Results and Discussion

### 2.1. Human DPD 3D wild type (WT) complete structure determined via homology modeling approaches

The availability of accurate and complete 3D structural information is fundamental aspect for molecular studies aimed at understanding protein function. With the absence of the human DPD X-ray structure in the protein data bank (PDB) [8], homology modeling approaches were used to calculate accurate models of the human DPD enzyme using MODELLER v9.15 [44], DiscoveryStudio4.5 [45] and pig X-ray structure (PDB ID: 1H7X, 2.01 Å) as template [1, 2]. The choice of the template was guided by the high sequence identity (93%) with the target human DPD enzyme. Additionally, it was in complex with the drug of interest (5-FU) and had a complete query coverage of 100%. Using the very slow refinement level in MODELLER v9.15, 100 apo protein models were generated. The three best models with the lowest z-DOPE scores of -1.36, -1.36 and -0.88, were chosen for further validation. z-DOPE score evaluates the closeness of a model in comparison with the native structure based on atomic distance-displacement statistical potential with a score of ≤ -1.0 being considered as a near-native structure [46, 47]. Consequently, holo (apo and cofactors) and holo-drug (5-FU) complexes structures were generated by incorporating the non-protein coordinates from the template in Discovery Studio 4.5 [45]. Additional model quality assessment (Table S1) was performed using VERIFY3D webserver [48], qualitative model energy analysis (QMEAN) [49], Protein Structure Analysis (ProSA) [50] and program to check the stereochemical quality of protein structures (PROCHECK) [51]. VERIFY3D utilizes pairwise interaction derived energy potentials to evaluate the local quality of a model based on each residue structure environment [48]. High-quality structures are predicted to have more than 80% of their residues with a 3D-1D score of 0.2 or higher [48]. The modeled structures had 3D-ID scores of 0.2 or higher (Table S1) in 85.01 of its residues. QMEAN estimates the quality of the submitted model based on its physicochemical properties, then derives a value corresponding to the overall quality of the structure and compares it to the calculated QMEAN-scores of 9766 high-resolution experimental structures [49]. The modelled structures of DPD holo and holo-drug complexes had a QMEAN-score of 0.90 and 0.89, which is similar to that of high-resolution experimental structures. ProSA assesses the quality of the submitted model by calculating its potential energy and comparing the resulting score to that of the experimental structures available in PDB [50]. At least each monomer of the holo and holo-drug complexes Z-score was between -13.41 to -13.56, which is similar to that of NMR structures of the same size.

PROCHECK assesses the stereochemical quality of the submitted protein models based on their phi/psi angle arrangement, and then produce Ramachandran plots which show the protein residues positions on most favored, allowed, and disallowed regions [51]. Each generated model had more than 83.8%, 16.0% and 0.2% of their residues in the most favored, allowed and disallowed regions, respectively, suggesting a good distribution of torsion angles [Table S1]. Overall, constructed holo and holo-drug complexes with consistently high-quality scores were obtained.

To remove steric clashes in the generated models (holo and holo-drug), 100 steps of minimization with steepest descent algorithm using GROMACS 5.14 MD simulation package [52] were performed and determined to be suitable for subsequent calculations.

### 2.2. AMBER force field parameters generated using bonded approaches

The metal coordination geometries in proteins are highly dependent on the protonation states of the residues involved. Thus, to achieve correct geometry arrangements in the human DPD protein, the protonation states of all titratable resides was determined at a pH of 7.5 using the H++ webserver (http://biophysics.cs.vt.edu/H++) [53] (Table S2). To ensure correct protonation, a visual inspection of all titratable residues was performed and corrected using Schrödinger Maestro version 11.8 [54]. Table 1 shows the protonation states of residues forming a bond with the metal ions in the Fe^2+^_4_S^2-^_4_ clusters. Cys protonated as CYM and interacted with the Fe^2+^ center through sulfur (SG) bond. On the other hand, Gln was protonated as GLH to coordinate with Fe^2+^ ion through oxygen (OE) atom.

**Table 1.**
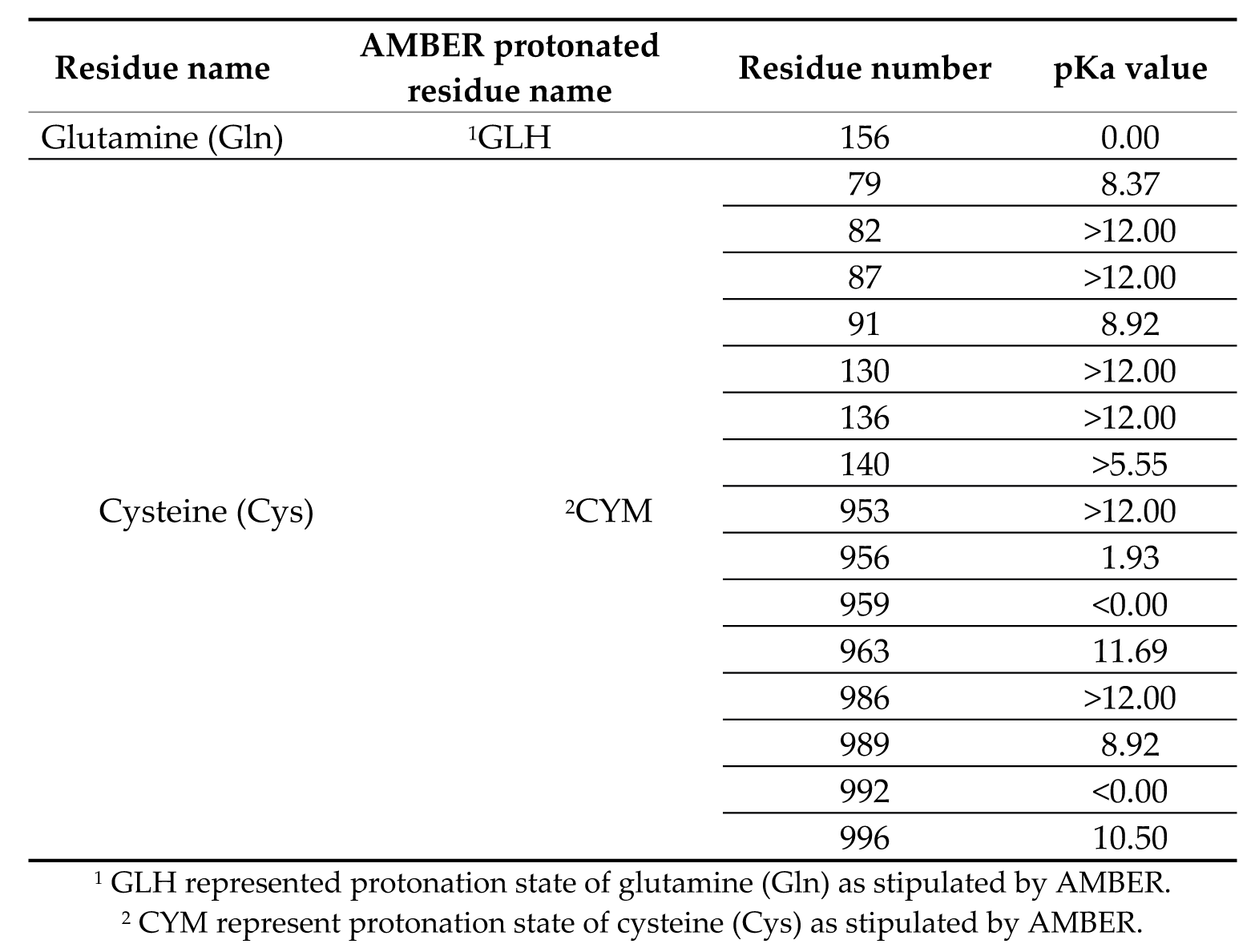
The protonation states and their pKa values of metal coordinating residues in human DPD protein model.

AMBER force field parameters of the Fe^2+^_4_S^2-^_4_ clusters in the human DPD protein were calculated using two approaches: the original Seminario method (Model 1) and collation features Seminario approach in visual-force field derivation tool (VFFDT) (Model 2). In each chain, two distinct residue coordinating environments were identified. Cluster 1026 (4 × Fe^2+^, 4 × S^2-^, 3 × Cys and 1 × Gln) coordination was different from those of clusters 1027, 1028 and 1029 (4 × Fe^2+^, 4 × S^2-^ and 4 × Cys). The four Fe^2+^ (FE1, FE2, FE3, FE4) bonded to the four S^2-^ (S1, S2, S3, S4) ions to form internal coordinates. Whereas four cysteine bounded the four Fe^2+^ (FE1, FE2, FE3, FE4) via a sulfide link (Cys [SG]) to form external coordinates of cluster 1027, 1028 and 1029. However, cluster 1026 coordinated externally to the four Fe^2+^ (FE1, FE2, FE3, FE4) through three Cys [SG] and the oxygen atom of Glutamine (Gln [OE]). Since the two monomers were a mirror image of each other, the Fe^2+^_4_S^2-^_4_ clusters with same geometry orientation were given similar number with a different letter representing their respective chains: chain-A (1026-A, 1027-A, 1028A and 1029-A) and chain-B (1026-B, 1027-B, 1028B and 1029-B). The subset structures representing all the possible coordination environment for F^e2+^ centers in DPD protein were used for QM calculation. By this approach, the computational time and resources utilized was greatly reduced compared to if all the clusters were to be considered. QM values for Fe^2+^_4_S^2-^_4_ subset clusters (1026-A and 1027-A) (Figure 2A) were generated for the Model 1. Whereas parameters for single internal coordinates (S3 and FE3) and external coordinates (Cys and Fe^2+^) and (Gln and Fe^2+^) were derived for Model 2 (Figure S1).

**Figure 2.**
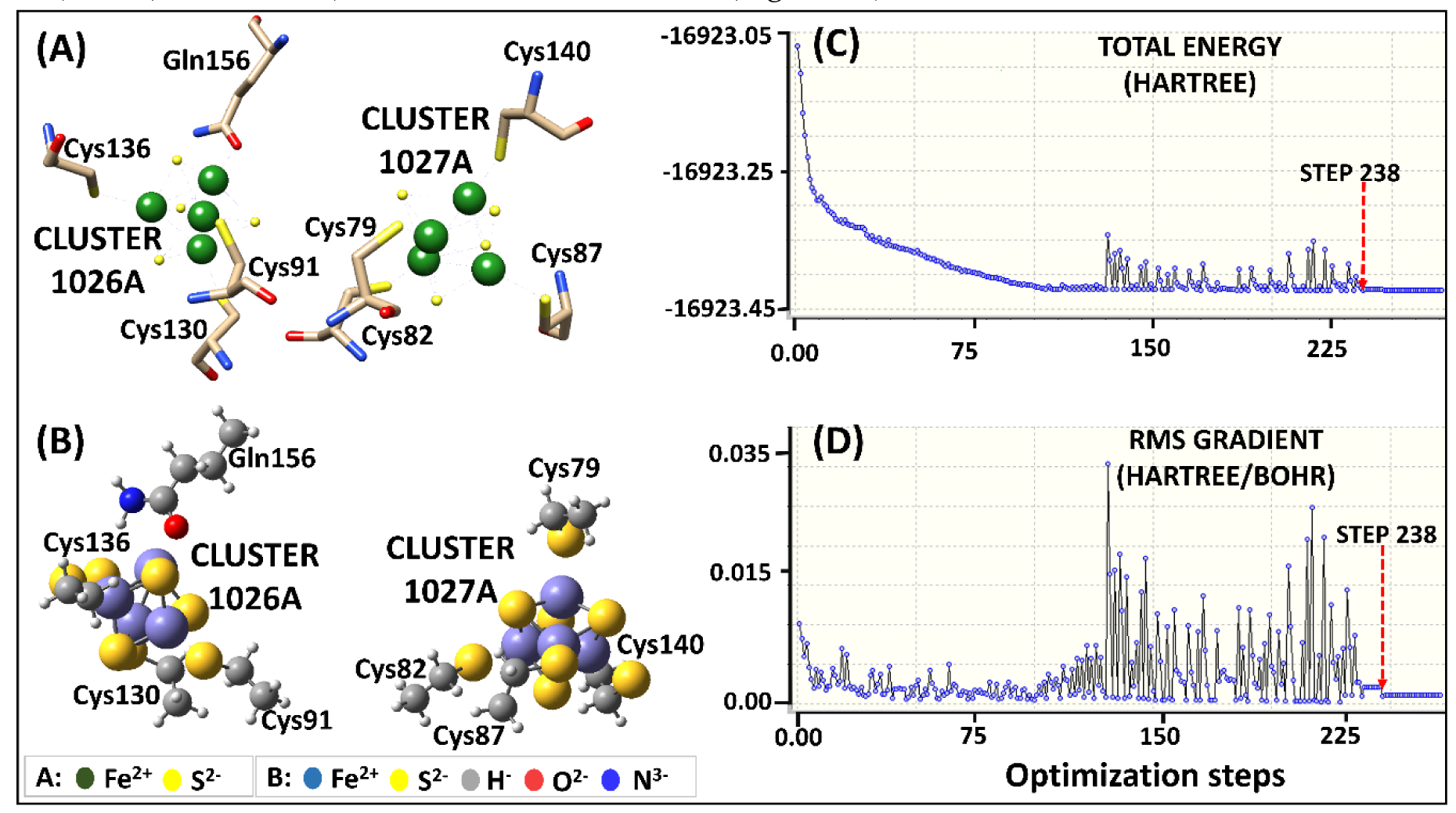
Human DPD Fe^2+^_4_S^2-^_4_clusters parameterization using original Seminario approach. **A**) 3D representation of Fe^2+^_4_S^2-^_4_ coordinating geometry. **B**) The optimized geometry of Model 1 human DPD subset at B3LYP/6-31G* level of theory. **C**) and **D**) Visualization of the energy potential using GaussView, showing the starting point of optimization at the lowest energy level (step 238).

#### 2.3.1. Geometry optimization

The subset structures for Model 1 attained the local minima at step number 238 initiating the optimization process (Figure 2C & D). During the optimization process, a significant energy variation between steps 120 and 230 was observed. The main cause of energy variation was due to the formation of a repulsive bond between Fe^2+^ and Fe^2+^ ions instead of Fe^2+^ and S^2-^ ions in cluster 1026. Nevertheless, the subset structures achieved correct optimization while maintaining their geometry as seen in Figure 2B.

The original Seminario method derived individual point value parameters for the subsets in Model 1 (Table S3). Contrastingly, the VFFDT (Model 2) approach/method generated average related parameters for internal bond length and angles, whereas the external parameters were averaged manually (Table S4). The equilibrium bond length and angle values obtained from QM (Models 1 and 2) showed some deviation from the crystal structure (Table 2, 3 and 4). These disparities might be due to deficient phase information on the x-ray structure since they give a static snapshot of the dynamic structure, contributing to spurious values [55]. Also, the disparity might have resulted from the lack of solvent effects and intermolecular interactions during the QM gas-phase optimization step [55, 56]. As expected, the average bond length and angle for Model 2 were within the range of those obtained from Model 1. Also, consistent with previous studies. In both models, the bond distances between Gln(OE)-Fe^2+^ were seemingly lower (Model 1: 1.92 Å and Model 2: 1.93 Å) (Table 2) as compared to the bond between Cys(S)-Fe^2+^. with force constant of 60.40 and 24.97 kcal mol^−1^Å^−2^, respectively. The short bond length might be attributed to the comparatively smaller atom radius of oxygen in Gln to that of sulfur in Cys [1, 2]. These values coincided with those obtained from previous related studies concerning Fe^2+^ and Cys [36, 37, 39, 57, 58]. However, there is limited literature in Fe^2+^ and Gln force field interactions which has been sufficiently addressed in this study.

**Table 2.**
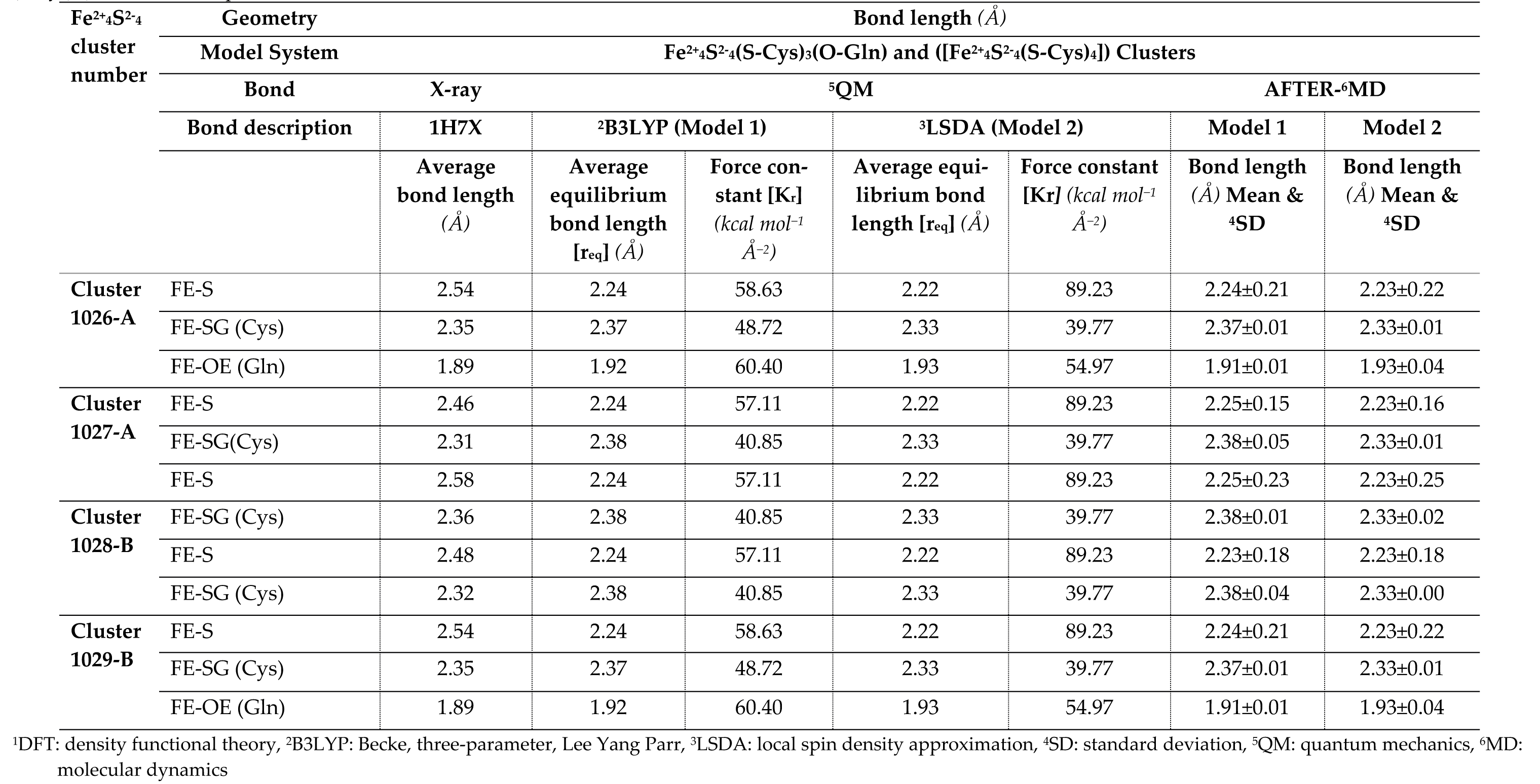
Comparison of average bond length (Å) calculated with X-ray, ^1^DFT ^2^(B3LYP) and ^3^(LSDA) method for the molecular cluster model ([Fe^2+^_4_S^2-^_4_ (S-Cys)_3_(S-Gln)]) and ([Fe^2+^_4_S^2-^_4_ (S-Cys)_4_]) of Native DPD protein.

**Table 3.** Comparison of average internal angles (^0^) calculated with X-ray, ^1^DFT ^2^(B3LYP) and ^3^(LSDA) method for the molecular cluster model ([Fe^2+^_4_S^2-^_4_(S-Cys)_3_(S-Gln)]) and ([Fe^2+^_4_S^2-^_4_(S-Cys)_4_]) of Native DPD protein

| Fe<sup>2+</sup><sub>4</sub>S<sup>2-</sup><sub>4</sub><br>cluster<br>number | Geometry |  |  |  | Angle (°) |  |  |  |
| --- | --- | --- | --- | --- | --- | --- | --- | --- |
|  | Model System Fe<sup>2+</sup><sub>4</sub>S<sup>2-</sup><sub>4</sub>(S-Cys)<sub>3</sub>(O-Gln) and ([Fe<sup>2+</sup><sub>4</sub>S<sup>2-</sup><sub>4</sub>(S-Cys)<sub>4</sub>]) Clusters |  |  |  |  |  |  |  |
|  | Angle | X-ray |  | <sup>5</sup>QM |  | AFTER-<sup>6</sup>MD |  |  |
|  | Angle description | 1H7X | <sup>2</sup>B3LYP (Model 1) |  | <sup>3</sup>LSDA (Model 2) |  | Model 1 | Model 2 |
|  |  | Average an-<br>gle (°) | Average<br>equilibrium<br>Angle [Θ<sub>eq</sub>]<br>(°) | Force con-<br>stant [K<sub>Θ</sub>]<br>(*kcal*<br>*mol<sup>-1</sup>rad<sup>-2</sup>*) | Average equi-<br>librium<br>angle[Θ<sub>eq</sub>](°) | Force constant<br>[K<sub>Θ</sub>]<br>(*kcal mol<sup>-1</sup>rad<sup>-2</sup>*) | Angle (°)<br>Mean & <sup>4</sup>SD | Angle (°)<br>Mean & <sup>4</sup>SD |
| Cluster<br>1026-A | FE-S-FE | 67.98 | 67.32 | 52.64 | 66.28 | 26.86 | 62.91±3.59 | 68.10±0.08 |
|  | S-FE-S | 106.03 | 108.50 | 39.12 | 109.21 | 39.52 | 109.25±2.28 | 106.99±0.68 |
| Cluster<br>1027-A | FE-S-FE | 68.39 | 67.61 | 49.30 | 66.28 | 26.86 | 64.55±2.72 | 68.24±0.11 |
|  | S-FE-S | 107.21 | 108.14 | 40.39 | 109.21 | 39.52 | 110.0±1.98 | 108.07±0.61 |
| Cluster<br>1028-B | FE-S-FE | 68.22 | 67.61 | 49.30 | 66.28 | 26.86 | 66.13±1.48 | 68.30±0.06 |
|  | S-FE-S | 106.51 | 108.14 | 40.39 | 109.21 | 39.52 | 107.02±0.36 | 106.97±0.33 |
| Cluster<br>1029-B | FE-S-FE | 67.97 | 67.61 | 49.30 | 66.28 | 26.86 | 65.15±1.99 | 67.48±0.35 |
|  | S-FE-S | 107.62 | 108.14 | 40.39 | 109.21 | 39.52 | 106.74±0.62 | 107.30±0.23 |

$^1$ DFT: density functional theory, $^2$ B3LYP: Becke, three-parameter, Lee Yang Parr, $^3$ LSDA: local spin density approximation, $^4$ SD: standard deviation, $^5$ QM: quantum mechanics, $^6$ MD: molecular dynamics

**Table 4.**
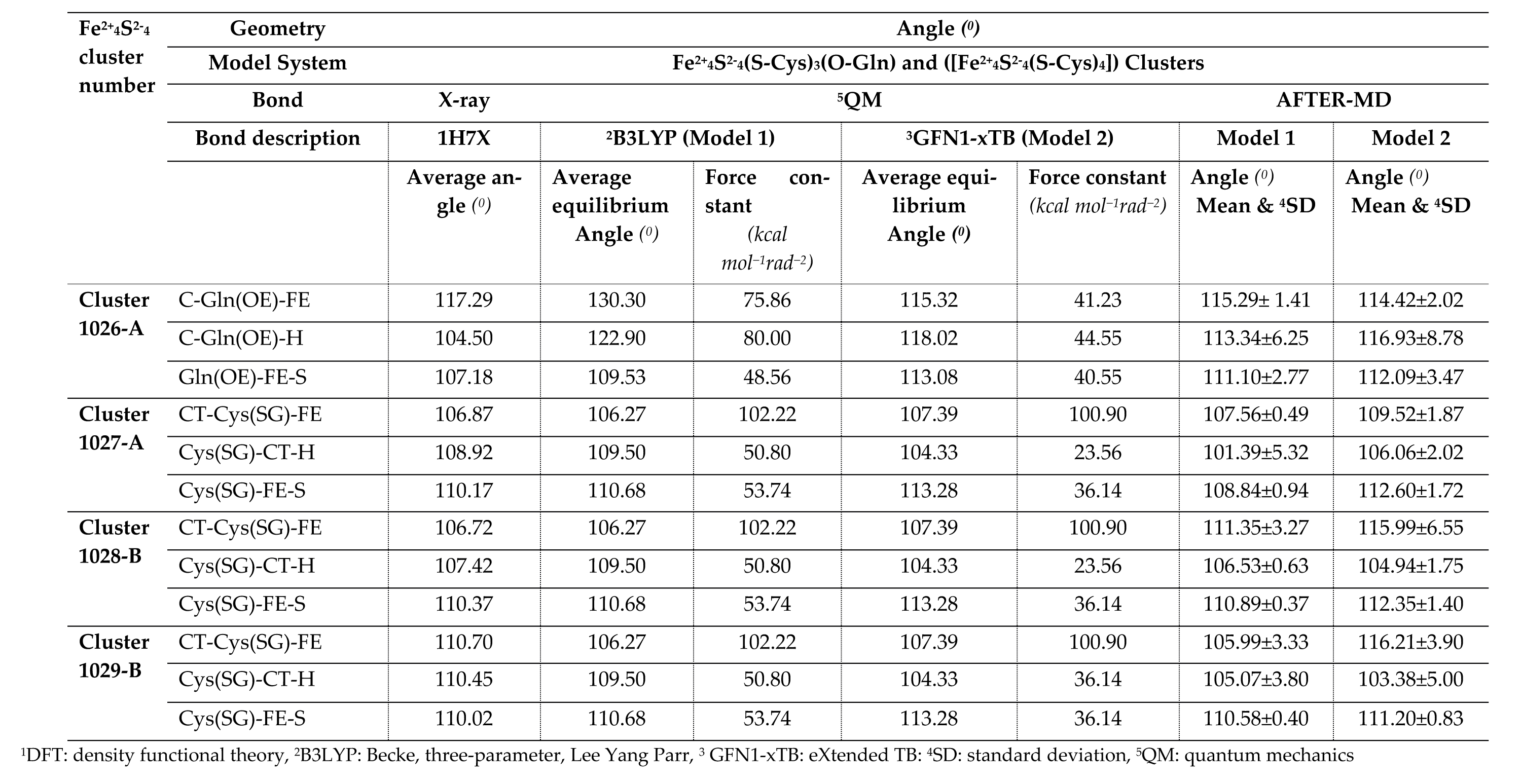
Comparison of average external angles (^0^) calculated with X-ray, ^1^DFT ^2^(B3LYP) and ^3^(GFN1-xTB) method for the molecular cluster model ([Fe^2+^_4_S^2-^_4_(S-Cys)_3_(S-Gln)]) and ([Fe^2+^_4_S^2-^_4_(S-Cys)_4_]) of Native DPD protein.

| Fe<sup>2+</sup><sub>4</sub>S<sup>2-</sup><sub>4</sub><br>cluster<br>number | Geometry | Angle ($^{\circ}$) | | | | | | |
| --- | --- | --- | --- | --- | --- | --- | --- | --- |
|  | Model System | Fe<sup>2+</sup><sub>4</sub>S<sup>2-</sup><sub>4</sub>(S-Cys)<sub>3</sub>(O-Gln) and ([Fe<sup>2+</sup><sub>4</sub>S<sup>2-</sup><sub>4</sub>(S-Cys)<sub>4</sub>]) Clusters |  |  |  |  |  |  |
|  | Bond | X-ray | <sup>5</sup>QM |  |  |  | AFTER-MD |  |
|  | Bond description | 1H7X | <sup>2</sup>B3LYP (Model 1) |  | <sup>3</sup>GFN1-xTB (Model 2) |  | Model 1 | Model 2 |
| | | Average an-<br>gle ($^{\circ}$) | Average<br>equilibrium<br>Angle ($^{\circ}$) | Force con-<br>stant<br>(*kcal*<br>*mol<sup>-1</sup>rad<sup>-2</sup>*) | Average equi-<br>librium<br>Angle ($^{\circ}$) | Force constant<br>(*kcal mol<sup>-1</sup>rad<sup>-2</sup>*) | Angle ($^{\circ}$)<br>Mean & <sup>4</sup>SD | Angle ($^{\circ}$)<br>Mean & <sup>4</sup>SD |
| **Cluster**<br>**1026-A** | C-Gln(OE)-FE | 117.29 | 130.30 | 75.86 | 115.32 | 41.23 | 115.29± 1.41 | 114.42±2.02 |
|  | C-Gln(OE)-H | 104.50 | 122.90 | 80.00 | 118.02 | 44.55 | 113.34±6.25 | 116.93±8.78 |
|  | Gln(OE)-FE-S | 107.18 | 109.53 | 48.56 | 113.08 | 40.55 | 111.10±2.77 | 112.09±3.47 |
| **Cluster**<br>**1027-A** | CT-Cys(SG)-FE | 106.87 | 106.27 | 102.22 | 107.39 | 100.90 | 107.56±0.49 | 109.52±1.87 |
|  | Cys(SG)-CT-H | 108.92 | 109.50 | 50.80 | 104.33 | 23.56 | 101.39±5.32 | 106.06±2.02 |
|  | Cys(SG)-FE-S | 110.17 | 110.68 | 53.74 | 113.28 | 36.14 | 108.84±0.94 | 112.60±1.72 |
| **Cluster**<br>**1028-B** | CT-Cys(SG)-FE | 106.72 | 106.27 | 102.22 | 107.39 | 100.90 | 111.35±3.27 | 115.99±6.55 |
|  | Cys(SG)-CT-H | 107.42 | 109.50 | 50.80 | 104.33 | 23.56 | 106.53±0.63 | 104.94±1.75 |
|  | Cys(SG)-FE-S | 110.37 | 110.68 | 53.74 | 113.28 | 36.14 | 110.89±0.37 | 112.35±1.40 |
| **Cluster**<br>**1029-B** | CT-Cys(SG)-FE | 110.70 | 106.27 | 102.22 | 107.39 | 100.90 | 105.99±3.33 | 116.21±3.90 |
|  | Cys(SG)-CT-H | 110.45 | 109.50 | 50.80 | 104.33 | 36.14 | 105.07±3.80 | 103.38±5.00 |
|  | Cys(SG)-FE-S | 110.02 | 110.68 | 53.74 | 113.28 | 36.14 | 110.58±0.40 | 111.20±0.83 |

<sup>1</sup>DFT: density functional theory, <sup>2</sup>B3LYP: Becke, three-parameter, Lee Yang Parr, <sup>3</sup> GFN1-xTB: eXtended TB; <sup>4</sup>SD: standard deviation, <sup>5</sup>QM: quantum mechanics

Despite the slight differences, the value of force constant from both systems (Model 1 and 2) were within the same range, and consistent with those obtained from previous studies [36, 59]. Commonly, force field parameter values of a model conducted under different systems are not exact but fall within expected range [36, 37, 56]. In generating new parameters, the state of structure geometry optimization is thought to be a contributing factor to the varied observation [60]. Previous findings [61] ascribed the discrepancies to the different methods used in obtaining the force constant and the opposite way of how the connectivity’s were defined. Most importantly, the derived values showed that both models maintained the subsets structures geometry following optimization step.

#### 2.2.2. RESP charges

Partial atomic charge calculations were derived for each atom interacting with the Fe^2+^ center for the optimized subset structures. Figure S2 and Table S5 illustrate differences in the WT DPD atomic charge distribution in the oxidized subsets. The RESP method derived these charges by fitting the molecular electrostatic potential obtained from the QM calculation based on the atom-centered point charge model. In their oxidized state, atoms within the DPD Fe^2+^ (S^2-^, Gln and Cys) center exhibited varied atomic charges due to the large electrostatic environment around the protein’s metal sphere. Such variations are known to influence charge transfer at the redox center bringing stability around the coordinating sphere of metalloproteins [62]. As such, they are vital components in the achievement of accurate inter- and intra-molecular potential electrostatic interaction [55].

#### 2.2.3. Inferring the generated QM force fields parameters to the corresponding identical clusters

The newly generated Fe^2+^ force field parameters for subsets 1026-A and 1027-A (Table S3) were inferred to the remaining Model 1 DPD clusters corresponding to their geometry mentioned earlier. Similarly, the generated internal and external parameters (Table S4) for Model 2 were also inferred to the corresponding clusters accordingly. At the end, each model featured a holo and a holo-drug (5-FU cancer drug) protein complex totaling to 64 internal (Fe-S) and 32 external (30 Cys-Fe; 2 Gln-Fe) parameter calculations of the DPD (Fe^2+^_4_S^2-^_4_) clusters. In terms of energy profile and range of force constants for Model 1 and 2, there were no significant differences observed in terms of DPD Fe^2+^ ion coordination to Cys, Gln residues, and S^2-^ ions. Table 2, 3, and 4 show a summary equilibrium bond length, angle, and related force constants with detailed information available in supporting information (Tables S6 and S7). Dihedral-related force constants were derived manually from the respective structures (Table S8).

### 2.3. Genereted force field parameters validated using MD simulations

#### 2.3.1. Analysis of protein stability and flexibility through RMSD, RMSF, Rg

Accurate parameters are necessary for maintaining the coordinating geometry of a metal center in metalloproteins [37]. Therefore, to evaluate the accuracy and reliability of the derived parameters (Model 1 and 2), all atom MD simulations (150 ns) for holo system and holo-drug complexes were performed. The derived parameters were validated by assessing the root mean square deviation (RMSD) (Figure 3A), the radius of gyration (Rg) (Figure 3B) and root mean square fluctuation (RMSF) (Figure 3C). Simulations of both models for holo and holo-ligand complexes showed minimal deviation from their initial structure, and were maintained across the simulation process (Figure 3A). Model 1 systems (holo and holo-drug) displayed multimodal RMSD density distribution, implying they sampled various local minima whereas, each of Model 2 proteins attained a single local minimum (unimodal distribution). The Rg (Figure 3B) revealed that the compactness of the various protein models remained the same during dynamics. However, differences were observed between the holo and holo-drug bound proteins. The ligand-bound protein was seen to generally have a higher Rg than the non-ligand bound protein in both model systems. This may be attributed to the presence of drug. Proteins from both models exhibited similar RMSF profiles (Figure 3C). However, the ligand-bound proteins appeared slightly flexible than the non-ligand bound ones. As expected, the loop regions which constitute ∼43 % of the entire protein structure, including the active-site loop (residues 675–679), were the most flexible regions while the metal site residues displayed minimal fluctuation (Figure S3). Visualization of the different trajectories through Visual Molecular Dynamics (VMD) [63] verified a high conformational change of the loop areas, while the protein central core containing Fe^2+^ clusters had vibrational like movements.

**Figure 3.**
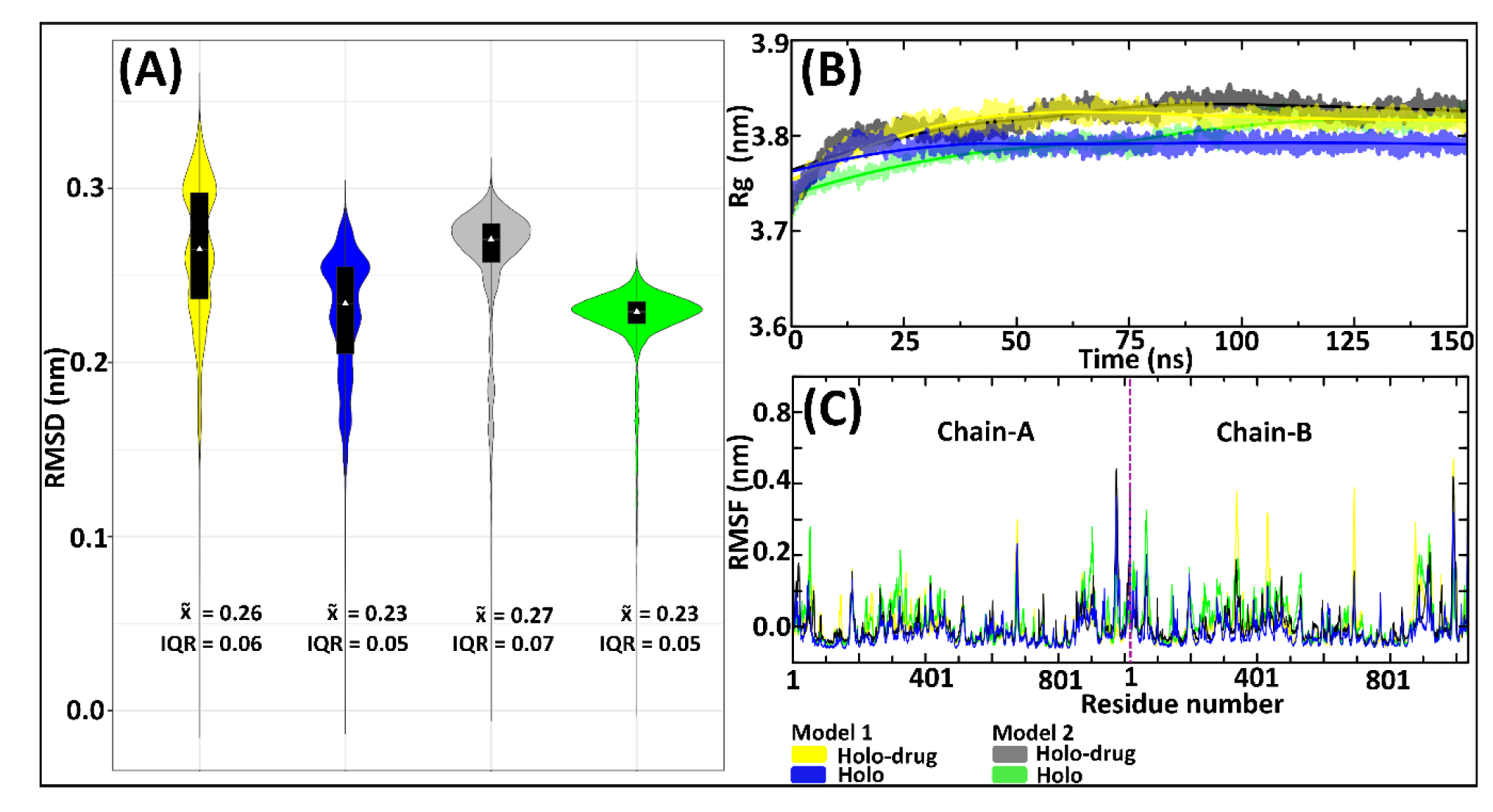
Evaluation of the DPD protein stability by RMSD, Rg and RMSF of the 150 ns MD simulations for Model 1 and Model 2, drug (5-FU) and non-drug bound systems. Model 1 is represented by yellow (drug) and blue (non-drug) bound systems. Model 2: grey (drug) and green (non-drug) bound systems. **A**) Bachbone RMSDs showing an overlay of boxplots on kernel density estimation graphs. Box-plots show the median, upper and lower quartiles. The non-parametric kernel density plots are represented as median and interquartile range (IQR). **B**) Rg line graphs showing the compactness of all system **C**) RMSF showing illustrating the fluctuation of residues with the puple line separating chain-A and chain-B.

The profiles of RMSDs (Figure 3A) exhibited higher variation in conformational changes across all systems. These variations were more apparent in the Model 1 system’s proteins compared to the Model 2 system. Considering the similarity of protein behavior with drug binding, it is apparent that both Model show similar atomic tendencies in the drug and non-drug bound systems. The disparities arising from conformational changes were because of the slight differences in the approaches used in the Models preparations. For instance, fixed bond parameters were assigned between Fe-S, Fe-Fe and the connecting residues (Fe-Cys or Fe-Gln) of Model 2 based on averages of crystallographic structure (Table S2), where- as Model 1 parameters were attained from single point atom calculation of the crystallographic structure. The RMSF values of both the holo and holo-drug bound complex demonstrated region of higher flexibility between residues in all models (Figure 3C).

Proteins are dynamic entities and as such they undergo conformational changes as part of their functionality. Elucidating these changes is necessary in understanding how their functionality is maintained [64]. Hence, we evaluated the conformational variations sampled by each system during simulation by plotting the free energy of each system snapshot as a function of RMSD and Rg using the Boltzmann constant (Figure 4). In both models, free energy investigations revealed similar tendencies as the kernel density map in all the systems. Both holo and holo-drug bound protein populated three main conformations in Model 1. However, the holo bound protein attained three energy minima at 0.18, 0.20 and 0.25 nm while the drug-bound protein energy minima were attained later at (0.22, 0.25 and 0.35 nm). On the other hand, Model 2 equilibrated at single energy minima for both the drug (0.28 nm) and holo (0.22 nm) bound complexes. Model 1 proteins attempted severally to find a high probability region that guaranteed more stability thermodynamically for its conformational state than Model 2. Yet, upon drug binding the conformation entropy was increased in both model which destabilized the transitional state and simultaneously slowed down protein equilibration. Visualization of the trajectories in VMD to establish the cause of trimodal ensemble showed alternating movements in the loop regions including the C-terminal, N-terminal and active-site loop areas. More importantly, the Fe^2+^_4_S^2-^_4_ clusters geometry was maintained during the simulation (Figure S4).

**Figure 4.**
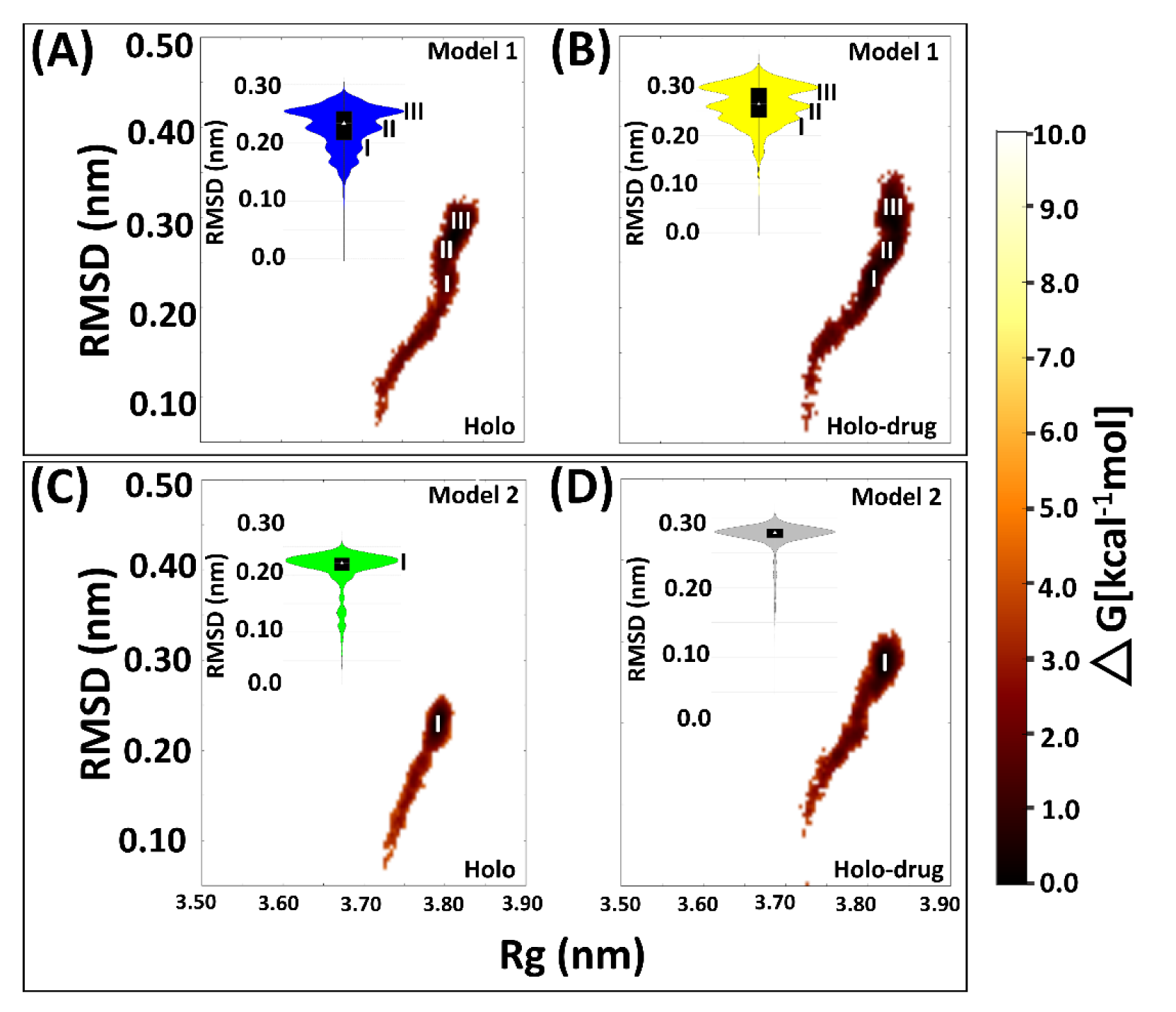
Free energy landscape of the four systems snapshots plotted as RMSD and Rg values derived from the Boltzmann constant in relation to the kernel density plot. **A**) Drug bound system (Model 1) showing three (I, II and III) major conformational changes sustained by the protein during MD simulation. **B**) Holo bound system (Model 1) showing three (I, II and III) major conformational changes. **C**) Drug bound system (Model 2) showing one (I) major conformational change **D**) Holo bound system (Model 1) protein exhibiting one major conformational change during 150 ns simulation. Model 1: 5-FU drug bound protein (yellow); holo bound protein (blue). Model 2: Holo-drug bound protein (grey) and holo bound protein (green). Free energy landscape (maroon).

#### 2.3.2. Fe^2+^_4_S^2-^_4_ clusters exhibited stability during MD simulations

Assessment of inter- or intra-molecular distances between groups of interest can be used to investigate stability changes during MD simulations [65]. In this study, distances between the center of mass (COM) of; 1) the entire DPD protein and each of the eight Fe^2+^_4_S^2-^_4_ clusters (Figure 5A); 2) each chain and the four Fe^2+^_4_S^2-^_4_ clusters therein (Figure 5B); and 3) the active site of each chain and its Fe^2+^_4_S^2-^_4_ clusters were evaluated (Figure 5C) for each model (Model 1 and 2: holo and holo-drug). From these calculations, the overall stability of the key components involved in the electron transfer process was evaluated. Generally, the inter-COM distances between the various groups in both models were nearly similar (Figure 5 A-C). Moreover, data was distributed with less standard deviation (uni-modal distribution) as seen from most kernel density plots, suggesting the distances within metal clusters remained in the same range across the 150 ns simulation and maintained stability within the metal clusters. Thus, the two methods can reliably be used to achieve accurate parameters for other metalloproteins.

**Figure 5.**
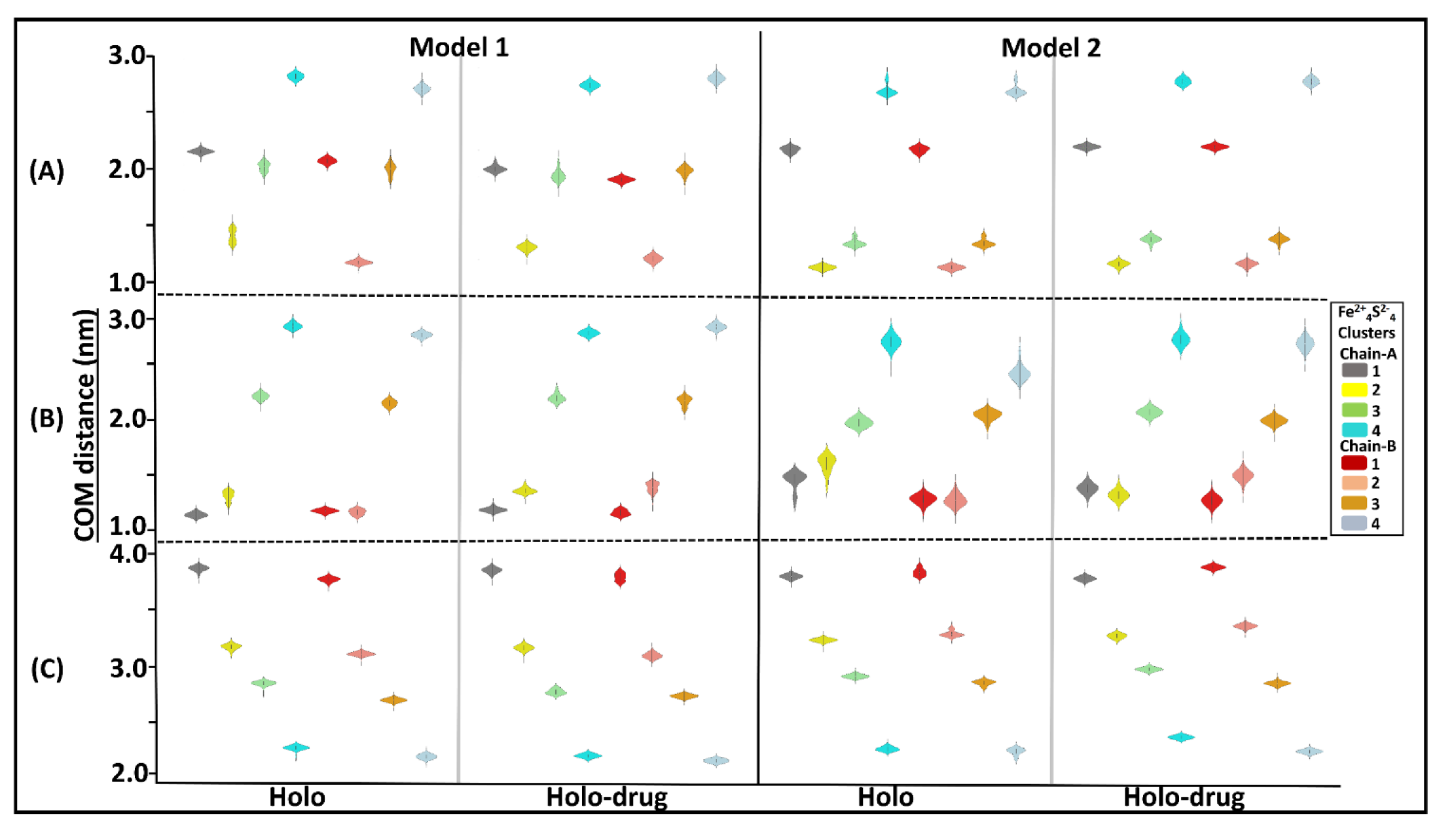
Kernel density showing distribution of COM distances across different Fe^2+^_4_S^2-^_4_ clusters and the protein, the chains (A and B) and the active sites. Grey represents cluster 1 (1026A), yellow: cluster 2 (1027A), green: cluster 3 (1028A), cyan: cluster 4 (1029A), red: cluster 5(1026B), salmon: cluster 6 (1027B) orange: cluster 7 (1028B) and light blue: cluster 8 (1027B). The interquartile range and the median are shown inside the kernel density plot. **A)** The distribution of COM distance between different clusters and the proteins **B)** chain-A (cluster 1, 2, 3 and 4) chain-B (cluster 5, 6, 7 and 8) and **C)** active-site. Generally, a uni-modal distribution is seen across all clusters in both models. The distance between the Fe^2+^ cluster and backbone of the protein remained within the same range during dynamics. Clusters compactness is an indication of the system stability. Respective clusters are colored accordingly.

In addition to the group inter-COM distance calculations, the distances between the Fe^2+^ centers and the coordinating residues were also determined for the holo-drug complexes in both models (Figure 6). By this approach, the integrity of the coordinating geometry can be accessed during simulations. From the results, a high bond length consistency was observed within all Fe^2+^_4_S^2-^_4_ centers, an indication that the derived parameters were accurately describing the cluster geometries. Furthermore, the obtained bond lengths were in agreement with those reported previously [34, 66]. The maintenance of the bond distances signified that the desired functionality and stability had not been jeopardized given that it is dependent on the protein environment [36]. Notably, Zheng et al. protocol on the evaluation of metal-binding structure confirmed that the coordinating tetrahedral geometry of Fe^2+^_4_S^2-^_4_ clusters was maintained during the entire simulation runs. Although our calculations agreed with previous findings [36, 39, 67], it is worth noting, to the best of the authors’ knowledge, none of the studies featured the force field parameters for glutamine interaction with a single or multiple Fe^2+^_4_S^2-^_4_ clusters in a single protein.

**Figure 6.**
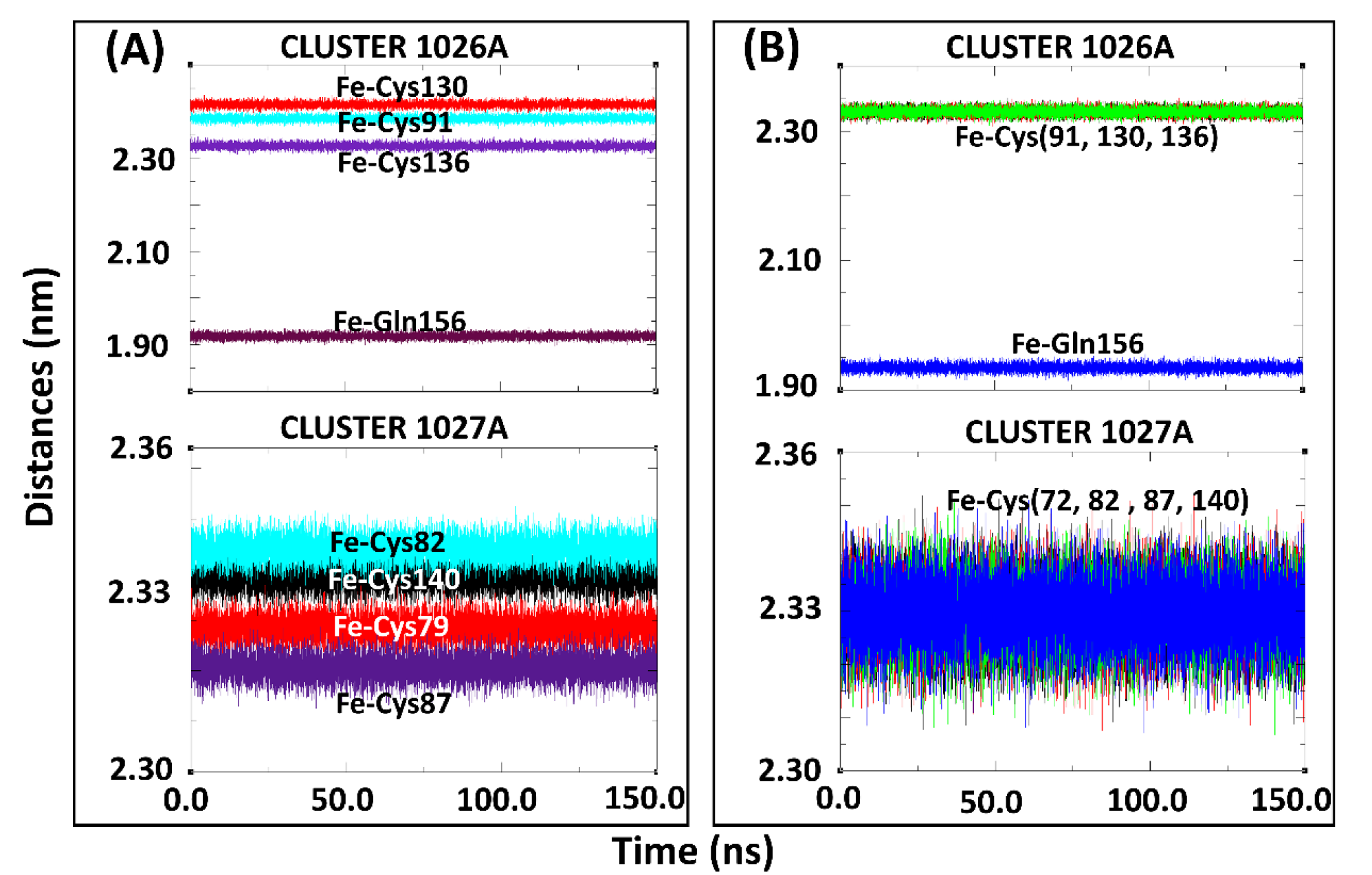
Coordination of subset structure residues to the Fe^2+^ centers for the holo-drug bound systems of Model 1 and Model 2 during 150 ns MD simulations **A**) Cluster 1026A and 1027A of Model 1 the hololigand system **B)** Cluster 1026A and 1027A of Model 2 the holo-drug bound system. The coordinating distances between the Fe^2+^ and the connecting residue is seen to be maintained throughout the simulation in both models.

#### 2.3.3 Validation of derived parameters in IH7X crystal structure

The derived Fe^2+^_4_S^2-^_4_ parameters coordinated uniquely to Cys and Glu residues were inferred to the template structure (PDB ID: 1H7X) for additional validation purposes. As in the modeled human structure, the four Fe^2+^_4_S^2-^_4_ clusters in each chain of the template also maintained the correct geometry as shown in Figure S5.

### 2.4. Essential motions of protein in phase space

Proteins are dynamic entities whose molecular motions are associated with many biological functions including redox reactions. Collective coordinates derived from atomic fluctuation principal component analysis (PCA) are widely used to predict a low-dimensional subspace in which essential protein motion is expected to occur [68]. These molecular motions are critical in biological function. Therefore, PCA were calculated to investigate the 3D conformational study and internal dynamic of holo and holo-drug complexes of both models (Model 1 and Model 2). The first (PC1) and the second (PC2) principal components captured the dominant protein motions of all atom 150 ns MD simulation (Figure 7). Both holo structures (Model 1 and Model 2) showed a U-shaped time evolution from from unfolded state (yellow) emerging from simple Brownian motion and ending in native state (dark blue) over 150 ns. Strikingly, the projection of holo-drug complexes from both Models (1 and 2) adopted a V-shaped time evolution space emerging from unfolded state (yellow) and ending in native state (dark blue). Model 1 and Model 2 holo structures accounted for 44.95 % of the global total structural variances. The holo-drug complexes reported 48.95 % and 36.5 % of global total variance for Model 1 and Model 2, respectively. In overall, the holo-drug complexes (Model 1 and Model 2) exhibited altered conformational evolution over time in-comparison to their respective holo structure, suggesting the newly force field parameters derived from both models maintains protein function.

**Figure 7:**
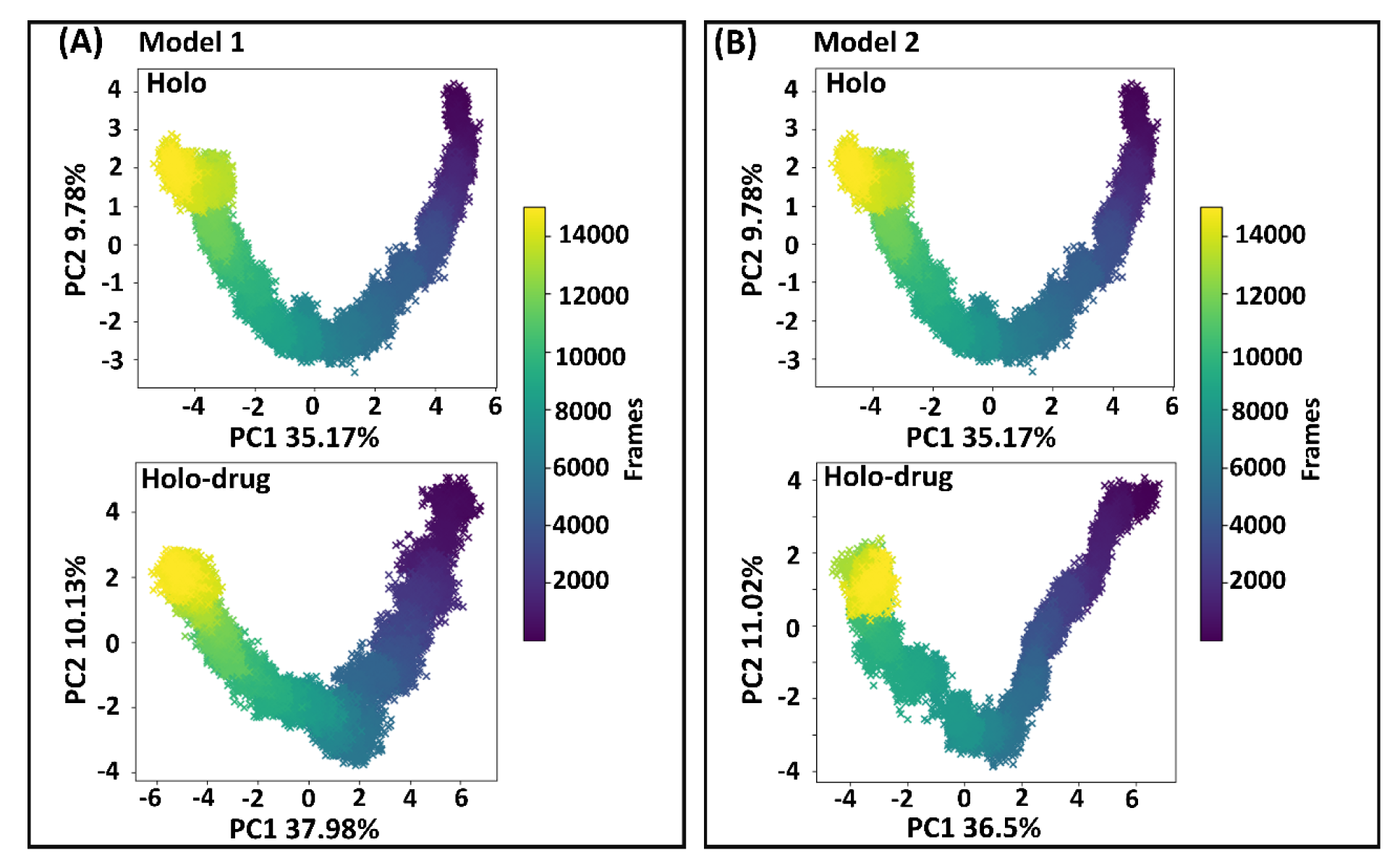
Principal component analysis. First and second principal component analysis (PC1 and PC2) of human DPD wild-type extracted from essential dynamics. The time evolution of transitional from unfolded state of the DPD protein (yellow) emerging from simple brownian motion and ending in the native state (dark blue) over 150 ns. A) The first two PCs of Model 1 accounting for 44.95 % and 48.95 % of the total structural variance of the holo and holo-drug complexes, respectively. B) The first two PCs of Model 2 accounting for 44.95 % and 47.52 % of the total structural variance of the holo and holo-drug complexes, respectively.

**Figure 8.**
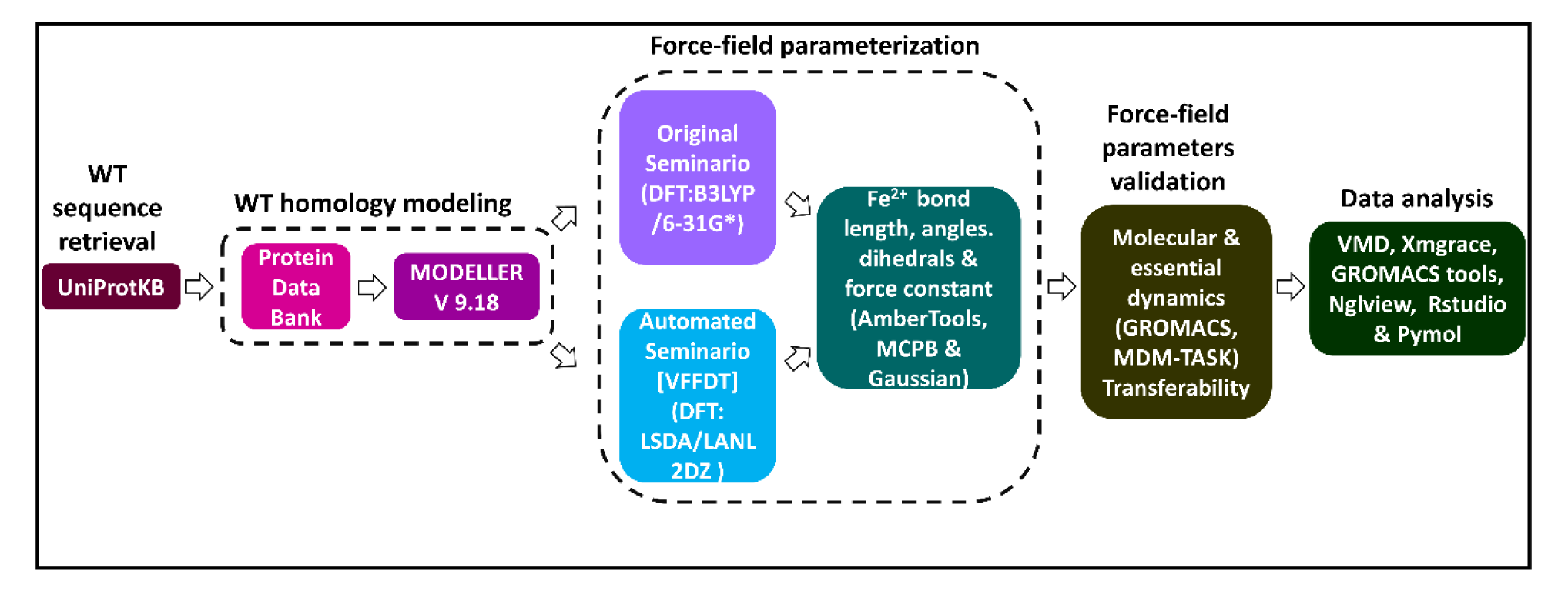
A flow diagram illustrating a summary of methods and tools used in the generation and validation of Fe^2+^_4_S^2-^_4_ force field parameters for human DPD protein model. Two approaches (original Seminario and automated Seminario) were used to determine bond lengths, angles and dihedrals around the Fe^2+^_4_S^2-^_4_ centers. The reliability of the generated parameters in describing the coordination geometry of the Fe^2+^_4_S^2-^_4_ centers was evaluated using all atom MD simulations.

## 3. Materials and Methods

### 3.1. Software

AMBER and AmberTools17, University of California, San Francisco, USA; AutoDock4.2 software, The Scripps Research Institute, San Deigo, USA; Discovery Studio v4.5, Dassault Systems BIOVIA, San Deigo, USA; GaussView 5.0.9, Carnegie Mellon University Gaussian, Conneticut, USA; GROMACS v5.1.5., University of Groningen, Uppsala Sweden; Rstudio v1.1.456, R Core Team; PyMOL Molecular Graphics System; v1.8.2.3 Schrödinger, New York, USA.

### 3.2. Homology modeling of native DPD protein

Due to the absence of the human DPD protein crystal structural information in the Protein Data Bank (PDB) database [10], homology modeling approach was used to obtain a complete 3D structure using MODELLER v9.15 [61]. The technique has become indispensable in obtaining 3D models structures of proteins with unknown structures and their assemblies by satisfying spatial constraints based on similar proteins with known structural information [69]. The restraints are derived automatically from associated structures and their alignment with the target sequence. The input consists of an alignment of a sequence to be modeled with a template protein which structure has been resolved, and a script file (Table S9). At first, the target sequence (human DPD enzyme - UniProt accession: Q12882) was obtained from the Universal Protein Resources (ref). Both HHPred [70] and PRIMO [71], were used to identify a suitable template for modelling the human DPD protein. From the potential templates listed by the two webservers, PDB 1H7X, a DPD crystal structure from pig with a resolution 2.01 Å was identified as the top structural template having sequence identity of 93% [1,2]. A *pir* alignment file was prepared between the Uniprot (UniProt accession: Q12882) target sequence and that of template using MUltiple Sequence Comparison by Log-Expectation (MUSCLE). Therefore, the template PDB ID: 1H7X. In MODELLER v9.14 [44], a total of 100 human DPD holo models were generated under ‘very-slow’ refinement level guided by the selected template. The resulting models devoid of both drug (5-FU and cofactors) were ranked based on their lowest normalized discrete optimized protein energy (z-DOPE) score [46], and the top three best models selected for further modeling. To incorporate the non-protein structural information, each of the selected models was separately superimposed onto the template in Discovery Studio 4.5 [45] and all non-protein information copied. The coordinates for cofactors and the drug were then transferred directly to the modeled structures. Further quality assessment of the resulting complexes was performed using VERIFY3D [48], PROCHECK [51], QMEAN [49] and ProSA [50]. The best model showing consistently high-quality score across the different validation programs was chosen for further studies.

### 3.3. Protonation of titrarable residues

To account for the correct protonation states of the system, all DPD titratable residues were protonated at pH 7.5 [1], system salinity of 0.5 M, internal and external default dielectric constants of 80 and 10 respectively in the H++ web server [53]. System coordinates (crd) and topology (top) files were used to build protonated protein structure files. A visual inspection of all titratable residues was performed and wrongly protonation corrected using Schrödinger Maestro version 11.8.

### 3.4. New force field parameters generation

Prior to the parameter generation process, the residue coordination present in chain-A and chain-B Fe^2+^_4_S^2-^_4_ centers were evaluated to identify representative subsets. Two unique coordination subset arrangements *viz*. 1026-A (4 × Fe^2+^, 4 × S^2-^, 3 × Cys and 1 × Gln) and 1027-B (4 × Fe^2+^, 4 × S^2-^ and 4 × Cys) were identified. The two subsets (1026-A and 1027-B) represented the coordinating geometry of all Fe^2+^_4_S^2-^_4_ clusters in the protein. Subsequently, force field parameters describing the coordinating interactions in these unique centers were determined via two approaches. Firstly, the original Seminario method (Model 1) was implemented using the bonded model approach in AmberTools16 [43] and python-based metal center parameter builder (MCPB) [42]. Gaussian 09 [71, 72] input files (.*com*) of the protonated protein incorporating the subsets structures (1026-A and 1027-B) were prepared. Thereafter, their geometries were optimized utilizing the hybrid DFT method at Becke three-parameter hybrid exchange and Lee Yang Parr (B3LYP) correlation function level of theory. This process utilized double split-valence with polarization [6-32G(d)] basis set [72, 73] (Table S1). Sub-matrices of Cartesian Hessian matrix were used in the derivation of the metal geometry force field parameters [41]. Bond and angle force constants were obtained via fitting to harmonic potentials. The potential energy of the relative position for each atom in the system was determined by Amber force field parameters calculated from the equation (1) below:

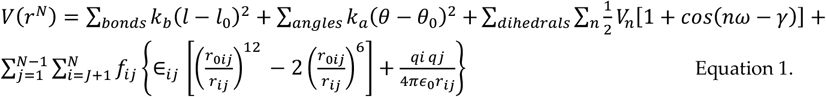

Where the bond lengths, angles values, torsion values and the interatomic distances were obtained. The first and second term of the harmonic potential energy function relates to bond bending and bond stretching, respectively, whereas the torsion angles have been described by the third term. Lastly, the van de Waals forces and electrostatic interaction were given by the non-bonded energy function involving the Lennard Jones (12-6) potential and Coulomb potential, respectively [41, 74]. The optimized/minimized structure were then visualized in GaussView 5.0.9 [75] to confirm that the bonds in the centers were intact. The atomic charges of the optimized subset structures were then derived from electrostatic potential (ESP). However, ESP assigns unreasonably charged values to buried atoms, which impair their conformational transferability. Therefore, the Restrained Electrostatic Potential (RESP) fitting technique, which considers the Coulomb potential for the calculation of electrostatic interaction, was devised to address these issues. This method has been highly regarded and widely used in assigning partial charges to various molecules utilizing B3LYP/6-31G(d) gas phase [76]. Restrains in terms of penalty functions are applied on the buried atoms, leading to multiple possible charged values. Hence, the quality of fit to the QM ESP is not compromised [77]. Herein, a default Merz-Kollman Restrained Electrostatic Potential (RESP) radius of 2.8 Å was allocated to the metal centers. An additional approach (herein named as Model 2) using collation features Seminario: VFFDT program was used [43]. Analysis data was acquired following optimization of subsets Fe^2+^-S^2-^, Fe^2+^-Cys and Fe^2+^-Gln coordination; the calculations were performed using density functional theory (DFT) featuring the local spin density approximation approach with the Los Alamos double-zeta basis (LSDA/LANL2DZ) (Table S2) [78, 79]. This factored in the internal covalent bonds; note that the calculation was not successful at the B3LYP level of theory. The external non-covalent bond calculation was determined by eXtended TB (GFN1-xTB) [80]. Retrieval of the force field parameters for the entire molecule was done through Protocol menu item “FF” for the whole General Small Molecule”. Since the system in this study was symmetrical, the atom types were left identical to Fe or S. The AMBER force field parameters for Fe^2+^ metal center bond and angles were then generated automatically. Individual detailed statistics were derived but only the final values were utilized for further calculations. The obtained parameters were then inferred to the other clusters in the modeled structures as well as the template crystal structure (PDB ID: 1H7X) using the LEaP [81] program. This was based on the similarity of the clusters coordinating geometry. As such, cluster 1026-A was inferred to 1026-B, and those for 1027-A were inferred to 1027-B, 1028-A, 1028-B, 1029-A and 1029-B as they depict identical co-ordination geometry. In total, 2 × ([Fe^2+^_4_S^2-^_4_(S-Cys)_3_(S-Gln)]) and 6 × ([Fe^2+^_4_S^2-^_4_ (S-Cys)_4_]) clusters parameters were derived for each model. To our knowledge, no other 3D structure with metal centers, such as the human DPD coordinating environment, was available in the PDB. Therefore, the pig crystal structure was used to validate the reliability and accuracy of the newly generated force field parameters.

### 3.5. Force field parameters validation and analysis

To evaluate the reliability of generated parameters derived from the original and automated Seminario approaches, duplicate all-atom MD simulations were conducted using GROMACS 5.14 MD package [52]. For each model system (Model 1, Model 2, 1H7X crystal structure), the holo (protein with only cofactors) and holo-drug (5-FU) complexes were considered for simulation studies. At first, AMBER topologies for each system were generated by Leap modelling with the AMBER ff14SB force field to incorporate all the generated parameters [82]. The resulting system topologies were converted to GROMACS-compatible input files for the structure (gro) and the topology (top), with the correct atom types and charges using the AnteChamber Python Parser interface (ACPYPE) tool [83]. The infinite systems were then solvated in an octahedron box system using the simple point charge (SPCE216) water model [84], with a padding distance of 10 Å set between the protein surface and the box face. The net charge for all systems was subsequently neutralized by adding 0.15 M NaCl counter-ions [85]. The neutralized systems were then subjected to an energy minimization phase (without constraints) using the steepest descent integrator 0.01 nm and a maximum force tolerance of 1000 kJ mol^−1^nm^−1^ was attained. This was necessary to get rid of steric clashes that may have resulted during incorporation of parameters and water molecules. Subsequently, the systems were equilibrated to ensure that they attained the correct temperature and pressure using a two-step conical ensemble (each 100 ps). Firstly, the temperature was set at 300 K (NVT - number of particles, volume, and temperature) using modified Berendsen thermostat. This was followed pressure equilibration at 1 atm (NPT - number of particles, volume and temperature) using Parrinello–Rahman barostat algorithm [86]. The ensembles utilized the revised coulomb type for long range electrostatic interactions with a gap cut of 8.0 Å as described by particle mesh Ewald (PME) [87] method, and LINCS algorithm was used to constrain bonds in all atoms [88]. Finally, the production MD simulations of 150 ns were performed for all the systems at the Centre for High Performance Computing (CHPC) in Cape Town South Africa using 72 Linux CPU cores, with time integrations step of 2 fs. Coordinates were written to file every 10 ps.The obtained MD trajectories were stripped off all periodic boundary conditions (PBC) and fitted to the reference starting structure.

#### 3.5.1. Root mean square, root mean square fluctuation and Radius of gyration analysis

Global and local conformational behaviour of the replicate ensembles were determined using various GROMACS modules *viz. gmx rms, gmx rmsf, gmx gyrate, gmx distance* and analysed in RStudio [89]. These packages were used to analyze the root mean square deviation (RMSD), root mean square fluctuation (RMSF) the radius of gyration (Rg), and the inter-center of mass between groups of interest, respectively. The overall conformational changes per system were observed using Visual Molecular Dynamics (VMD) [63] to ensure that the derived parameters correctly maintained the geometry of the various Fe^2+^_4_S^2-^_4_ clusters.

#### 3.5.2. Principal Component analysis

Principal component analysis (PCA) was conducted in the MDM-TASK-web to investigate the time evolution of the protein’s conformational changes in MD trajectories [68, 90]. PCA is a linear transformation technique that extracts the most important element from a data set by using a covariance matrix built from atomic coordinates defining the protein’s accessible degree of freedom. The calculation of the coordinate covariance matrix for the Cα and Cβ atoms were implemented after RMS best-fit of the trajectories was applied to an average structure [68, 90]. Corresponding eigenvectors and eigenvalues were then obtained from diagonalized matrix. Protein coordinates were then projected using eigenvectors. PC1 verses PC2 plots were then derived from the normalized primary and secondary projections.

#### 3.5.3. Additional analytical approach

Molecular graphics were then prepared with PyMOL v1.8 [91], Anaconda 4.3.1 Jupyter Notebooks [92], and various open-source Python libraries, such as matplotlib [93], Seaborn, Pandas [94], NumPy [95], and NGLview [96].

To ascertain how accurate the generated force field parameters were, the average bond lengths and force constants from the derived parameters were compared to those of the x-ray structure. All statistical calculation was performed using *Welch t-test* in Rstudio v1.1. 456 [89] where p – value (< 0.05) was considered significant.

## 4. Conclusions

In addition to the nucleotide metabolizing function of the DPD metalloenzyme in humans, the dimeric protein also serves as an important anti-cancer drug target [4, 5, 6]. Deficiency or dysfunction of the enzyme as a result of mutations results to increased exposure to active fluoropyrimidines metabolites leading to severe toxicity effects. Computational approaches such as MD simulations have become integral components of elucidating protein function as well as the effects of mutations [4]. MD simulations allow the elucidation of conformational evolution of protein systems overtime during a reaction process [26, 31, 32]. MD simulations require appropriate mathematical functions and set of parameters collectively known as force fields which describe the protein energy as a function of its atomic coordinates. In cases where adequate parameters are lacking, especially those describing non-protein components in a system, additional descriptors are necessary. In this work, which forms a platform for future studies towards anticancer personalized medicine, we report new validated AMBER parameters that can be used to accurately describe the complex Fe^2+^_4_S^2-^_4_ clusters in the DPD protein and related systems. This was motivated by the absence of ready to use force field parameters enabling in silico studies on the DPD system. The development of combined QM/MM methods has provided the most effective accurate and theoretical description of the molecular system [72]. They enable a comprehensive analysis of the structural, functional, and coordinating environment in metal-binding sites [27]. Thus, we highlight two similar methods’ capabilities, yet with different approaches and aspects of algorithms in deriving authentic force field parameters for Fe^2+^ centers in DPD protein.

First and foremost, we have reported the generation of force field parameters using the original Seminario method [41]. We went further and exploited the collation features VFFDT Seminario method in obtaining the force field parameters of the same Fe^2+^ ions as a supportive measure [43]. This was performed by considering the dimeric functionality of the human DPD protein, which relies on the well-organized inter-chain electron transfer across eight Fe^2+^_4_S^2-^_4_ clusters complex. A double displacement reaction across the two chains leads to the activation and deactivation of the third most commonly prescribed anticancer (5F-U) drug globally [97]. It was remarkable that we successfully derived the desired force constants and bond distances for the Fe^2+^ centers using both Seminario approaches. Furthermore, validation of the generated parameters through MD simulations yielded satisfactory results. MD simulation is an integral tool of depicting atomistic or electronic information concerning effects of site-specific interactions on the reaction path, process, and the structure of evolution state [26, 31, 32].

Above all, the derived parameters could easily be incorporated into consolidated MM packages. Furthermore, we ascertained that irrespective of the DFT (B3LYP HF/6-31G* and (LSDA/LANL2DZ) logarithm application, the original Seminario approach is not inferior to the modified Seminario (collation features VFFDT) approach. For instance, the parameters obtained from other studies [36] did not address the coordinating geometry of the clusters in this study. Also, none of the studies focused on force field parameters for multiple clusters in a protein. Therefore, from the range of force field parameters generated from both approaches, it would be best to obtain averages of such force fields for future use in other similar systems. These averaged values will allow for some degree of transferability.

Most importantly, concerning the generation of AMBER force field parameters, the authors acknowledge no other compatible parameters for this unique system. The derived novel force field parameters have paved way for further simulations and enhance the mechanistic understanding of metal cluster function in the human DPD protein through higher-level MD simulation methods. Additionally, the derived parameters are currently being applied to study the structural and changes in stability effects due to existing mutations in the human DPD protein. Together, the results from these studies will provide the atomistic details of mutation effects involving the DPD protein. This will open a platform for the implementation of in silico cancer pharmacogenomics and drug discovery research on 5-FU drug efficacy and toxicity effects.

## Supplementary Materials

The following information is available free of charge on the IJMS publication website. Supplementary Materials can be found at www.mdpi.com/xxxxxxxxx/

**Figure S1**. An illustration Fe^2+^ center parameterization in the native human DPD Model 2, utilizing automated (VFFDT) Seminario approach.

**Figure S2**. An illustration of charge allocation to all the atoms coordinating with the metal center of subset clusters 1026A and 1027A for Model 1.

**Figure S3**. A representation of the root mean square of fluctuation (RMSF) for ligand bound (complex) and non-ligand bound (holo) DPD Model 1 during 150 ns simulation. **A)** Fe^2+^_4_S^2-^_4_ clusters in 1026 and 1027 located in domain 1. **B)** Fe^2+^_4_S^2-^_4_ clusters in 1028 and 1029 located in domain 5. The area of fluctuation coincides to the protein loop area while the iron cluster remains intact.

**Figure S4**. 3D structures of Model 2 MD simulations snapshots (timeframes) from regions exhibiting higher conformational changes with atomistic details represented **A**) at 110.0 ns for drug bound protein and **B)** at 70.4 ns for holo proteins (without 5-fluorouracil drug). The Fe^2+^ clusters remained intact throughout the different conformation timelines.

**Figure S5:** Color coded violin plots showing the crystal structure bond distances between the Fe^2+^ and S^2-^ derived by original (Model 1) and VFFDT [automated] (Model 2) Seminario method during 150 ns MD simulation. In Model 1, the orange and pink violin plots represent the holo and holo-drug bound complexes, respectively. In Model 2, the yellow and grey violin plots represent the holo and holo-drug complexes, respectively. The two clusters (1026 and 1027) represent the crystal structures (1H7X) unique Fe^2+^4S^2-^4 clusters coordination.

**Table S1**. Quality assessment of Human DPD protein modeled structures.

**Table S2**. Titratable residues in the human DPD protein and their respective pKa values.

**Table S3**. A representation of human DPD parameters and coordinate files for Model 1 (B3LYP/6-31G*): AMBER parameter file.

**Table S4**. A representation of human DPD parameters and coordinate files for Model 2 (LSDA/LANL2DZ): AM-BER_VFFDT parameter file.

**Table S5**. Listing of charge allocation to all the atoms interacting with the metal center (B3LYP/6-31G*)

**Table S6**. Comparison of **A** bond length, **B** internal and **C** external angles (Å) calculated with X-ray, DFT (B3LYP) and (LSDA/LANL2DZ) method for the molecular cluster model ([Fe^2+^_4_S^2-^_4_ (S-Cys)_3_(S-Gln)]) 1026A of Native DPD protein.

**Table S7**. Comparison of **A** bond length, **B** internal and **C** external angles (Å) calculated with X-ray, DFT (B3LYP) and (LSDA/LANL2DZ) method for the molecular cluster model ([Fe_4_S_4_(S-Cys)_4_]) 1027A of native DPD protein.

**Table S8**. Dihedral related force constants for X-ray and post-MD simulation for both models’ clusters ([Fe4S4(S-Cys)_3_(S-Gln)]) and ([Fe_4_S_4_(S-Cys)_4_]) of Native DPD protein.

**Table S9**. DPD. *pir* sequence file used for modeling human dihydropyramidine dehydrogenase structure based on pig crystal structure template and human target sequence

## Author Contributions

Conceptualization, Özlem Tastan Bishop; Formal analysis, Maureen Bilinga Tendwa, Lorna Chebon-Bore, Thommas Musyoka and Özlem Tastan Bishop; Funding acquisition, Özlem Tastan Bishop; Methodology, Maureen Bilinga Tendwa, Lorna Chebon-Bore, Kevin Lobb, Thommas Musyoka and Özlem Tastan Bishop; Resources, Özlem Tastan Bishop; Supervision, Thommas Musyoka and Özlem Tastan Bishop; Validation, Maureen Bilinga Tendwa; Visualization, Maureen Bilinga Tendwa; Writing – original draft, Maureen Bilinga Tendwa; Writing – review & editing, Lorna Chebon-Bore, Thommas Musyoka and Özlem Tastan Bishop.

## Funding

This work has been funded by H3ABioNet (U24HG006941) which is supported by National Institutes of Health. H3ABioNet is a Pan African Bioinformatics Network for the Human Heredity and Health in Africa (H3Africa) consortium. L.C.B. is funded by DELTAS Africa Initiative under Wellcome Trust (DELGEME grant number 107740/Z/15/Z) for PhD fellowship. The DELTAS Africa Initiative is an independent funding scheme of the African Academy of Sciences (AAS)’s Alliance for Accelerating Excellence in Science in Africa (AESA) and supported by the New Partnership for Africa’s Development Planning and Coordinating Agency (NEPAD Agency) with funding from the Wellcome Trust [DELGEME grant 107740/Z/15/Z] and the UK government. T.M.M. is funded as a postdoctoral fellow by Grand Challenges Africa program [GCA/DD/rnd3/023]. Grand Challenges Africa is a program of the African Academy of Sciences (AAS) implemented through the Alliance for Accelerating Excellence in Science in Africa (AESA) platform, an initiative of the AAS and the African Union Development Agency (AUDA-NEPAD). GC Africa is supported by the Bill & Melinda Gates Foundation (BMGF), Swedish International Development Cooperation Agency (SIDA), German Federal Ministry of Education and Research (BMBF), Medicines for Malaria Venture (MMV), and Drug Discovery and Development Centre of University of Cape Town (H3D). Expressed views, and conclusion drawn for the content of the publication are those of the author’s and are not necessarily credited the funder’s official views.

## Acknowledgments

The authors thank to the Center for High Performance Computing (CHPC), South Africa, for computational clusters and RUBi colleagues for their constructive discussions.

## Conflicts of Interest

The authors declare no conflict of interest. The funders had no role in the study design, collection, analysis or interpretation of the data and decision to publish the data.in the writing of the manuscript, or in the decision to publish the results.

### Abbreviations

3D: Three-dimensional
5-FU: Five-fluorouracil
ACPYPE: Ante-Chamber Python Parser interface
CHPC: Center for high performance computing
CPU: Central processing unit
DPD: Dihydropyrimidine dehydrogenase
FAD: Flavin adenine dinucleotide
FMN: Flavin mononucleotide
MCBP: Metal center parameter builder
MD: Molecular dynamics
MM: Molecular mechanics
NADP: Nicotinamide adenine dinucleotide phosphate
PBC: Periodic boundary conditions
PDB: Protein Data bank
PME: Particle mesh Ewald
RESP: Restricted electrostatic potential
QM: Quantum mechanics
URF: Five fluorouracil
VFFDT: Visual force field derivation toolkit
WT: Wild type

## Notes

### Competing Interest Statement

The authors have declared no competing interest.

## References

1. Dobritzsch D., Ricagno, S., Schneider, G., Schnackerz, K.D., and Lindqvist, Y. Crystal structure of the productive ternary complex of dihydropyrimidine dehydrogenase with NADPH and 5-iodouracil implications for mechanism of inhibition and electron transfer. J. Biol. Chem, 2002, 277, 13155–13166. Doi.org/10.1074/jbc.M111877200.

2. Dobritzsch, D., Schneider, G., Schnackerz, K.D., and Lindqvist, Y. Crystal structure of dihydropyrimidine dehydrogenase, a major determinant of the pharmacokinetics of the anti-cancer drug 5-fluorouracil. EMBO J., 2001, 20, 650–660. Doi.10.1093/emboj/20.4.650.

3. Miteva-Marcheva NN, Ivanov HY, Dimitrov DK, Stoyanova VK: Application of pharmacogenetics in oncology. Biomark. Res. 2020, 8,1, 1–10. Doi.org/10.1186/s40364-020-00213-4.

4. Mazzuca, Federica, Marina Borro, Andrea Botticelli, Eva Mazzotti, Luca Marchetti, Giovanna Gentile, Marco La Torre, Luana Lionetto, Maurizio Simmaco, and Paolo Marchetti. Pre-Treatment Evaluation of 5-Fluorouracil Degradation Rate: Association of Poor and Ultra-Rapid Metabolism with Severe Toxicity in a Colorectal Cancer Patients Cohort. Oncotarget, 2016, 7, 20612–20620. Doi.10.18632/oncotarget.7991.

5. Wigle, T.J., Tsvetkova, E.V., Welch, S.A., and Kim, R.B. DPYD and fluorouracil-based chemotherapy: Mini review and case report. Int. J. Pharm., 2019, 11, 199. Doi: 10.3390/pharmaceutics11050199.

6. García-González X, Kaczmarczyk B, Abarca-Zabalía J, Thomas F, García-Alfonso P, Robles L, Pachón V, Vaz Á, Salvador-Martín S, Sanjurjo-Sáez M: New DPYD variants causing DPD deficiency in patients treated with fluoropyrimidine. Cancer Chemother Pharmacol. 2020, 86(1):45–54. Doi: 10.1007/s00280-020-04093-1.

7. Tozer T, Heale K, Manto Chagas C, de Barros ALB, Alisaraie L: Interdomain twists of human thymidine phosphorylase and its active–inactive conformations: Binding of 5-FU and its analogues to human thymidine phosphorylase versus dihydropyrimidine dehydrogenase. Chem Biol Drug Des, 2019, 94, 5, 1956–1972. Doi.org/10.1111/cbdd.13596.

8. Berman HM, Westbrook J, Feng Z, Gilliland G, Bhat TN, Weissig H, Shindyalov IN, Bourne PE: The protein data bank. Nucleic Acids Res, 2000, 28, 1, 235–242. Doi.org/10.1093/nar/28.1.235.

9. Lohkamp, B., Voevodskaya, N., Lindqvist, Y., and Dobritzsch, D. Insights into the mechanism of dihydropyrimidine dehydrogenase from site-directed mutagenesis targeting the active site loop and redox cofactor coordination. Biochim Biophys Acta Proteins, 2010, 1804, 2198–2206. Doi.10.1016/j.bbapap.2010.08.014.

10. Schnackerz, K.D., Dobritzsch, D., Lindqvist, Y., and Cook, P.F. Dihydropyrimidine dehydrogenase: a flavoprotein with four iron–sulfur clusters. BBA-Proteins Proteom, 2004, 1701, 61–74. Doi.10.1016/j.bbapap.2004.06.009.

11. Fagan, R.L., and Palfey, B.A.. Flavin-dependent enzymes. Enzyme Res, 2020, 47, 1–36. Doi.org/10.1016/bs.enz.2020.06.006.

12. Mander, L., and Liu, H.-W. Comprehensive natural products II: Chemistry and Biology, 2010, Volume 1, (Elsevier).

13. Podschun, B., Wahler, G., and Schnackerz, K.D. Purification and characterization of dihydropyrimidine dehydrogenase from pig liver. Eur. J. Chem., 1989, 185, 219–224. Doi.org/10.1111/j.1432-1033.1989.tb15105.x.

14. Podschun, B., Cook, P., and Schnackerz, K. Kinetic mechanism of dihydropyrimidine dehydrogenase from pig liver. J. Biol. Chem, 1990, 265, 12966–12972. Doi.10.1016/S0021-9258(19)38254-7.

15. Hagen, W., Vanoni, M., Rosenbaum, K., and Schnackerz, K. On the iron–sulfur clusters in the complex redox enzyme dihydropyrimidine dehydrogenase. Eur. J. Chem., 2000, 267, 3640–3646. Doi.10.1046/j.1432-1327.2000.01393.x.

16. Dudev, T., and Lim, C. Metal binding affinity and selectivity in metalloproteins: insights from computational studies. Annu. Rev. Biophys, 2008, 37, 97–116. Doi.10.1146/annurev.biophys.37.032807.125811.

17. Ziller, A., Yadav, R.K., Capdevila, M., Reddy, M.S., Vallon, L., Marmeisse, R., Atrian, S., Palacios, Ò., and Fraissinet-Tachet, L. Metagenomics analysis reveals a new metallothionein family: sequence and metal-binding features of new environmental cysteine-rich proteins. J. Inorg. Biochem, 2017, 167, 1–11. Doi. 10.1016/j.jinorgbio.2016.11.017.

18. Jing, Z., Liu, C., Qi, R., and Ren, P. Many-body effect determines the selectivity for Ca2+ and Mg2+ in proteins. PNAS, 2018, 115, E7495–E7501. Doi.10.1073/pnas.1805049115.

19. Valasatava Y, Rosato A, Furnham N, Thornton JM, Andreini C: To what extent do structural changes in catalytic metal sites affect enzyme function? J. Inorg. Biochem, 2018, 179:40–53. Doi.10.1016/j.jinorgbio.2017.11.002.

20. Barber-Zucker S, Shaanan B, Zarivach R: Transition metal binding selectivity in proteins and its correlation with the phylogenomic classification of the cation diffusion facilitator protein family. Sci. Rep., 2017, 7(1):1–12. Doi.10.1038/s41598-017-16777-5.

21. Lopes PE, Guvench O, MacKerell AD: Current status of protein force fields for molecular dynamics simulations. In: J. Mol. Model.. Springer, 2015, 47–71. Doi.10.1007/978-1-4939-1465-4_3.

22. Robustelli P, Piana S, Shaw DE: Developing a molecular dynamics force field for both folded and disordered protein states. Proc. Natl. Acad. Sci. U. S. A., 2018, 115,21,E4758–E4766. Doi.org/10.1073/pnas.1800690115.

23. Singh S, Singh VK: Molecular Dynamics Simulation: Methods and Application. In: Frontiers in Protein Structure, Function, and Dynamics. Springer; 2020: 213–238.

24. Shi, W., and Chance, M.R. Metalloproteomics: forward and reverse approaches in metalloprotein structural and functional characterization. Curr Opin Chem Biol, 2011, 15, 144–148. Doi.10.1016/j.cbpa.2010.11.004.

25. Musyoka, T.M., Tastan Bishop, Ö., Lobb, K., and Moses, V. The determination of CHARMM force field parameters for the Mg2+ containing HIV-1 integrase. Chem. Phys. Lett., 2018, 711, 1–7. Doi.10.1016/j.cplett.2018.09.019.

26. Bray, S., and Furriols, M. Notch pathway: making sense of suppressor of hairless. Curr. Biol., 2001, 11, R217–R221. Doi.org/10.1016/S0960-9822(01)00109-9.

27. Carloni P, Rothlisberger U, Parrinello M: The role and perspective of ab initio molecular dynamics in the study of biological systems. Acc. Chem. Res., 2002, 35(6):455–464. Doi.10.1021/ar010018u.

28. Hancock, R.D. Molecular mechanics calculations and metal ion recognition. Acc. Chem. Res., 1990, 23, 253–257. Doi.org/10.1021/ar00176a003.

29. Stote RH, Karplus M: Zinc binding in proteins and solution: a simple but accurate nonbonded representation. Proteins: Struct., Funct., 1995, 23,1,12–31. Doi.org/10.1021/ar010018u.

30. Pang, Y.-P. Novel zinc protein molecular dynamics simulations: Steps toward antiangiogenesis for cancer treatment. J. Mol. Model., 1999, 5, 196–202. Doi.org/10.1007/s008940050119.

31. Åqvist, J. Modeling of ion-ligand interactions in solutions and biomolecules. J. Mol. Struct., 1992, 256, 135–152. Doi.org/10.1016/0166-1280(92)87163-T.

32. Sakharov, D.V., and Lim, C. Zn protein simulations including charge transfer and local polarization effects. J. Am. Chem. Soc., 2005, 127, 4921–4929. Doi.org/10.1021/ja0429115.

33. Dal Peraro, M., Spiegel, K., Lamoureux, G., De Vivo, M., DeGrado, W.F., and Klein, M.L. Modeling the charge distribution at metal sites in proteins for molecular dynamics simulations. J. Struct. Biol, 2007, 157, 444–453. Doi.10.1016/j.jsb.2006.10.019.

34. Zhu, T., Xiao, X., Ji, C., and Zhang, J.Z. A new quantum calibrated force field for zinc–protein complex. J. Chem. Theory Comput., 2013, 9, 1788–1798. Doi.10.1021/ct301091z.

35. Vedani, A., and Huhta, D.W. A new force field for modeling metalloproteins. J. Am. Chem. Soc., 1990, 112, 4759–4767. Doi.org/10.1021/ja00168a021.

36. Carvalho, A.T., Teixeira, A.F., and Ramos, M.J. Parameters for molecular dynamics simulations of iron-sulfur proteins. J. Comput. Chem., 2012, 34, 1540–1548. Doi.org/10.1002/jcc.23287.

37. Sheik Amamuddy, O.S., Musyoka, T.M., Boateng, R.A., Zabo, S., and Tastan Bishop, Ö. Determining the unbinding events and conserved motions associated with the pyrazinamide release due to resistance mutations of Mycobacterium tuberculosis pyrazinamidase. Comput. Struct. Biotechnol. J., 2020, 18, 1103-1120. Doi: 10.1016/j.csbj.2020.05.009.

38. Teixeira, M.H., Curtolo, F., Camilo, S.R., Field, M.J., Zheng, P., Li, H., and Arantes, G.M. Modeling the Hydrolysis of Iron–Sulfur Clusters. J Chem Inf Model, 2019, 60, 653–660. Doi.10.1021/acs.jcim.9b00881.

39. Oda, A., Yamaotsu, N., and Hirono, S. New AMBER force field parameters of heme iron for cytochrome P450s determined by quantum chemical calculations of simplified models. J. Comput. Chem., 2005, 26, 818–826. Doi.org/10.1002/jcc.20221.

40. Li, P., and Merz Jr, K.M. Metal ion modeling using classical mechanics. Chem. Rev., 2017, 117, 1564–1686. Doi.org/10.1021/acs.chemrev.6b00440.

41. Seminario, J.M. Calculation of intramolecular force fields from second-derivative tensors. Int. J. Quantum Chem., 1996, 60, 1271–1277. Doi.org/10.1002/(SICI)1097-461X(1996)60:7<1271::AID-QUA8>3.0.CO;2-W

42. Li, P., and Merz Jr, K.M. MCPB. py: A python based metal center parameter builder. J Chem Inf Model, 2016, 56, 4, 599–604. Doi.org/10.1021/acs.jcim.5b00674

43. Zheng, S., Tang, Q., He, J., Du, S., Xu, S., Wang, C., Xu, Y., and Lin, F. VFFDT: a new software for preparing AMBER force field parameters for metal-containing molecular systems. J Chem Inf Model, 2016, 56, 811–818. Doi.10.1021/acs.jcim.5b00687

44. Kelm, S., Shi, J., and Deane, C.M. MEDELLER: homology-based coordinate generation for membrane proteins. Bioinformatics, 2010, 26, 2833–2840. Doi.org/10.1093/bioinformatics/btq554

45. Biovia, D.S., and Dsme, R. 2015. San Diego: Dassault Systèmes. (Release).

46. Shen, M.y., and Sali, A. Statistical potential for assessment and prediction of protein structures. Protein Sci., 2006, 15, 2507–2524. Doi.10.1110/ps.062416606

47. Eramian, D., Eswar, N., Shen, M.Y., and Sali, A. How well can the accuracy of comparative protein structure models be predicted? Protein Sci., 2008, 17, 1881–1893. Doi.10.1110/ps.036061.108.

48. Eisenberg D, Lüthy R, Bowie JU: [20] VERIFY3D: assessment of protein models with three-dimensional profiles. Meth. Enzymol, 1997, 277, 396–404. Doi.org/10.1016/S0076-6879(97)77022-8.

49. Benkert P, Künzli M, Schwede T: QMEAN server for protein model quality estimation. Nucleic Acids Res., 2002, 37(Suppl_2):W510–W514. Doi: 10.1093/nar/gkp322

50. Wiederstein, M., and Sippl, M.J. ProSA-web: interactive web service for the recognition of errors in three-dimensional structures of proteins. Nucleic Acids Res., 2007, 35, W407–W410. Doi.org/10.1016/S0959-440X(98)80156-5

51. Laskowski, R.A., MacArthur, M.W., and Thornton, J.M. Validation of protein models derived from experiment. Curr. Opin. Struct. Biol., 1998, 8, 631–639. Doi.org/10.1016/S0959-440X(98)80156-5

52. Van Der Spoel D, Lindahl E, Hess B, Groenhof G, Mark AE, Berendsen HJ: GROMACS: fast, flexible, and free. J. Comput. Chem., 2005, 26(16):1701–1718. Doi: 10.1002/jcc.20291

53. Anandakrishnan, R., Aguilar, B., and Onufriev, A.V. H++ 3.0: automating p K prediction and the preparation of biomolecular structures for atomistic molecular modeling and simulations. Nucleic Acids Res., 2012, 40, W537–W541. Doi.10.1093/nar/gks375.

54. Bowers, K.J., Chow, D.E., Xu, H., Dror, R.O., Eastwood, M.P., Gregersen, B.A., Klepeis, J.L., Kolossvary, I., Moraes, M.A., and Sacerdoti, F.D. Scalable algorithms for molecular dynamics simulations on commodity clusters. IEEE, 2006, pp. 43–43. Doi.org/10.1145/1188455.1188544.

55. Khairallah, A., Tastan Bishop, Ö., and Moses, V. AMBER force field parameters for the Zn (II) ions of the tunneling-fold enzymes GTP cyclohydrolase I and 6-pyruvoyl tetrahydropterin synthase. J. Biomol. Struct. Dyn., 2020, 28, 1–18. Doi.10.1080/07391102.2020.1796800.

56. Li, X., Hayik, S.A., and Merz Jr, K.M. QM/MM X-ray refinement of zinc metalloenzymes. J. Inorg. Biochem., 2010, 104, 512–522. Doi.10.1016/j.jinorgbio.2009.12.022.

57. Tan, L.L., Holm, R., and Lee, S.C. Structural analysis of cubane-type iron clusters. Polyhedron, 2013, 58, 206–217. Doi.10.1016/j.poly.2013.02.031.

58. Moriarty, N.W., and Adams, P.D. Iron–sulfur clusters have no right angles. Acta Crystallographica Section D. J. Struct. Biol., 2019, 75, 16–20. Doi.org/10.1107/S205979831801519X.

59. Harding MM: Small revisions to predicted distances around metal sites in proteins. Acta Crystallogr D Biol Crystallogr., 2006, 62(6):678–682. Doi: 10.1107/S0907444906014594

60. Tuccinardi, T., Martinelli, A., Nuti, E., Carelli, P., Balzano, F., Uccello-Barretta, G., Murphy, G., and Rossello, A. Amber force field implementation, molecular modeling study, synthesis and MMP-1/MMP-2 inhibition profile of (R)-and (S)-N-hydroxy-2-(N-isopropoxybiphenyl-4-ylsulfonamido)-3-methylbutanamides. Bioorg. Med. Chem., 2006, 14, 4260–4276. Doi.10.1016/j.bmc.2006.01.056.

61. Smith, D.M., Xiong, Y., Straatsma, T., Rosso, K.M., and Squier, T.C. Force-field development and molecular dynamics of [NiFe] hydrogenase. J. Chem. Theory Comput., 2012, 8, 2103–2114. Doi.org/10.1021/ct300185u.

62. Wei, C., Lazim, R., and Zhang, D. Importance of polarization effect in the study of metalloproteins: Application of polarized protein specific charge scheme in predicting the reduction potential of azurin. Proteins, 2014, 82, 2209–2219. Doi.org/10.1002/prot.24584.

63. Humphrey W, Dalke A, Schulten K: VMD: visual molecular dynamics. J. Mol. Graph., 1996, 14(1):33–38. Doi: 10.1016/0263-7855(96)00018-5.

64. Haspel, N., Moll, M., Baker, M.L., Chiu, W., and Kavraki, L.E. Tracing conformational changes in proteins. BMC Struct. Biol., 2010, 10, 1, S1. Doi.org/10.1186/1472-6807-10-S1-S1.

65. Orellana, L. Large-scale conformational changes and protein function: breaking the in silico barrier. Front. Mol. Biosci, 2019, 6, 117–136. Doi.org/10.3389/fmolb.2019.00117.

66. Torrens F, Castellano G: Calculation of partition coefficients of Fe–S/Se protein models. 2007. Doi: 10.3390/ecsoc-11-01365

67. Harding, M.M. The geometry of metal–ligand interactions relevant to proteins. Acta Crystallogr., Sect. D: Biol. Crystallogr, 1999, 55, 1432–1443. Doi.10.1107/s0907444999007374

68. Sheik Amamuddy OS, Glenister M, Tastan Bishop, Ö. MDM-TASK-web: A web platform for protein dynamic residue networks and modal analysis. bioRxiv, 2021. Doi.org/10.1101/2021.01.29.428734.

69. Tastan Bishop, Ö., De Beer TA, Joubert F: Protein homology modelling and its use in South Africa. S. Afr. J. Sci., 2008, 104(1-2):2–6.

70. Söding J, Biegert A, Lupas AN: The HHpred interactive server for protein homology detection and structure prediction. Nucleic Acids Res., 2005, 33(Suppl_2):W244–W248. Doi: 10.1093/nar/gki408.

71. Ryde, U. Molecular dynamics simulations of alcohol dehydrogenase with a four-or five-coordinate catalytic zinc ion. Proteins, 1995, 21, 40–56. Doi.org/10.1002/prot.340210106.

72. Ryde, U. Combined quantum and molecular mechanics calculations on metalloproteins. Curr Opin Chem Biol, 2003, 7, 136–142. Doi.org/10.1016/S1367-5931(02)00016-9.

73. Lee, C., Yang, W., and Parr, R.G. Development of the Colle-Salvetti correlation-energy formula into a functional of the electron density. Phys. Rev. Lett., 1988, B 37, 785. Doi.10.1103/physrevb.37.785.

74. Cornell WD, Cieplak P, Bayly CI, Gould IR, Merz KM, Ferguson DM, Spellmeyer DC, Fox T, Caldwell JW, Kollman PA: A second generation force field for the simulation of proteins, nucleic acids, and organic molecules. J. Am. Chem., 1995, 117,19, :5179–5197. Doi.org/10.1021/ja00124a002

75. Frisch, M. Trucks, GW Schlegel, H. et al. B., et a/., Gaussian 3. 1998.

76. Case, D., Darden, T., Cheatham, T., Simmerling, C., Wang, J., Duke, R., Luo, R., Crowley, M., Walker, R., and Zhang, W. AMBER 12. 2012. University of California. San Francisco.

77. Bayly, C.I., Cieplak, P., Cornell, W., and Kollman, P.A. A well-behaved electrostatic potential based method using charge restraints for deriving atomic charges: the RESP model. J. Phys. Chem., 1993, 97, 10269–10280. Doi.org/10.1021/j100142a004.

78. Ehlers, A., Böhme, M., Dapprich, S., Gobbi, A., Höllwarth, A., Jonas, V., Köhler, K., Stegmann, R., Veldkamp, A., and Frenking, G. A set of f-polarization functions for pseudo-potential basis sets of the transition metals Sc Cu, Y Ag and La Au. Chem. Phys. Lett., 1993, 208, 111–114. Doi.org/10.1016/0009-2614(93)80086-5.

79. Hehre, W.J. Ab initio molecular orbital theory. Acc. Chem. Res., 1976, 9, 399–406. Doi.org/10.1021/ar50107a003.

80. Grimme, S., Bannwarth, C., and Shushkov, P. A robust and accurate tight-binding quantum chemical method for structures, vibrational frequencies, and noncovalent interactions of large molecular systems parametrized for all spd-block elements (Z= 1–86). J. Chem. Theory Comput., 2017, 13, 1989–2009. Doi: 10.1021/acs.jctc.7b00118.

81. Schafmeister, C., Ross, W., and Romanovski, V.LEaP. 1995. University of California, San Francisco.

82. Maier JA, Martinez C, Kasavajhala K, Wickstrom L, Hauser KE, Simmerling C: ff14SB: improving the accuracy of protein side chain and backbone parameters from ff99SB. J. Chem. Theory Comput., 2015, 11(8):3696–3713. Doi.org/10.1021/acs.jctc.5b00255

83. Da Silva, A.W.S., and Vranken, W.F. ACPYPE-Antechamber python parser interface. BMC Res, 2012, 5, 367. Doi.org/10.1186/1756-0500-5-367.

84. Glättli A, Daura X, van Gunsteren WF: Derivation of an improved simple point charge model for liquid water: SPC/A and SPC/L. J. Chem. Phys., 2002, 116, 22, 9811–9828. Doi.10.1063/1.1476316.

85. Hockney RW, Goel S, Eastwood J: Quiet high-resolution computer models of a plasma. J. Chem. Theory Comput.,1974, 14(2):148–158.

86. Petersen HG: Accuracy and efficiency of the particle mesh Ewald method. J. Chem. Phys. 1995, 103,9, 3668–3679. Doi.org/10.1063/1.470043.

87. Essman U, Perera L, Berkowitz M, Darden T, Lee H, Pedersen L: A smooth particle mesh ewald potential. J Chem Phys, 1995, 103:8577–8592.

88. Ryckaert, J.-P., Ciccotti, G., and Berendsen, H.J. Numerical integration of the cartesian equations of motion of a system with constraints: molecular dynamics of n-alkanes. J. Comput. Phys., 1977, 23, 327–341. Doi.org/10.1016/0021-9991(77)90098-5

89. Team R: RStudio: integrated development for R. RStudio, Inc, Boston, MA URL http://www.rstudiocom, 2015, 42:14. http://www.rstudio.com/.

90. Brown DK, Penkler DL, Sheik Amamuddy O, Ross C, Atilgan AR, Atilgan C, Tastan Bishop Ö: MD-TASK: a software suite for analyzing molecular dynamics trajectories. Bioinformatics, 2017, 33(17):2768-2771.90. Doi: 10.1093/bioinformatics/btx349

91. Schrodinger, L. The PyMOL molecular graphics system. 2010, Version 1, 0., Doi, 10.1021/acs.jctc.5b00864.

92. Kluyver, T., Ragan-Kelley, B., Pérez, F., Granger, B.E., Bussonnier, M., Frederic, J., Kelley, K., Hamrick, J.B., Grout, J., and Corlay, S. Jupyter Notebooks-a publishing format for reproducible computational workflows. In ELPUB. 2016, pp. 87–90.

93. Hunter, J.D. Matplotlib: A 2D graphics environment. Comput Sci Eng, 2007, 9, 90–95. Doi.org/10.1109/MCSE.2007.55.

94. McKinney, W. Data structures for statistical computing in python. In Proceedings of the 9th Python in Science Conference, 2010, 445, 51–56.

95. Walt, S.v.d., Colbert, S.C., and Varoquaux, G. The NumPy array: a structure for efficient numerical computation. Comput Sci Eng, 2011, 13, 22–30. Doi.10.1109/MCSE.2011.37.

96. Nguyen, H., Case, D.A., and Rose, A.S. NGLview–interactive molecular graphics for Jupyter notebooks. Bioinform, 2018, 34, 1241–1242. Doi.10.1093/bioinformatics/btx789

97. Sara, J.D., Kaur, J., Khodadadi, R., Rehman, M., Lobo, R., Chakrabarti, S., Herrmann, J., Lerman, A., and Grothey, A. 5-fluorouracil and cardiotoxicity: a review. Ther Adv Med Oncol, 2018, 10, Doi. 10.1177/1758835918780140

