## Supplementary data for "Force field parameters for Fe^2+^_4_S^2-^_4_ clusters of dihydropyrimidine dehydrogenase, the 5-fluorouracil cancer drug deactivation protein: a step towards *in silico* pharmacogenomics studies"

### Table of Contents

**Figure S1.** An illustration $Fe^{2+}$ center parameterization in the native human DPD Model 2, utilizing automated (VFFDT) Seminario approach.
**Figure S2**. An illustration of charge allocation to all the atoms coordinating with the metal center of subset clusters 1026A and 1027A for Model 1.
**Figure S3.** A representation of the root mean square of fluctuation (RMSF) for ligand bound (complex) and non-ligand bound (holo) DPD Model 1 during 150 ns simulation. **A)** $Fe^{2+}{}_4S^{2-}{}_4$ clusters in 1026 and 1027 located in domain 1. **B)** $Fe^{2+}{}_4S^{2-}{}_4$ clusters in 1028 and 1029 located in domain 5. The area of fluctuation coincides to the protein loop area while the iron cluster remains intact.
**Figure S4.** 3D structures of Model 2 MD simulations snapshots (timeframes) from regions exhibiting higher conformational changes with atomistic details represented **A**) at 110.0 ns for drug bound protein and **B)** at 70.4 ns for holo proteins (without 5-fluorouracil drug). The $Fe^{2+}$ clusters remained intact throughout the different conformation timelines.
**Figure S5:** Color coded violin plots showing the crystal structure bond distances between the $Fe^{2+}$ and $S^{2-}$ derived by original (Model 1) and VFFDT [automated] (Model 2) Seminario method during 150 ns MD simulation. In Model 1, the orange and pink violin plots represent the holo and holo-drug bound complexes, respectively. In Model 2, the yellow and grey violin plots represent the holo and holo-drug complexes, respectively. The two clusters (1026 and 1027) represent the crystal structures (1H7X) unique $Fe^{2+}4S^{2-}4$ clusters coordination.
**Table S1.** Quality assessment of Human DPD protein modeled structures.
**Table S2**. Titratable residues in the human DPD protein and their respective pKa values.
**Table S3.** A representation of human DPD parameters and coordinate files for Model 1 (B3LYP/6-31G*): AMBER parameter file.
**Table S4.** A representation of human DPD parameters and coordinate files for Model 2 (LSDA/LANL2DZ): AMBER_VFFDT parameter file.
**Table S5**. Listing of charge allocation to all the atoms interacting with the metal center (B3LYP/6-31G*)
**Table S6.** Comparison of **A** bond length, **B** internal and **C** external angles (Å) calculated with X-ray, DFT (B3LYP) and (LSDA/LANL2DZ) method for the molecular cluster model ([$Fe^{2+}{}_4S^{2-}{}_4$ (S-Cys)$_3$(S-Gln)]) 1026A of Native DPD protein.
**Table S7.** Comparison of **A** bond length, **B** internal and **C** external angles (Å) calculated with X-ray, DFT (B3LYP) and (LSDA/LANL2DZ) method for the molecular cluster model ([$Fe_4S_4$(S-Cys)$_4$]) 1027A of native DPD protein.
**Table S8.** Dihedral related force constants for X-ray and post-MD simulation for both models' clusters ([Fe4S4(S-Cys)$_3$(S-Gln)]) and ([$Fe_4S_4$(S-Cys)$_4$]) of Native DPD protein.
**Table S9**. DPD *.pir* sequence file used for modeling human dihydropyramidine dehydrogenase structure based on pig crystal structure template and human target sequence

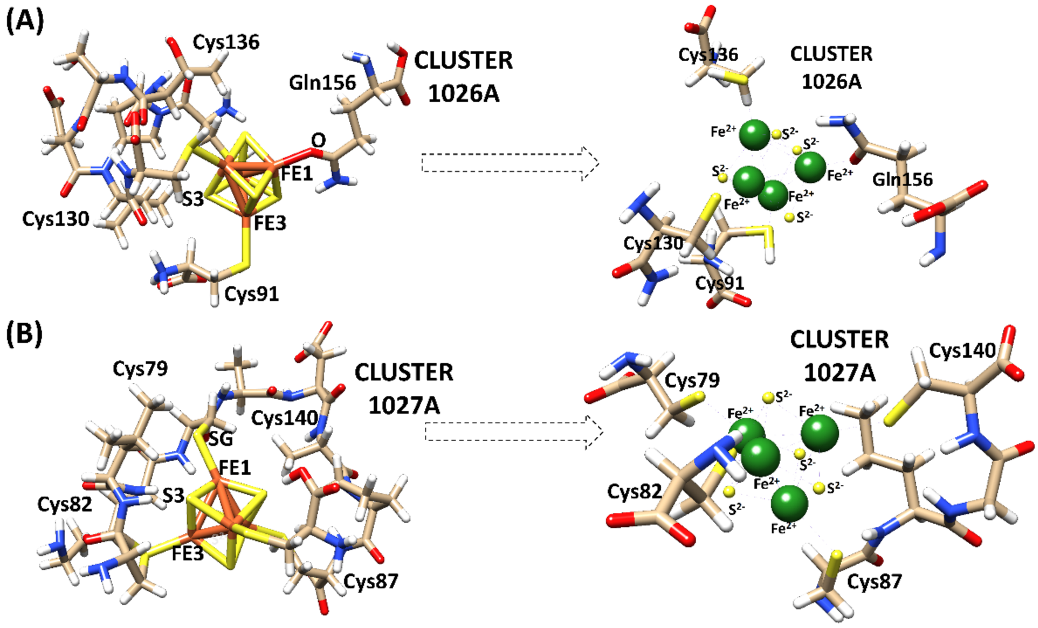


**Figure S1.** An illustration $Fe^{2+}$ center parameterization in the native human DPD Model 2, utilizing automated (VFFDT) Seminario approach A**)** showing a 3D representation of coordinating $Fe^{2+}$ sphere for cluster 1026A and an adjacent structure which has undergone parameterization, similarly **B)** shows a 3D representation of coordinating $Fe^{2+}$ sphere for cluster 1027A and an adjacent structure which has undergone parameterization.

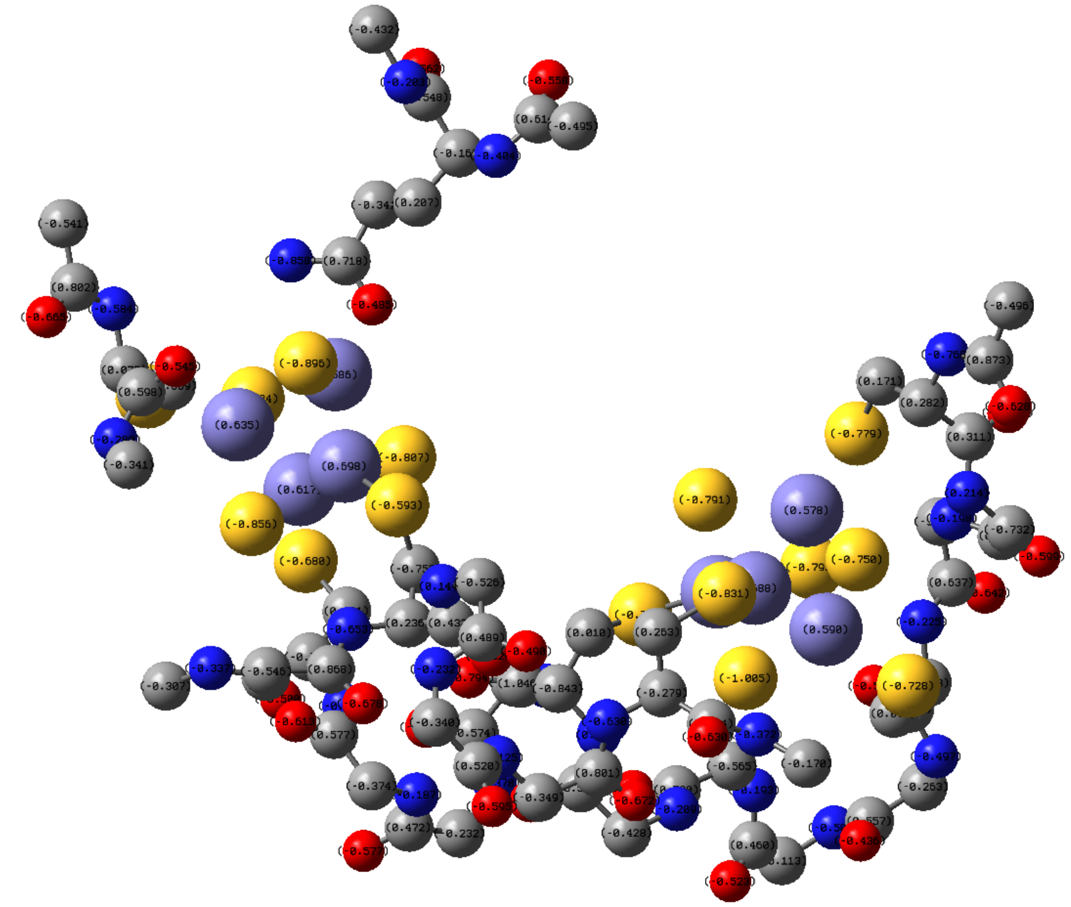

**Figure S2.** An illustration of charge allocation to all the atoms coordinating with the metal center of subset clusters 1026A and 1027A for Model 1**.** Purple, yellow, red, blue, and grey spheres represent Iron, sulfur, oxygen, nitrogen, and carbon atoms, respectively.

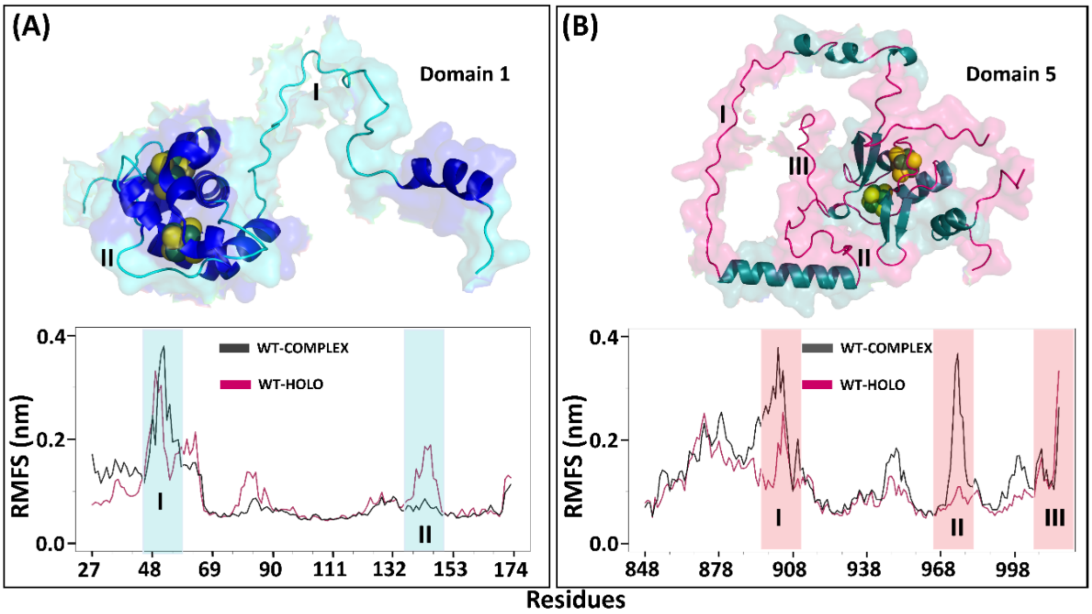


**Figure S3.** A representation of the root mean square of fluctuation (RMSF) for ligand bound (complex) and non-ligand bound (holo) DPD Model 1 during 150 ns simulation**. A)** $Fe^{2+}{}_4S^{2-}{}_4$ clusters in 1026 and 1027 located in domain 1. **B)** $Fe^{2+}{}_4S^{2-}{}_4$ clusters in 1028 and 1029 located in domain 5. The area of fluctuation coincides to the protein loop area while the iron cluster remains intact.

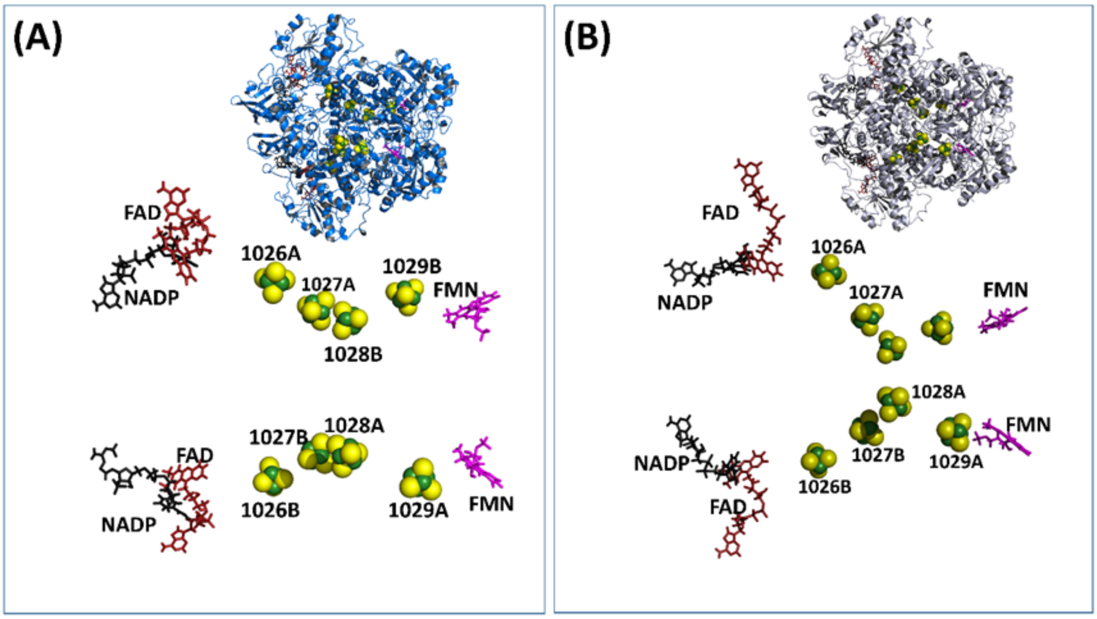


**Figure S4.** 3D structures of Model 2 MD simulations snapshots (timeframes) from regions exhibiting higher conformational changes with atomistic details represented **A**) at 110.0 ns for drug bound protein and **B)** at 70.4 ns for holo proteins (without 5-fluorouracil drug). The $Fe^{2+}$ clusters remained intact throughout the different conformation timelines.

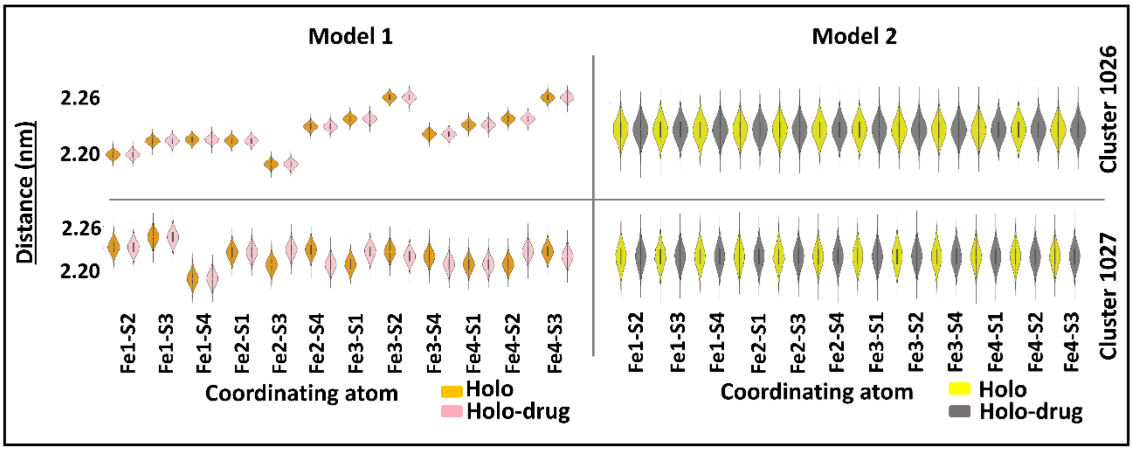


**Figure S5:** Color coded violin plots showing the crystal structure bond distances between the $Fe^{2+}$ and $S^{2-}$ derived by original (Model 1) and VFFDT [automated] (Model 2) Seminario method during 150 ns MD simulation. In Model 1, the orange and pink violin plots represent the holo and holo-drug bound complexes, respectively. In Model 2, the yellow and grey violin plots represent the holo and holo-drug complexes, respectively. The two clusters (1026 and 1027) represent the crystal structures (1H7X) unique $Fe^{2+}4S^{2-}4$ clusters coordination.

**Table S1:** Quality assessment of Human DPD protein modeled structures before and after 150 ns molecular dynamic simulation

| | | Pre and Post MD human DPD protein model structure quality validation | | | | | | |
|---|---|---|---|---|---|---|---|---|
| **PROTEINS** | | VERIFY 3D (%) | **QMEAN6** | **ProSA** | | **PROCHECK (%)** | | |
| | | **3D-ID score** | **QMEAN score** | **Monomers Z-score** | | **Ramachandran (residues location)** | | |
| | | | | **A** | **B** | **Favored** | **Allowed** | **Disallowed** |
| Template | Pre-MD | 85.79 | 0.91 | -13.56 | -13.47 | 93.5 | 6.4 | 0.1 |
| | Post-MD | 84.30 | 0.84 | -12.2 | -12.53 | 84.4 | 15.1 | 0.0 |
| Holo | Pre-MD | 85.01 | 0.90 | -13.41 | -13.44 | 89.4 | 10.2 | 0.0 |
| | Post-MD | 82.46 | 0.83 | -12.22 | -12.17 | 83.8 | 16.0 | 0.2 |
| Holo-drug | Pre-MD | 85.01 | 0.89 | - 13.42 | -13.45 | 92.2 | 7.7 | 0.1 |
| | Post-MD | 81.92 | 0.83 | - 12.61 | -12.62 | 85.8 | 14.1 | 0.1 |

MD: Molecular dynamics simulations

**Table S2**. Titratable residues in the human DPD protein and their respective pKa values

| Residue | pK (int) | pKa_(1/2) | Residue | pK (int) | pKa_(1/2) | Residue | pK (int) | pKa_(1/2) |
|---|---|---|---|---|---|---|---|---|
| **Chain-A** | | | | | | | | |
| NTALA-2 | 7.417 | 7.972 | ARG-678 | 11.484 | >12.000 | CYS-324 | 8.990 | 9.066 |
| LYS-7 | 10.092 | 11.678 | CYS-684 | 11.950 | >12.000 | HIS-325 | 6.186 | 6.037 |
| ASP-8 | 4.515 | 0.918 | ASP-687 | 3.873 | 1.358 | ARG-332 | 12.252 | >12.000 |
| ASP-10 | 3.600 | 3.978 | GLU-689 | 4.008 | 3.203 | ASP-342 | 2.523 | <0.000 |
| ASP-11 | 3.436 | 3.435 | ARG-692 | 11.329 | >12.000 | ASP-346 | 4.452 | <0.000 |
| GLU-13 | 4.390 | 4.841 | CYS-695 | 11.905 | >12.000 | CYS-347 | 9.618 | >12.000 |
| ARG-21 | 11.752 | >12.000 | ARG-696 | 11.889 | >12.000 | ARG-353 | 12.006 | >12.000 |
| HIS-25 | 6.649 | 7.179 | ARG-699 | 12.639 | >12.000 | CYS-354 | 12.003 | >12.000 |
| CYS-29 | 9.529 | 10.217 | LYS-709 | 6.035 | 11.023 | ARG-357 | 11.430 | >12.000 |
| LYS-34 | 9.536 | >12.000 | ASP-716 | 2.681 | <0.000 | ARG-358 | 11.391 | >12.000 |
| LYS-35 | 9.563 | >12.000 | ARG-722 | 11.424 | >12.000 | ARG-364 | 11.276 | >12.000 |
| ASP-37 | 5.142 | 1.200 | LYS-725 | 10.304 | 10.679 | LYS-365 | 9.534 | 9.163 |
| LYS-38 | 9.329 | >12.000 | GLU-726 | 5.068 | 2.383 | ARG-371 | 11.195 | >12.000 |
| LYS-39 | 10.143 | 10.937 | ASP-730 | 5.918 | 2.792 | GLU-375 | 3.112 | <0.000 |
| HIS-40 | 6.461 | 6.871 | LYS-745 | 9.516 | >12.000 | GLU-376 | 4.400 | 6.860 |
| LYS-42 | 9.998 | >12.000 | ASP-747 | 3.896 | <0.000 | GLU-378 | 4.413 | <0.000 |
| ARG-43 | 13.175 | >12.000 | LYS-758 | 10.327 | 11.421 | LYS-381 | 8.938 | >12.000 |
| ASP-46 | 4.116 | <0.000 | ARG-759 | 12.228 | >12.000 | GLU-382 | 5.675 | <0.000 |
| LYS-47 | 9.719 | >12.000 | TYR-762 | 14.594 | >12.000 | GLU-383 | 6.836 | <0.000 |
| CYS-49 | 9.248 | 9.348 | ARG-771 | 10.083 | >12.000 | LYS-384 | 10.693 | >12.000 |
| CYS-52 | 10.767 | >12.000 | ARG-776 | 10.282 | >12.000 | CYS-385 | 11.195 | >12.000 |
| GLU-53 | 5.888 | 0.730 | ARG-783 | 11.892 | >12.000 | GLU-386 | 5.553 | <0.000 |
| LYS-54 | 9.683 | 10.896 | ASP-797 | 4.647 | <0.000 | ARG-394 | 11.524 | >12.000 |
| GLU-56 | 6.258 | <0.000 | GLU-800 | 3.816 | <0.000 | LYS-395 | 10.171 | 10.585 |
| ASP-60 | 5.726 | <0.000 | HIS-807 | 5.207 | 3.368 | LYS-399 | 10.108 | 10.435 |
| ASP-61 | 5.047 | <0.000 | CYS-816 | 7.570 | 7.492 | ARG-402 | 12.151 | >12.000 |

| | | |
|---|---|---|
| LYS-63 9.179 >12.000 | ASP-823 2.599 <0.000 | ARG-410 11.826 >12.000 |
| HIS-64 5.720 9.050 | GLU-828 3.751 5.528 | GLU-412 5.136 0.902 |
| GLU-69 3.222 <0.000 | ASP-829 1.478 <0.000 | ASP-414 3.839 2.274 |
| ARG-70 8.644 >12.000 | TYR-830 10.911 >12.000 | GLU-415 4.126 3.367 |
| ARG-74 10.915 >12.000 | CYS-831 9.358 >12.000 | LYS-418 10.218 10.910 |
| GLU-75 4.124 <0.000 | LYS-835 9.380 >12.000 | GLU-421 4.361 1.315 |
| ARG-78 10.937 >12.000 | TYR-839 12.375 >12.000 | ASP-422 4.847 5.473 |
| CYS-79 7.155 8.371 | LYS-841 9.611 >12.000 | GLU-423 5.357 3.329 |
| LYS-81 9.191 >12.000 | GLU-844 4.631 3.112 | ASP-424 3.742 2.658 |
| CYS-82 6.821 >12.000 | GLU-845 5.283 5.206 | HID-428 6.245 6.107 |
| ASP-84 5.617 <0.000 | ASP-848 4.229 3.553 | LYS-430 9.983 >12.000 |
| CYS-87 5.793 >12.000 | ASP-850 3.751 <0.000 | ASP-432 4.667 2.550 |
| LYS-89 10.120 >12.000 | HIS-859 2.394 <0.000 | ASP-444 2.973 1.331 |
| CYS-91 6.640 8.917 | LYS-861 10.276 >12.000 | LYS-446 9.801 10.964 |
| HIS-94 5.713 0.409 | LYS-863 10.097 >12.000 | LYS-448 9.760 11.020 |
| ASP-96 2.303 <0.000 | ARG-867 11.666 >12.000 | GLU-449 4.386 4.021 |
| LYS-98 6.466 >12.000 | GLU-870 6.530 3.817 | LYS-455 10.222 11.077 |
| LYS-107 8.251 >12.000 | ASP-873 3.827 3.252 | ARG-458 11.797 >12.000 |
| TYR-109 12.293 >12.000 | LYS-874 10.222 11.604 | GLU-463 4.330 4.038 |
| TYR-110 11.905 >12.000 | LYS-875 10.158 >12.000 | ASP-465 3.138 2.845 |
| LYS-114 8.864 >12.000 | TYR-882 11.326 >12.000 | GLU-467 3.882 3.682 |
| ASP-119 5.422 <0.000 | GLU-884 4.874 2.698 | GLU-473 4.164 3.200 |
| CYS-126 6.601 >12.000 | ARG-886 11.628 >12.000 | ASP-481 1.702 <0.000 |
| CYS-130 6.202 >12.000 | LYS-887 10.151 >12.000 | GLU-491 3.011 <0.000 |
| ASP-134 4.533 <0.000 | LYS-888 10.369 11.315 | ASP-495 2.170 <0.000 |
| CYS-136 6.036 >12.000 | GLU-892 5.494 4.079 | LYS-497 8.758 >12.000 |
| CYS-140 6.456 5.546 | GLU-893 5.290 3.716 | TYR-502 10.836 >12.000 |
| TYR-143 10.672 >12.000 | LYS-894 9.690 >12.000 | HIS-504 5.309 4.651 |
| GLU-146 4.597 <0.000 | ARG-896 12.114 >12.000 | LYS-505 10.092 10.876 |
| GLU-147 5.224 <0.000 | LYS-898 10.086 >12.000 | TYR-506 11.696 >12.000 |
| GLU-161 4.307 2.857 | GLU-899 4.934 3.926 | TYR-511 11.474 >12.000 |

| | | |
|---|---|---|
| LYS-164 9.536 >12.000 | LYS-908 10.362 10.775 | LYS-518 9.969 >12.000 |
| ARG-172 13.132 >12.000 | ARG-909 11.520 >12.000 | GLU-520 5.117 2.691 |
| GLU-180 4.415 3.790 | CYS-911 10.130 11.404 | TYR-525 11.106 >12.000 |
| LYS-181 10.258 10.573 | LYS-915 10.243 11.792 | ASP-529 4.834 <0.000 |
| GLU-184 4.462 3.974 | ARG-916 11.939 >12.000 | ASP-532 3.498 2.437 |
| TYR-186 11.038 11.742 | LYS-922 10.179 11.304 | GLU-536 4.535 3.981 |
| LYS-189 10.624 >12.000 | ASP-923 4.440 2.744 | LYS-541 10.366 11.337 |
| CYS-202 10.875 >12.000 | LYS-927 10.571 10.938 | ARG-561 11.833 >12.000 |
| ARG-208 11.892 >12.000 | TYR-931 11.367 >12.000 | ARG-562 9.217 >12.000 |
| TYR-211 12.788 >12.000 | GLU-937 4.620 2.157 | GLU-565 4.518 3.413 |
| ASP-213 4.946 1.769 | GLU-942 4.228 3.649 | LYS-574 7.509 >12.000 |
| GLU-218 2.317 <0.000 | ASP-949 3.028 0.531 | ASP-579 3.747 2.915 |
| LYS-219 8.693 >12.000 | GLU-950 4.668 5.855 | LYS-580 9.498 >12.000 |
| GLU-221 4.132 3.561 | GLU-951 5.702 1.520 | ASP-581 4.111 3.531 |
| TYR-222 9.094 >12.000 | CYS-953 7.496 >12.000 | ARG-589 8.869 >12.000 |
| GLU-230 3.569 <0.000 | CYS-956 7.107 1.932 | ARG-592 12.314 >12.000 |
| ARG-235 9.353 >12.000 | LYS-958 7.074 >12.000 | TYR-600 10.708 >12.000 |
| TYR-238 11.091 >12.000 | CYS-959 6.653 <0.000 | GLU-611 5.412 <0.000 |
| ASP-239 4.261 0.642 | TYR-960 12.121 >12.000 | GLU-615 4.869 <0.000 |
| GLU-244 2.724 <0.000 | CYS-963 5.350 11.686 | LYS-616 8.766 >12.000 |
| GLU-246 4.835 3.403 | ASP-965 2.428 <0.000 | TYR-620 10.801 >12.000 |
| LYS-249 9.374 >12.000 | TYR-968 11.555 >12.000 | CYS-622 11.760 >12.000 |
| ASP-250 4.804 3.488 | ASP-974 2.649 0.129 | GLU-627 5.247 3.649 |
| LYS-254 10.056 >12.000 | GLU-976 4.583 3.605 | LYS-629 10.732 >12.000 |
| CYS-257 9.490 >12.000 | HIS-978 5.305 9.984 | ASP-631 5.100 2.340 |
| LYS-259 10.095 10.908 | ASP-984 4.401 3.121 | ASP-634 4.392 3.873 |
| GLU-265 4.494 3.918 | CYS-986 7.289 >12.000 | CYS-643 10.096 >12.000 |
| LYS-272 9.978 11.037 | CYS-989 6.958 8.920 | TYR-645 10.610 >12.000 |
| GLU-273 4.767 3.907 | CYS-992 5.732 <0.000 | LYS-647 10.186 10.183 |
| LYS-274 9.903 10.056 | CYS-996 7.455 10.503 | ASP-649 4.813 1.835 |
| TYR-276 11.567 >12.000 | ASP-1000 3.607 <0.000 | GLU-652 4.490 4.296 |

| | | |
|---|---|---|
| LYS-277 10.604 11.424 | CYS-1001 8.150 >12.000 | LYS-655 10.192 11.038 |
| GLU-287 3.240 2.792 | LYS-1002 3.228 >12.000 | LYS-656 9.621 >12.000 |
| LYS-290 9.997 11.006 | ARG-1003 10.588 >12.000 | GLU-658 5.018 4.231 |
| ASP-291 3.660 1.830 | ARG-1007 11.908 >12.000 | ASP-659 6.916 7.080 |
| ASP-300 4.078 3.428 | TYR-1011 12.126 >12.000 | ASP-663 5.385 1.746 |
| TYR-304 11.589 >12.000 | GLU-1012 4.464 3.817 | GLU-666 4.003 <0.000 |
| LYS-307 7.682 >12.000 | LYS-1014 9.453 11.036 | CYS-671 9.081 10.646 |
| ASP-308 2.913 1.506 | ARG-1015 12.068 >12.000 | HIS-673 4.876 5.868 |
| LYS-315 9.571 >12.000 | LEU-1017 3.812 2.839 | GLU-677 4.172 3.721 |
| LYS-318 10.523 >12.000 | **Chain-B** | ARG-678 11.497 >12.000 |
| CYS-322 8.439 8.884 | NTALA-2 7.303 7.567 | CYS-684 10.768 >12.000 |
| CYS-324 8.474 9.151 | LYS-7 10.092 11.514 | ASP-687 3.422 1.126 |
| HIS-325 6.353 6.228 | ASP-8 4.485 0.829 | GLU-689 3.988 3.859 |
| ARG-332 12.495 >12.000 | ASP-11 3.481 3.217 | ARG-692 11.261 >12.000 |
| ASP-342 2.251 <0.000 | GLU-13 4.381 4.529 | CYS-695 11.885 >12.000 |
| ASP-346 3.570 <0.000 | ARG-21 11.916 >12.000 | ARG-696 11.809 >12.000 |
| CYS-347 9.395 >12.000 | HIS-25 6.433 7.127 | ARG-699 13.391 >12.000 |
| ARG-353 12.182 >12.000 | CYS-29 9.514 10.390 | LYS-709 6.259 >12.000 |
| CYS-354 12.357 >12.000 | LYS-34 9.319 >12.000 | ASP-716 3.023 <0.000 |
| ARG-357 11.291 >12.000 | LYS-35 8.741 10.536 | ARG-722 11.522 >12.000 |
| ARG-358 10.450 >12.000 | ASP-37 4.901 0.807 | LYS-725 10.246 10.437 |
| ARG-364 11.344 >12.000 | LYS-38 8.850 >12.000 | GLU-726 5.090 2.468 |
| LYS-365 9.444 8.767 | LYS-39 10.086 11.352 | LYS-745 9.406 >12.000 |
| ARG-371 11.444 >12.000 | HIS-40 6.881 7.522 | ASP-747 4.039 <0.000 |
| GLU-375 3.182 <0.000 | LYS-42 9.927 >12.000 | LYS-758 10.182 11.247 |
| GLU-376 4.609 7.909 | ARG-43 13.227 >12.000 | ARG-759 12.513 >12.000 |
| GLU-378 4.472 <0.000 | ASP-46 3.757 <0.000 | TYR-762 13.151 >12.000 |
| LYS-381 8.961 >12.000 | LYS-47 9.620 >12.000 | ARG-771 9.652 >12.000 |
| GLU-382 5.890 <0.000 | CYS-49 9.602 9.668 | ARG-776 10.476 >12.000 |
| GLU-383 7.131 2.331 | CYS-52 9.631 10.695 | ARG-783 12.031 >12.000 |
| LYS-384 10.273 >12.000 | GLU-53 5.615 1.985 | ASP-797 4.369 <0.000 |

| | | |
|---|---|---|
| CYS-385 11.369 >12.000 | LYS-54 10.179 >12.000 | GLU-800 3.723 <0.000 |
| GLU-386 5.498 <0.000 | GLU-56 6.027 0.376 | HIS-807 4.995 2.920 |
| ARG-394 12.049 >12.000 | ASP-60 5.580 <0.000 | CYS-816 7.583 7.301 |
| LYS-395 10.185 10.533 | ASP-61 4.931 <0.000 | ASP-823 2.425 <0.000 |
| LYS-399 10.127 10.237 | LYS-63 9.065 >12.000 | GLU-828 3.787 5.495 |
| ARG-402 12.348 >12.000 | HIS-64 5.372 8.066 | ASP-829 1.085 <0.000 |
| ARG-410 12.126 >12.000 | GLU-69 3.049 <0.000 | TYR-830 10.028 >12.000 |
| GLU-412 5.336 3.049 | ARG-70 8.610 >12.000 | CYS-831 9.409 >12.000 |
| ASP-414 3.957 2.361 | ARG-74 10.614 >12.000 | LYS-835 9.531 >12.000 |
| GLU-415 4.611 3.124 | GLU-75 4.176 <0.000 | TYR-839 12.083 >12.000 |
| LYS-418 10.487 11.457 | ARG-78 11.185 >12.000 | LYS-841 9.345 >12.000 |
| GLU-421 5.204 2.049 | CYS-79 7.452 10.201 | GLU-844 4.858 3.424 |
| ASP-422 5.009 6.599 | LYS-81 9.384 >12.000 | GLU-845 4.934 4.976 |
| GLU-423 4.637 2.362 | CYS-82 6.829 >12.000 | ASP-848 4.196 3.298 |
| ASP-424 3.906 3.001 | ASP-84 5.188 <0.000 | ASP-850 3.373 <0.000 |
| HID-428 6.190 5.720 | CYS-87 6.049 >12.000 | HIS-859 4.635 5.008 |
| LYS-430 10.049 11.656 | LYS-89 10.015 >12.000 | LYS-861 10.211 >12.000 |
| ASP-432 4.726 2.443 | CYS-91 6.684 9.325 | LYS-863 9.993 >12.000 |
| ASP-444 3.025 1.501 | HIS-94 5.457 <0.000 | ARG-867 11.714 >12.000 |
| LYS-446 9.783 10.975 | ASP-96 1.971 <0.000 | GLU-870 5.014 3.574 |
| LYS-448 10.003 11.215 | LYS-98 6.142 >12.000 | ASP-873 3.845 3.093 |
| GLU-449 4.446 4.084 | LYS-107 8.064 >12.000 | LYS-874 10.169 11.356 |
| LYS-455 10.237 11.121 | TYR-109 11.951 >12.000 | LYS-875 10.049 >12.000 |
| ARG-458 11.879 >12.000 | TYR-110 12.459 >12.000 | TYR-882 11.538 >12.000 |
| GLU-463 4.395 4.226 | LYS-114 8.918 >12.000 | GLU-884 4.766 2.427 |
| ASP-465 2.478 1.902 | ASP-119 5.577 <0.000 | ARG-886 11.545 >12.000 |
| GLU-467 3.620 3.475 | CYS-126 7.598 >12.000 | LYS-887 10.081 >12.000 |
| GLU-473 4.169 3.252 | CYS-130 6.616 >12.000 | LYS-888 10.359 11.369 |
| ASP-481 1.697 <0.000 | ASP-134 4.849 <0.000 | GLU-892 5.120 3.584 |
| GLU-491 1.527 <0.000 | CYS-136 5.948 >12.000 | LYS-894 9.588 >12.000 |
| ASP-495 1.865 <0.000 | CYS-140 6.569 6.413 | ARG-896 12.028 >12.000 |

| | | |
|---|---|---|
| LYS-497 8.910 >12.000 | TYR-143 11.062 >12.000 | LYS-898 10.313 >12.000 |
| TYR-502 10.721 >12.000 | GLU-146 4.899 <0.000 | GLU-899 4.713 3.870 |
| HIS-504 5.529 5.057 | GLU-147 5.091 <0.000 | GLU-908 4.451 4.161 |
| LYS-505 10.020 10.886 | GLU-161 3.736 <0.000 | ARG-909 12.053 >12.000 |
| TYR-506 11.472 >12.000 | LYS-164 8.930 >12.000 | ASP-910 4.599 4.392 |
| TYR-511 11.391 >12.000 | ARG-172 12.886 >12.000 | CYS-911 11.082 >12.000 |
| LYS-518 10.405 >12.000 | GLU-180 4.340 3.833 | LYS-915 10.368 >12.000 |
| GLU-520 4.694 2.460 | LYS-181 10.263 10.540 | ARG-916 11.973 >12.000 |
| TYR-525 11.171 >12.000 | GLU-184 4.454 3.989 | LYS-922 10.215 11.315 |
| ASP-529 4.550 <0.000 | TYR-186 10.640 11.045 | ASP-923 3.488 1.495 |
| ASP-532 3.367 2.155 | LYS-189 10.554 11.622 | LYS-927 10.526 10.847 |
| GLU-536 4.520 4.070 | CYS-202 11.328 >12.000 | TYR-931 11.357 >12.000 |
| LYS-541 10.375 11.439 | ARG-208 12.173 >12.000 | GLU-937 5.376 3.135 |
| ARG-561 11.682 >12.000 | TYR-211 14.406 >12.000 | GLU-942 4.193 3.579 |
| ARG-562 9.172 >12.000 | ASP-213 5.427 2.277 | ASP-949 3.081 0.434 |
| GLU-565 4.588 3.426 | GLU-218 2.676 <0.000 | GLU-950 4.768 5.720 |
| LYS-574 7.621 >12.000 | LYS-219 8.580 >12.000 | GLU-951 5.644 0.188 |
| ASP-579 5.227 9.491 | GLU-221 4.261 3.937 | CYS-953 7.489 >12.000 |
| LYS-580 9.587 >12.000 | TYR-222 9.707 >12.000 | CYS-956 7.052 <0.000 |
| ASP-581 4.074 3.498 | GLU-230 4.551 0.334 | LYS-958 7.356 >12.000 |
| ARG-589 8.773 >12.000 | ARG-235 9.374 >12.000 | CYS-959 7.047 <0.000 |
| ARG-592 12.279 >12.000 | TYR-238 11.339 >12.000 | TYR-960 12.823 >12.000 |
| TYR-600 10.478 >12.000 | ASP-239 3.611 <0.000 | CYS-963 5.987 >12.000 |
| GLU-611 4.902 <0.000 | GLU-244 3.536 <0.000 | ASP-965 2.615 <0.000 |
| GLU-615 4.875 <0.000 | GLU-246 5.141 4.394 | TYR-968 11.129 >12.000 |
| LYS-616 8.724 >12.000 | LYS-249 9.644 >12.000 | ASP-974 3.421 1.243 |
| TYR-620 11.923 >12.000 | ASP-250 5.261 4.174 | GLU-976 4.826 3.810 |
| CYS-622 11.465 >12.000 | LYS-254 10.513 >12.000 | HIS-978 5.660 10.280 |
| GLU-627 3.900 1.193 | CYS-257 9.804 >12.000 | ASP-984 4.239 2.667 |
| LYS-629 10.761 >12.000 | LYS-259 10.095 10.949 | CYS-986 7.143 9.614 |
| ASP-631 4.956 2.740 | GLU-265 4.239 3.526 | CYS-989 7.164 6.202 |

| | | |
|---|---|---|
| ASP-634 4.364 3.885 | LYS-272 10.041 11.198 | CYS-992 6.714 >12.000 |
| CYS-643 9.773 >12.000 | GLU-273 4.637 3.923 | CYS-996 7.589 9.756 |
| TYR-645 10.854 11.797 | LYS-274 9.928 9.807 | ASP-1000 3.752 <0.000 |
| LYS-647 9.652 9.712 | TYR-276 11.844 >12.000 | CYS-1001 8.191 >12.000 |
| ASP-649 4.764 2.051 | LYS-277 10.656 11.399 | LYS-1002 4.736 >12.000 |
| GLU-652 4.299 4.117 | GLU-287 3.373 3.024 | ARG-1003 10.742 >12.000 |
| LYS-655 9.934 10.986 | LYS-290 9.694 10.778 | ARG-1007 11.792 >12.000 |
| LYS-656 9.203 11.476 | ASP-291 3.366 1.735 | TYR-1011 12.215 >12.000 |
| GLU-658 4.790 4.743 | ASP-300 3.895 3.517 | GLU-1012 4.463 3.777 |
| ASP-659 4.923 4.970 | TYR-304 11.644 >12.000 | LYS-1014 9.489 10.840 |
| ASP-663 5.380 1.857 | LYS-307 7.604 >12.000 | ARG-1015 11.964 >12.000 |
| GLU-666 3.471 <0.000 | ASP-308 2.671 1.112 | CTala-2035 4.036 2.880 |
| CYS-671 10.614 >12.000 | LYS-315 9.386 >12.000 | |
| HIS-673 5.824 6.323 | LYS-318 11.015 >12.000 | |
| GLU-677 4.057 2.963 | CYS-322 9.919 11.243 | |

**Table S3.** A representation of human DPD parameters and coordinate files for Model 1 (B3LYP/6-31G*): AMBER parameter file

##Model 1 (B3LYP/6-31G*) AMBER parameter file

REMARK GOES HERE, THIS FILE IS GENERATED BY MCPB.PY
MASS

| | | | |
|---|---|---|---|
| FE1 | 55.85 | | Fe ion |
| FE2 | 55.85 | | Fe ion |
| FE3 | 55.85 | | Fe ion |
| FE4 | 55.85 | | Fe ion |
| FE5 | 55.85 | | Fe ion |
| FE6 | 55.85 | | Fe ion |
| FE7 | 55.85 | | Fe ion |
| FE8 | 55.85 | | Fe ion |
| CysSG(78) | 32.06 | 2.900 | S in cystine |
| CysSG(81) | 32.06 | 2.900 | S in cystine |
| CysSG(86) | 32.06 | 2.900 | S in cystine |
| CysSG(90) | 32.06 | 2.900 | S in cystine |
| CysSG(129) | 32.06 | 2.900 | S in cystine |
| CysSG(135) | 32.06 | 2.900 | S in cystine |
| CysSG(139) | 32.06 | 2.900 | S in cystine |
| GnOE(155) | 16.00 | 0.434 | carbonyl group oxygen |
| S1 | 32.060 | 2.900 | S ion |
| S2 | 32.060 | 2.900 | S ion |
| S3 | 32.060 | 2.900 | S ion |
| S4 | 32.060 | 2.900 | S ion |
| S5 | 32.060 | 2.900 | S ion |
| S6 | 32.060 | 2.900 | S ion |
| S7 | 32.060 | 2.900 | S ion |
| S8 | 32.060 | 2.900 | S ion |

| BOND | (kcal/mol/Å) | (Å) | |
|---|---|---|---|
| CysSG(90)-FE1 | 44.6 | 2.3851 | Created by Seminario method using MCPB.py |
| S2-FE1 | 56.1 | 2.2759 | Created by Seminario method using MCPB.py |
| S2-FE3 | 57.3 | 2.3031 | Created by Seminario method using MCPB.py |
| S2-FE4 | 54.0 | 2.2876 | Created by Seminario method using MCPB.py |
| S3-FE1 | 55.1 | 2.2007 | Created by Seminario method using MCPB.py |
| S3-FE2 | 66.5 | 2.1837 | Created by Seminario method using MCPB.py |
| S3-FE4 | 55.6 | 2.1893 | Created by Seminario method using MCPB.py |
| S4-FE1 | 66.1 | 2.2277 | Created by Seminario method using MCPB.py |
| S4-FE2 | 62.0 | 2.2510 | Created by Seminario method using MCPB.py |
| S4-FE3 | 76.3 | 2.2375 | Created by Seminario method using MCPB.py |
| CysSG(135)-FE2 | 53.4 | 2.3262 | Created by Seminario method using MCPB.py |
| S1-FE2 | 54.7 | 2.2552 | Created by Seminario method using MCPB.py |
| S1-FE3 | 49.3 | 2.2649 | Created by Seminario method using MCPB.py |
| S1-FE4 | 50.5 | 2.2541 | Created by Seminario method using MCPB.py |
| GnOE(155)-FE3 | 60.4 | 1.9176 | Created by Seminario method using MCPB.py |
| CysSG(129)-FE4 | 36.4 | 2.4150 | Created by Seminario method using MCPB.py |
| CysSG(139)-FE5 | 36.4 | 2.3965 | Created by Seminario method using MCPB.py |
| S6-FE5 | 45.1 | 2.2410 | Created by Seminario method using MCPB.py |
| S6-FE7 | 45.3 | 2.2325 | Created by Seminario method using MCPB.py |
| S6-FE8 | 41.5 | 2.2698 | Created by Seminario method using MCPB.py |
| S7-FE5 | 64.3 | 2.2502 | Created by Seminario method using MCPB.py |

| | | | |
|---|---|---|---|
| S7-FE6 | 69.0 | 2.2387 | Created by Seminario method using MCPB.py |
| S7-FE8 | 61.8 | 2.2645 | Created by Seminario method using MCPB.py |
| S8-FE5 | 58.3 | 2.2124 | Created by Seminario method using MCPB.py |
| S8-FE6 | 51.2 | 2.2250 | Created by Seminario method using MCPB.py |
| S8-FE7 | 64.7 | 2.1906 | Created by Seminario method using MCPB.py |
| CysSG(86)-FE6 | 50.0 | 2.3515 | Created by Seminario method using MCPB.py |
| S5-FE6 | 66.3 | 2.2363 | Created by Seminario method using MCPB.py |
| S5-FE7 | 60.8 | 2.2371 | Created by Seminario method using MCPB.py |
| S5-FE8 | 57.0 | 2.2642 | Created by Seminario method using MCPB.py |
| CysSG(78)-FE7 | 40.2 | 2.3710 | Created by Seminario method using MCPB.py |
| CysSG(81)-FE8 | 37.1 | 2.4103 | Created by Seminario method using MCPB.py |
| C -GnOE(155) | 570.0 | 1.229 | JCC,7,(1986),230; AA,CYT,GUA,THY,URA |
| CT-CysSG(90) | 237.0 | 1.810 | changed from 222.0 based on methanethiol nmodes |
| CT-CysSG(135) | 237.0 | 1.810 | changed from 222.0 based on methanethiol nmodes |
| CT-CysSG(129) | 237.0 | 1.810 | changed from 222.0 based on methanethiol nmodes |
| CT-CysSG(139) | 237.0 | 1.810 | changed from 222.0 based on methanethiol nmodes |
| CT-CysSG(86) | 237.0 | 1.810 | changed from 222.0 based on methanethiol nmodes |
| CT-CysSG(78) | 237.0 | 1.810 | changed from 222.0 based on methanethiol nmodes |
| CT-CysSG(81) | 237.0 | 1.810 | changed from 222.0 based on methanethiol nmodes |

| ANGLE | (kcal mol/rad) | (degrees) | |
|---|---|---|---|
| C -GnOE(155)-FE3 | 75.86 | 130.30 | Created by Seminario method using MCPB.py |
| CT-CysSG(90)-FE1 | 106.68 | 103.98 | Created by Seminario method using MCPB.py |
| CT-CysSG(135)-FE2 | 100.90 | 107.39 | Created by Seminario method using MCPB.py |
| CT-CysSG(129)-FE4 | 97.96 | 96.66 | Created by Seminario method using MCPB.py |
| CT-CysSG(139)-FE5 | 94.60 | 105.21 | Created by Seminario method using MCPB.py |
| CT-CysSG(86)-FE6 | 101.58 | 107.61 | Created by Seminario method using MCPB.py |
| CT-CysSG(78)-FE7 | 100.02 | 108.65 | Created by Seminario method using MCPB.py |
| CT-CysSG(81)-FE8 | 106.65 | 103.60 | Created by Seminario method using MCPB.py |
| FE2-S3-FE1 | 75.11 | 68.39 | Created by Seminario method using MCPB.py |
| FE2-S4-FE1 | 80.79 | 66.76 | Created by Seminario method using MCPB.py |
| FE3-S2-FE1 | 23.20 | 69.98 | Created by Seminario method using MCPB.py |
| FE3-S4-FE1 | 26.76 | 72.04 | Created by Seminario method using MCPB.py |
| FE3-S4-FE2 | 40.49 | 66.60 | Created by Seminario method using MCPB.py |
| FE3-S1-FE2 | 47.73 | 66.08 | Created by Seminario method using MCPB.py |
| FE4-S2-FE1 | 53.67 | 65.65 | Created by Seminario method using MCPB.py |
| FE4-S2-FE3 | 20.95 | 64.15 | Created by Seminario method using MCPB.py |
| FE4-S3-FE1 | 29.62 | 68.59 | Created by Seminario method using MCPB.py |
| FE4-S3-FE2 | 61.07 | 68.33 | Created by Seminario method using MCPB.py |
| FE4-S1-FE2 | 78.37 | 66.00 | Created by Seminario method using MCPB.py |
| FE4-S1-FE3 | 53.87 | 65.30 | Created by Seminario method using MCPB.py |
| FE6-S7-FE5 | 63.39 | 66.89 | Created by Seminario method using MCPB.py |
| FE6-S8-FE5 | 77.31 | 67.77 | Created by Seminario method using MCPB.py |
| FE7-S6-FE5 | 42.52 | 67.16 | Created by Seminario method using MCPB.py |
| FE7-S8-FE5 | 64.00 | 68.38 | Created by Seminario method using MCPB.py |
| FE7-S8-FE6 | 42.93 | 68.59 | Created by Seminario method using MCPB.py |
| FE7-S5-FE6 | 35.00 | 67.59 | Created by Seminario method using MCPB.py |
| FE8-S6-FE5 | 61.00 | 67.08 | Created by Seminario method using MCPB.py |
| FE8-S6-FE7 | 44.11 | 66.60 | Created by Seminario method using MCPB.py |
| FE8-S7-FE5 | 46.02 | 67.02 | Created by Seminario method using MCPB.py |
| FE8-S7-FE6 | 47.60 | 70.85 | Created by Seminario method using MCPB.py |
| FE8-S5-FE6 | 36.63 | 70.89 | Created by Seminario method using MCPB.py |
| FE8-S5-FE7 | 67.61 | 66.62 | Created by Seminario method using MCPB.py |
| CysSG(90)-FE1-S2 | 41.30 | 111.43 | Created by Seminario method using MCPB.py |
| CysSG(90)-FE1-S3 | 63.82 | 115.56 | Created by Seminario method using MCPB.py |
| CysSG(90)-FE1-S4 | 65.80 | 106.49 | Created by Seminario method using MCPB.py |
| S2-FE1-S3 | 43.74 | 109.45 | Created by Seminario method using MCPB.py |

| | | | |
|---|---|---|---|
| S2-FE1-S4 | 28.56 | 103.16 | Created by Seminario method using MCPB.py |
| S2-FE3-S4 | 28.52 | 101.98 | Created by Seminario method using MCPB.py |
| S2-FE4-S3 | 60.52 | 109.42 | Created by Seminario method using MCPB.py |
| S3-FE1-S4 | 35.95 | 109.98 | Created by Seminario method using MCPB.py |
| S3-FE2-S4 | 69.35 | 109.74 | Created by Seminario method using MCPB.py |
| CysSG(135)-FE2-S3 | 52.84 | 115.18 | Created by Seminario method using MCPB.py |
| CysSG(135)-FE2-S4 | 75.62 | 109.23 | Created by Seminario method using MCPB.py |
| CysSG(135)-FE2-S1 | 47.43 | 105.58 | Created by Seminario method using MCPB.py |
| S1-FE2-S3 | 62.28 | 106.66 | Created by Seminario method using MCPB.py |
| S1-FE2-S4 | 34.47 | 110.31 | Created by Seminario method using MCPB.py |
| S1-FE3-S2 | 16.26 | 111.70 | Created by Seminario method using MCPB.py |
| S1-FE3-S4 | 22.23 | 110.45 | Created by Seminario method using MCPB.py |
| S1-FE4-S2 | 28.28 | 112.69 | Created by Seminario method using MCPB.py |
| S1-FE4-S3 | 59.25 | 106.50 | Created by Seminario method using MCPB.py |
| GnOE(155)-FE3-S2 | 34.09 | 115.71 | Created by Seminario method using MCPB.py |
| GnOE(155)-FE3-S4 | 59.86 | 109.31 | Created by Seminario method using MCPB.py |
| GnOE(155)-FE3-S1 | 51.72 | 107.58 | Created by Seminario method using MCPB.py |
| CysSG(129)-FE4-S2 | 53.37 | 112.42 | Created by Seminario method using MCPB.py |
| CysSG(129)-FE4-S3 | 46.08 | 111.58 | Created by Seminario method using MCPB.py |
| CysSG(129)-FE4-S1 | 64.85 | 104.01 | Created by Seminario method using MCPB.py |
| CysSG(139)-FE5-S6 | 53.22 | 109.82 | Created by Seminario method using MCPB.py |
| CysSG(139)-FE5-S7 | 45.83 | 111.05 | Created by Seminario method using MCPB.py |
| CysSG(139)-FE5-S8 | 51.92 | 109.60 | Created by Seminario method using MCPB.py |
| S6-FE5-S7 | 28.91 | 110.49 | Created by Seminario method using MCPB.py |
| S6-FE5-S8 | 36.64 | 105.86 | Created by Seminario method using MCPB.py |
| S6-FE7-S8 | 42.99 | 106.90 | Created by Seminario method using MCPB.py |
| S6-FE8-S7 | 20.07 | 108.94 | Created by Seminario method using MCPB.py |
| S7-FE5-S8 | 59.08 | 109.89 | Created by Seminario method using MCPB.py |
| S7-FE6-S8 | 37.10 | 109.86 | Created by Seminario method using MCPB.py |
| CysSG(86)-FE6-S7 | 45.25 | 109.29 | Created by Seminario method using MCPB.py |
| CysSG(86)-FE6-S8 | 63.88 | 116.82 | Created by Seminario method using MCPB.py |
| CysSG(86)-FE6-S5 | 53.36 | 107.61 | Created by Seminario method using MCPB.py |
| S5-FE6-S7 | 29.55 | 104.82 | Created by Seminario method using MCPB.py |
| S5-FE6-S8 | 44.49 | 107.73 | Created by Seminario method using MCPB.py |
| S5-FE7-S6 | 29.29 | 111.70 | Created by Seminario method using MCPB.py |
| S5-FE7-S8 | 76.82 | 108.93 | Created by Seminario method using MCPB.py |
| S5-FE8-S6 | 42.92 | 109.34 | Created by Seminario method using MCPB.py |
| S5-FE8-S7 | 16.77 | 103.08 | Created by Seminario method using MCPB.py |
| CysSG(78)-FE7-S6 | 62.02 | 113.79 | Created by Seminario method using MCPB.py |
| CysSG(78)-FE7-S8 | 55.31 | 113.14 | Created by Seminario method using MCPB.py |
| CysSG(78)-FE7-S5 | 62.04 | 102.35 | Created by Seminario method using MCPB.py |
| CysSG(81)-FE8-S6 | 47.02 | 116.54 | Created by Seminario method using MCPB.py |
| CysSG(81)-FE8-S7 | 51.70 | 110.21 | Created by Seminario method using MCPB.py |
| CysSG(81)-FE8-S5 | 53.95 | 107.88 | Created by Seminario method using MCPB.py |
| 2C-C -GnOE(155) | 80.0 | 120.40 | |
| CX-CT-CysSG(90) | 50.0 | 108.60 | AA cys (was CT-CT-SH) |
| CX-CT-CysSG(135) | 50.0 | 108.60 | AA cys (was CT-CT-SH) |
| CX-CT-CysSG(129) | 50.0 | 108.60 | AA cys (was CT-CT-SH) |
| CX-CT-CysSG(139) | 50.0 | 108.60 | AA cys (was CT-CT-SH) |
| CX-CT-CysSG(86) | 50.0 | 108.60 | AA cys (was CT-CT-SH) |
| CX-CT-CysSG(78) | 50.0 | 108.60 | AA cys (was CT-CT-SH) |
| CX-CT-CysSG(81) | 50.0 | 108.60 | AA cys (was CT-CT-SH) |
| N -C -GnOE(155) | 80.0 | 122.90 | AA general |
| CysSG(90)-CT-H1 | 50.0 | 109.50 | AA cyx changed based on NMA nmodes |
| CysSG(135)-CT-H1 | 50.0 | 109.50 | AA cyx changed based on NMA nmodes |
| CysSG(129)-CT-H1 | 50.0 | 109.50 | AA cyx changed based on NMA nmodes |
| CysSG(139)-CT-H1 | 50.0 | 109.50 | AA cyx changed based on NMA nmodes |

| | | | | |
|---|---|---|---|---|
| CysSG(86)-CT-H1 | 50.0 | 109.50 | AA cyx | changed based on NMA nmodes |
| CysSG(78)-CT-H1 | 50.0 | 109.50 | AA cyx | changed based on NMA nmodes |
| CysSG(81)-CT-H1 | 50.0 | 109.50 | AA cyx | changed based on NMA nmodes |

DIHE

| | | | | | |
|---|---|---|---|---|---|
| 2C-2C-C -GnOE(155) | 1 | 0.0 | 0.0 | 1.0 | |
| 2C-C -GnOE(155)-FE3 | 3 | 0.00 | 0.00 | 3.0 | Treat as zero by MCPB.py |
| C -GnOE(155)-FE3-S2 | 3 | 0.00 | 0.00 | 3.0 | Treat as zero by MCPB.py |
| C -GnOE(155)-FE3-S4 | 3 | 0.00 | 0.00 | 3.0 | Treat as zero by MCPB.py |
| C -GnOE(155)-FE3-S1 | 3 | 0.00 | 0.00 | 3.0 | Treat as zero by MCPB.py |
| CT-CysSG(90)-FE1-S2 | 3 | 0.00 | 0.00 | 3.0 | Treat as zero by MCPB.py |
| CT-CysSG(90)-FE1-S3 | 3 | 0.00 | 0.00 | 3.0 | Treat as zero by MCPB.py |
| CT-CysSG(90)-FE1-S4 | 3 | 0.00 | 0.00 | 3.0 | Treat as zero by MCPB.py |
| CT-CysSG(135)-FE2-S3 | 3 | 0.00 | 0.00 | 3.0 | Treat as zero by MCPB.py |
| CT-CysSG(135)-FE2-S4 | 3 | 0.00 | 0.00 | 3.0 | Treat as zero by MCPB.py |
| CT-CysSG(135)-FE2-S1 | 3 | 0.00 | 0.00 | 3.0 | Treat as zero by MCPB.py |
| CT-CysSG(129)-FE4-S2 | 3 | 0.00 | 0.00 | 3.0 | Treat as zero by MCPB.py |
| CT-CysSG(129)-FE4-S3 | 3 | 0.00 | 0.00 | 3.0 | Treat as zero by MCPB.py |
| CT-CysSG(129)-FE4-S1 | 3 | 0.00 | 0.00 | 3.0 | Treat as zero by MCPB.py |
| CT-CysSG(139)-FE5-S6 | 3 | 0.00 | 0.00 | 3.0 | Treat as zero by MCPB.py |
| CT-CysSG(139)-FE5-S7 | 3 | 0.00 | 0.00 | 3.0 | Treat as zero by MCPB.py |
| CT-CysSG(139)-FE5-S8 | 3 | 0.00 | 0.00 | 3.0 | Treat as zero by MCPB.py |
| CT-CysSG(86)-FE6-S7 | 3 | 0.00 | 0.00 | 3.0 | Treat as zero by MCPB.py |
| CT-CysSG(86)-FE6-S8 | 3 | 0.00 | 0.00 | 3.0 | Treat as zero by MCPB.py |
| CT-CysSG(86)-FE6-S5 | 3 | 0.00 | 0.00 | 3.0 | Treat as zero by MCPB.py |
| CT-CysSG(78)-FE7-S6 | 3 | 0.00 | 0.00 | 3.0 | Treat as zero by MCPB.py |
| CT-CysSG(78)-FE7-S8 | 3 | 0.00 | 0.00 | 3.0 | Treat as zero by MCPB.py |
| CT-CysSG(78)-FE7-S5 | 3 | 0.00 | 0.00 | 3.0 | Treat as zero by MCPB.py |
| CT-CysSG(81)-FE8-S6 | 3 | 0.00 | 0.00 | 3.0 | Treat as zero by MCPB.py |
| CT-CysSG(81)-FE8-S7 | 3 | 0.00 | 0.00 | 3.0 | Treat as zero by MCPB.py |
| CT-CysSG(81)-FE8-S5 | 3 | 0.00 | 0.00 | 3.0 | Treat as zero by MCPB.py |
| CX-CT-CysSG(90)-FE1 | 3 | 0.00 | 0.00 | 3.0 | Treat as zero by MCPB.py |
| CX-CT-CysSG(135)-FE2 | 3 | 0.00 | 0.00 | 3.0 | Treat as zero by MCPB.py |
| CX-CT-CysSG(129)-FE4 | 3 | 0.00 | 0.00 | 3.0 | Treat as zero by MCPB.py |
| CX-CT-CysSG(139)-FE5 | 3 | 0.00 | 0.00 | 3.0 | Treat as zero by MCPB.py |
| CX-CT-CysSG(86)-FE6 | 3 | 0.00 | 0.00 | 3.0 | Treat as zero by MCPB.py |
| CX-CT-CysSG(78)-FE7 | 3 | 0.00 | 0.00 | 3.0 | Treat as zero by MCPB.py |
| CX-CT-CysSG(81)-FE8 | 3 | 0.00 | 0.00 | 3.0 | Treat as zero by MCPB.py |
| H -N -C -GnOE(155) | 1 | 2.5 | 180.0 | -2.0 | JCC,7,(1986),230 |
| H -N -C -GnOE(155) | 1 | 2.0 | 0.0 | 1.0 | J.C.cistrans-NMA DE |
| FE1-CysSG(90)-CT-H1 | 3 | 0.00 | 0.00 | 3.0 | Treat as zero by MCPB.py |
| FE2-S3-FE1-S4 | 3 | 0.00 | 0.00 | 3.0 | Treat as zero by MCPB.py |
| FE2-CysSG(135)-CT-H1 | 3 | 0.00 | 0.00 | 3.0 | Treat as zero by MCPB.py |
| FE3-S2-FE1-S3 | 3 | 0.00 | 0.00 | 3.0 | Treat as zero by MCPB.py |
| FE3-S2-FE1-S4 | 3 | 0.00 | 0.00 | 3.0 | Treat as zero by MCPB.py |
| FE3-S1-FE2-S3 | 3 | 0.00 | 0.00 | 3.0 | Treat as zero by MCPB.py |
| FE3-S1-FE2-S4 | 3 | 0.00 | 0.00 | 3.0 | Treat as zero by MCPB.py |
| FE4-S2-FE1-S3 | 3 | 0.00 | 0.00 | 3.0 | Treat as zero by MCPB.py |
| FE4-S2-FE1-S4 | 3 | 0.00 | 0.00 | 3.0 | Treat as zero by MCPB.py |
| FE4-S2-FE3-S4 | 3 | 0.00 | 0.00 | 3.0 | Treat as zero by MCPB.py |
| FE4-S3-FE1-S4 | 3 | 0.00 | 0.00 | 3.0 | Treat as zero by MCPB.py |
| FE4-S3-FE2-S4 | 3 | 0.00 | 0.00 | 3.0 | Treat as zero by MCPB.py |
| FE4-S1-FE2-S3 | 3 | 0.00 | 0.00 | 3.0 | Treat as zero by MCPB.py |
| FE4-S1-FE2-S4 | 3 | 0.00 | 0.00 | 3.0 | Treat as zero by MCPB.py |
| FE4-S1-FE3-S2 | 3 | 0.00 | 0.00 | 3.0 | Treat as zero by MCPB.py |
| FE4-S1-FE3-S4 | 3 | 0.00 | 0.00 | 3.0 | Treat as zero by MCPB.py |
| FE4-CysSG(129)-CT-H1 | 3 | 0.00 | 0.00 | 3.0 | Treat as zero by MCPB.py |

| | | | | | |
|---|---|---|---|---|---|
| FE5-CysSG(139)-CT-H1 | 3 | 0.00 | 0.00 | 3.0 | Treat as zero by MCPB.py |
| FE6-S7-FE5-S8 | 3 | 0.00 | 0.00 | 3.0 | Treat as zero by MCPB.py |
| FE6-CysSG(86)-CT-H1 | 3 | 0.00 | 0.00 | 3.0 | Treat as zero by MCPB.py |
| FE7-S6-FE5-S7 | 3 | 0.00 | 0.00 | 3.0 | Treat as zero by MCPB.py |
| FE7-S6-FE5-S8 | 3 | 0.00 | 0.00 | 3.0 | Treat as zero by MCPB.py |
| FE7-S5-FE6-S7 | 3 | 0.00 | 0.00 | 3.0 | Treat as zero by MCPB.py |
| FE7-S5-FE6-S8 | 3 | 0.00 | 0.00 | 3.0 | Treat as zero by MCPB.py |
| FE7-CysSG(78)-CT-H1 | 3 | 0.00 | 0.00 | 3.0 | Treat as zero by MCPB.py |
| FE8-S6-FE5-S7 | 3 | 0.00 | 0.00 | 3.0 | Treat as zero by MCPB.py |
| FE8-S6-FE5-S8 | 3 | 0.00 | 0.00 | 3.0 | Treat as zero by MCPB.py |
| FE8-S6-FE7-S8 | 3 | 0.00 | 0.00 | 3.0 | Treat as zero by MCPB.py |
| FE8-S7-FE5-S8 | 3 | 0.00 | 0.00 | 3.0 | Treat as zero by MCPB.py |
| FE8-S7-FE6-S8 | 3 | 0.00 | 0.00 | 3.0 | Treat as zero by MCPB.py |
| FE8-S5-FE6-S7 | 3 | 0.00 | 0.00 | 3.0 | Treat as zero by MCPB.py |
| FE8-S5-FE6-S8 | 3 | 0.00 | 0.00 | 3.0 | Treat as zero by MCPB.py |
| FE8-S5-FE7-S6 | 3 | 0.00 | 0.00 | 3.0 | Treat as zero by MCPB.py |
| FE8-S5-FE7-S8 | 3 | 0.00 | 0.00 | 3.0 | Treat as zero by MCPB.py |
| FE8-CysSG(81)-CT-H1 | 3 | 0.00 | 0.00 | 3.0 | Treat as zero by MCPB.py |
| N -C -GnOE(155)-FE3 | 3 | 0.00 | 0.00 | 3.0 | Treat as zero by MCPB.py |
| CysSG(90)-FE1-S2-FE3 | 3 | 0.00 | 0.00 | 3.0 | Treat as zero by MCPB.py |
| CysSG(90)-FE1-S2-FE4 | 3 | 0.00 | 0.00 | 3.0 | Treat as zero by MCPB.py |
| CysSG(90)-FE1-S3-FE2 | 3 | 0.00 | 0.00 | 3.0 | Treat as zero by MCPB.py |
| CysSG(90)-FE1-S3-FE4 | 3 | 0.00 | 0.00 | 3.0 | Treat as zero by MCPB.py |
| CysSG(90)-FE1-S4-FE2 | 3 | 0.00 | 0.00 | 3.0 | Treat as zero by MCPB.py |
| CysSG(90)-FE1-S4-FE3 | 3 | 0.00 | 0.00 | 3.0 | Treat as zero by MCPB.py |
| S2-FE1-S3-FE2 | 3 | 0.00 | 0.00 | 3.0 | Treat as zero by MCPB.py |
| S2-FE1-S3-FE4 | 3 | 0.00 | 0.00 | 3.0 | Treat as zero by MCPB.py |
| S2-FE1-S4-FE2 | 3 | 0.00 | 0.00 | 3.0 | Treat as zero by MCPB.py |
| S2-FE1-S4-FE3 | 3 | 0.00 | 0.00 | 3.0 | Treat as zero by MCPB.py |
| S2-FE3-S4-FE1 | 3 | 0.00 | 0.00 | 3.0 | Treat as zero by MCPB.py |
| S2-FE3-S4-FE2 | 3 | 0.00 | 0.00 | 3.0 | Treat as zero by MCPB.py |
| S2-FE3-S1-FE2 | 3 | 0.00 | 0.00 | 3.0 | Treat as zero by MCPB.py |
| S2-FE4-S3-FE1 | 3 | 0.00 | 0.00 | 3.0 | Treat as zero by MCPB.py |
| S2-FE4-S3-FE2 | 3 | 0.00 | 0.00 | 3.0 | Treat as zero by MCPB.py |
| S2-FE4-S1-FE2 | 3 | 0.00 | 0.00 | 3.0 | Treat as zero by MCPB.py |
| S2-FE4-S1-FE3 | 3 | 0.00 | 0.00 | 3.0 | Treat as zero by MCPB.py |
| S3-FE1-S4-FE2 | 3 | 0.00 | 0.00 | 3.0 | Treat as zero by MCPB.py |
| S3-FE1-S4-FE3 | 3 | 0.00 | 0.00 | 3.0 | Treat as zero by MCPB.py |
| S3-FE2-S4-FE1 | 3 | 0.00 | 0.00 | 3.0 | Treat as zero by MCPB.py |
| S3-FE2-S4-FE3 | 3 | 0.00 | 0.00 | 3.0 | Treat as zero by MCPB.py |
| S3-FE4-S2-FE1 | 3 | 0.00 | 0.00 | 3.0 | Treat as zero by MCPB.py |
| S3-FE4-S2-FE3 | 3 | 0.00 | 0.00 | 3.0 | Treat as zero by MCPB.py |
| S3-FE4-S1-FE2 | 3 | 0.00 | 0.00 | 3.0 | Treat as zero by MCPB.py |
| S3-FE4-S1-FE3 | 3 | 0.00 | 0.00 | 3.0 | Treat as zero by MCPB.py |
| S4-FE2-S3-FE1 | 3 | 0.00 | 0.00 | 3.0 | Treat as zero by MCPB.py |
| S4-FE3-S2-FE1 | 3 | 0.00 | 0.00 | 3.0 | Treat as zero by MCPB.py |
| S4-FE3-S1-FE2 | 3 | 0.00 | 0.00 | 3.0 | Treat as zero by MCPB.py |
| CysSG(135)-FE2-S3-FE1 | 3 | 0.00 | 0.00 | 3.0 | Treat as zero by MCPB.py |
| CysSG(135)-FE2-S3-FE4 | 3 | 0.00 | 0.00 | 3.0 | Treat as zero by MCPB.py |
| CysSG(135)-FE2-S4-FE1 | 3 | 0.00 | 0.00 | 3.0 | Treat as zero by MCPB.py |
| CysSG(135)-FE2-S4-FE3 | 3 | 0.00 | 0.00 | 3.0 | Treat as zero by MCPB.py |
| CysSG(135)-FE2-S1-FE3 | 3 | 0.00 | 0.00 | 3.0 | Treat as zero by MCPB.py |
| CysSG(135)-FE2-S1-FE4 | 3 | 0.00 | 0.00 | 3.0 | Treat as zero by MCPB.py |
| S1-FE2-S3-FE1 | 3 | 0.00 | 0.00 | 3.0 | Treat as zero by MCPB.py |
| S1-FE2-S3-FE4 | 3 | 0.00 | 0.00 | 3.0 | Treat as zero by MCPB.py |
| S1-FE2-S4-FE1 | 3 | 0.00 | 0.00 | 3.0 | Treat as zero by MCPB.py |
| S1-FE2-S4-FE3 | 3 | 0.00 | 0.00 | 3.0 | Treat as zero by MCPB.py |

| | | | | | |
|---|---|---|---|---|---|
| S1-FE3-S2-FE1 | 3 | 0.00 | 0.00 | 3.0 | Treat as zero by MCPB.py |
| S1-FE3-S2-FE4 | 3 | 0.00 | 0.00 | 3.0 | Treat as zero by MCPB.py |
| S1-FE3-S4-FE1 | 3 | 0.00 | 0.00 | 3.0 | Treat as zero by MCPB.py |
| S1-FE3-S4-FE2 | 3 | 0.00 | 0.00 | 3.0 | Treat as zero by MCPB.py |
| S1-FE4-S2-FE1 | 3 | 0.00 | 0.00 | 3.0 | Treat as zero by MCPB.py |
| S1-FE4-S2-FE3 | 3 | 0.00 | 0.00 | 3.0 | Treat as zero by MCPB.py |
| S1-FE4-S3-FE1 | 3 | 0.00 | 0.00 | 3.0 | Treat as zero by MCPB.py |
| S1-FE4-S3-FE2 | 3 | 0.00 | 0.00 | 3.0 | Treat as zero by MCPB.py |
| GnOE(155)-C -2C-HC | 1 | 0.08 | 180.0 | -3.0 | Junmei et al, 1999 (HC-CT-C -O ) |
| GnOE(155)-C -2C-HC | 1 | 0.0 | 0.0 | -2.0 | |
| GnOE(155)-C -2C-HC | 1 | 0.8 | 0.0 | 1.0 | |
| GnOE(155)-FE3-S2-FE1 | 3 | 0.00 | 0.00 | 3.0 | Treat as zero by MCPB.py |
| GnOE(155)-FE3-S2-FE4 | 3 | 0.00 | 0.00 | 3.0 | Treat as zero by MCPB.py |
| GnOE(155)-FE3-S4-FE1 | 3 | 0.00 | 0.00 | 3.0 | Treat as zero by MCPB.py |
| GnOE(155)-FE3-S4-FE2 | 3 | 0.00 | 0.00 | 3.0 | Treat as zero by MCPB.py |
| GnOE(155)-FE3-S1-FE2 | 3 | 0.00 | 0.00 | 3.0 | Treat as zero by MCPB.py |
| GnOE(155)-FE3-S1-FE4 | 3 | 0.00 | 0.00 | 3.0 | Treat as zero by MCPB.py |
| CysSG(129)-FE4-S2-FE1 | 3 | 0.00 | 0.00 | 3.0 | Treat as zero by MCPB.py |
| CysSG(129)-FE4-S2-FE3 | 3 | 0.00 | 0.00 | 3.0 | Treat as zero by MCPB.py |
| CysSG(129)-FE4-S3-FE1 | 3 | 0.00 | 0.00 | 3.0 | Treat as zero by MCPB.py |
| CysSG(129)-FE4-S3-FE2 | 3 | 0.00 | 0.00 | 3.0 | Treat as zero by MCPB.py |
| CysSG(129)-FE4-S1-FE2 | 3 | 0.00 | 0.00 | 3.0 | Treat as zero by MCPB.py |
| CysSG(129)-FE4-S1-FE3 | 3 | 0.00 | 0.00 | 3.0 | Treat as zero by MCPB.py |
| CysSG(139)-FE5-S6-FE7 | 3 | 0.00 | 0.00 | 3.0 | Treat as zero by MCPB.py |
| CysSG(139)-FE5-S6-FE8 | 3 | 0.00 | 0.00 | 3.0 | Treat as zero by MCPB.py |
| CysSG(139)-FE5-S7-FE6 | 3 | 0.00 | 0.00 | 3.0 | Treat as zero by MCPB.py |
| CysSG(139)-FE5-S7-FE8 | 3 | 0.00 | 0.00 | 3.0 | Treat as zero by MCPB.py |
| CysSG(139)-FE5-S8-FE6 | 3 | 0.00 | 0.00 | 3.0 | Treat as zero by MCPB.py |
| CysSG(139)-FE5-S8-FE7 | 3 | 0.00 | 0.00 | 3.0 | Treat as zero by MCPB.py |
| S6-FE5-S7-FE6 | 3 | 0.00 | 0.00 | 3.0 | Treat as zero by MCPB.py |
| S6-FE5-S7-FE8 | 3 | 0.00 | 0.00 | 3.0 | Treat as zero by MCPB.py |
| S6-FE5-S8-FE6 | 3 | 0.00 | 0.00 | 3.0 | Treat as zero by MCPB.py |
| S6-FE5-S8-FE7 | 3 | 0.00 | 0.00 | 3.0 | Treat as zero by MCPB.py |
| S6-FE7-S8-FE5 | 3 | 0.00 | 0.00 | 3.0 | Treat as zero by MCPB.py |
| S6-FE7-S8-FE6 | 3 | 0.00 | 0.00 | 3.0 | Treat as zero by MCPB.py |
| S6-FE7-S5-FE6 | 3 | 0.00 | 0.00 | 3.0 | Treat as zero by MCPB.py |
| S6-FE8-S7-FE5 | 3 | 0.00 | 0.00 | 3.0 | Treat as zero by MCPB.py |
| S6-FE8-S7-FE6 | 3 | 0.00 | 0.00 | 3.0 | Treat as zero by MCPB.py |
| S6-FE8-S5-FE6 | 3 | 0.00 | 0.00 | 3.0 | Treat as zero by MCPB.py |
| S6-FE8-S5-FE7 | 3 | 0.00 | 0.00 | 3.0 | Treat as zero by MCPB.py |
| S7-FE5-S8-FE6 | 3 | 0.00 | 0.00 | 3.0 | Treat as zero by MCPB.py |
| S7-FE5-S8-FE7 | 3 | 0.00 | 0.00 | 3.0 | Treat as zero by MCPB.py |
| S7-FE6-S8-FE5 | 3 | 0.00 | 0.00 | 3.0 | Treat as zero by MCPB.py |
| S7-FE6-S8-FE7 | 3 | 0.00 | 0.00 | 3.0 | Treat as zero by MCPB.py |
| S7-FE8-S6-FE5 | 3 | 0.00 | 0.00 | 3.0 | Treat as zero by MCPB.py |
| S7-FE8-S6-FE7 | 3 | 0.00 | 0.00 | 3.0 | Treat as zero by MCPB.py |
| S7-FE8-S5-FE6 | 3 | 0.00 | 0.00 | 3.0 | Treat as zero by MCPB.py |
| S7-FE8-S5-FE7 | 3 | 0.00 | 0.00 | 3.0 | Treat as zero by MCPB.py |
| S8-FE6-S7-FE5 | 3 | 0.00 | 0.00 | 3.0 | Treat as zero by MCPB.py |
| S8-FE7-S6-FE5 | 3 | 0.00 | 0.00 | 3.0 | Treat as zero by MCPB.py |
| S8-FE7-S5-FE6 | 3 | 0.00 | 0.00 | 3.0 | Treat as zero by MCPB.py |
| CysSG(86)-FE6-S7-FE5 | 3 | 0.00 | 0.00 | 3.0 | Treat as zero by MCPB.py |
| CysSG(86)-FE6-S7-FE8 | 3 | 0.00 | 0.00 | 3.0 | Treat as zero by MCPB.py |
| CysSG(86)-FE6-S8-FE5 | 3 | 0.00 | 0.00 | 3.0 | Treat as zero by MCPB.py |
| CysSG(86)-FE6-S8-FE7 | 3 | 0.00 | 0.00 | 3.0 | Treat as zero by MCPB.py |
| CysSG(86)-FE6-S5-FE7 | 3 | 0.00 | 0.00 | 3.0 | Treat as zero by MCPB.py |
| CysSG(86)-FE6-S5-FE8 | 3 | 0.00 | 0.00 | 3.0 | Treat as zero by MCPB.py |

| | | | | | |
|---|---|---|---|---|---|
| S5-FE6-S7-FE5 | 3 | 0.00 | 0.00 | 3.0 | Treat as zero by MCPB.py |
| S5-FE6-S7-FE8 | 3 | 0.00 | 0.00 | 3.0 | Treat as zero by MCPB.py |
| S5-FE6-S8-FE5 | 3 | 0.00 | 0.00 | 3.0 | Treat as zero by MCPB.py |
| S5-FE6-S8-FE7 | 3 | 0.00 | 0.00 | 3.0 | Treat as zero by MCPB.py |
| S5-FE7-S6-FE5 | 3 | 0.00 | 0.00 | 3.0 | Treat as zero by MCPB.py |
| S5-FE7-S6-FE8 | 3 | 0.00 | 0.00 | 3.0 | Treat as zero by MCPB.py |
| S5-FE7-S8-FE5 | 3 | 0.00 | 0.00 | 3.0 | Treat as zero by MCPB.py |
| S5-FE7-S8-FE6 | 3 | 0.00 | 0.00 | 3.0 | Treat as zero by MCPB.py |
| S5-FE8-S6-FE5 | 3 | 0.00 | 0.00 | 3.0 | Treat as zero by MCPB.py |
| S5-FE8-S6-FE7 | 3 | 0.00 | 0.00 | 3.0 | Treat as zero by MCPB.py |
| S5-FE8-S7-FE5 | 3 | 0.00 | 0.00 | 3.0 | Treat as zero by MCPB.py |
| S5-FE8-S7-FE6 | 3 | 0.00 | 0.00 | 3.0 | Treat as zero by MCPB.py |
| CysSG(78)-FE7-S6-FE5 | 3 | 0.00 | 0.00 | 3.0 | Treat as zero by MCPB.py |
| CysSG(78)-FE7-S6-FE8 | 3 | 0.00 | 0.00 | 3.0 | Treat as zero by MCPB.py |
| CysSG(78)-FE7-S8-FE5 | 3 | 0.00 | 0.00 | 3.0 | Treat as zero by MCPB.py |
| CysSG(78)-FE7-S8-FE6 | 3 | 0.00 | 0.00 | 3.0 | Treat as zero by MCPB.py |
| CysSG(78)-FE7-S5-FE6 | 3 | 0.00 | 0.00 | 3.0 | Treat as zero by MCPB.py |
| CysSG(78)-FE7-S5-FE8 | 3 | 0.00 | 0.00 | 3.0 | Treat as zero by MCPB.py |
| CysSG(81)-FE8-S6-FE5 | 3 | 0.00 | 0.00 | 3.0 | Treat as zero by MCPB.py |
| CysSG(81)-FE8-S6-FE7 | 3 | 0.00 | 0.00 | 3.0 | Treat as zero by MCPB.py |
| CysSG(81)-FE8-S7-FE5 | 3 | 0.00 | 0.00 | 3.0 | Treat as zero by MCPB.py |
| CysSG(81)-FE8-S7-FE6 | 3 | 0.00 | 0.00 | 3.0 | Treat as zero by MCPB.py |
| CysSG(81)-FE8-S5-FE6 | 3 | 0.00 | 0.00 | 3.0 | Treat as zero by MCPB.py |
| CysSG(81)-FE8-S5-FE7 | 3 | 0.00 | 0.00 | 3.0 | Treat as zero by MCPB.py |

IMPR

| | | | | |
|---|---|---|---|---|
| X -X -C -GnOE(155) | 10.5 | 180. | 2. | JCC,7,(1986),230 |

NONB

| | | | |
|---|---|---|---|
| FE1 | 1.4090 | 0.0172100000 | IOD set for Fe2+ ion from Li et al. JCTC, 2013, 9, 2733 |
| FE2 | 1.4090 | 0.0172100000 | IOD set for Fe2+ ion from Li et al. JCTC, 2013, 9, 2733 |
| FE3 | 1.4090 | 0.0172100000 | IOD set for Fe2+ ion from Li et al. JCTC, 2013, 9, 2733 |
| FE4 | 1.4090 | 0.0172100000 | IOD set for Fe2+ ion from Li et al. JCTC, 2013, 9, 2733 |
| FE5 | 1.4090 | 0.0172100000 | IOD set for Fe2+ ion from Li et al. JCTC, 2013, 9, 2733 |
| FE6 | 1.4090 | 0.0172100000 | IOD set for Fe2+ ion from Li et al. JCTC, 2013, 9, 2733 |
| FE7 | 1.4090 | 0.0172100000 | IOD set for Fe2+ ion from Li et al. JCTC, 2013, 9, 2733 |
| FE8 | 1.4090 | 0.0172100000 | IOD set for Fe2+ ion from Li et al. JCTC, 2013, 9, 2733 |
| CysSG(78) | 2.0000 | 0.2500 | W. Cornell CH3SH and CH3SCH3 FEP's |
| CysSG(81) | 2.0000 | 0.2500 | W. Cornell CH3SH and CH3SCH3 FEP's |
| CysSG(86) | 2.0000 | 0.2500 | W. Cornell CH3SH and CH3SCH3 FEP's |
| CysSG(90) | 2.0000 | 0.2500 | W. Cornell CH3SH and CH3SCH3 FEP's |
| CysSG(129) | 2.0000 | 0.2500 | W. Cornell CH3SH and CH3SCH3 FEP's |
| CysSG(135) | 2.0000 | 0.2500 | W. Cornell CH3SH and CH3SCH3 FEP's |
| CysSG(139) | 2.0000 | 0.2500 | W. Cornell CH3SH and CH3SCH3 FEP's |
| GnOE(155) | 1.6612 | 0.2100 | OPLS |
| S1 | 2.0000 | 0.2500 | S ion |
| S2 | 2.0000 | 0.2500 | S ion |
| S3 | 2.0000 | 0.2500 | S ion |
| S4 | 2.0000 | 0.2500 | S ion |
| S5 | 2.0000 | 0.2500 | S ion |
| S6 | 2.0000 | 0.2500 | S ion |
| S7 | 2.0000 | 0.2500 | S ion |
| S8 | 2.0000 | 0.2500 | S ion |

**Table S4.** A representation of human DPD parameters and coordinate files for Model 2 (LSDA/LANL2DZ): AMBER_VFFDT parameter file

```
#This file is generated by the VFFDT program for AMBER.

MASS
S3              32.06500  2.90000          ; S
F3              55.84500  2.05000          ; Fe


BOND            (kcal/mol/Å)     (Å)
S3-F3           89.23755         2.22545   ; Averaged bond related parameters from VFFDT
F3-F3           59.53185         2.43332   ; Averaged bond related parameters from VFFDT
SG-F3            39.7711         2.3294    ; Averaged manually with SD(10.7658) and SD( 1.0804)
O-F3            24.9728          1.9340    ; Averaged manually with SD( 3.5472) and SD( 0.9834)

ANGL            (kcal mol/rad)   (degrees)
S3-F3-S3         39.52095        109.20    ; Averaged angle related parameters from VFFDT
S3-F3-F3        43.33559         73.06     ; Averaged angle related parameters from VFFDT
F3-S3-F3        26.86171         66.28     ; Averaged angle related parameters from VFFDT
F3-F3-F3        52.40760         60.00     ; Averaged angle related parameters from VFFDT
SG-F3-F3         38.8905         145.0629 ; Averaged manually with SD(11.4418) and SD( 9.6176)
SG-F3-S3        36.1370          113.2785 ; Averaged manually with SD( 6.7190) and SD( 9.1432)
H-SG-F3         23.5646          104.3280 ; Averaged manually with SD( 4.0419) and SD( 8.1811)
F3-O-C          33.8107          134.0050 ; Averaged manually with SD( 4.1504) and SD( 8.2116)
S3-F3-O         41.2264          115.3200 ; Averaged manually with SD( 5.1145) and SD( 8.6921)
O-F3-F3         56.4637          150.9850 ; Averaged manually with SD( 5.3204) and SD( 8.7809)
F3-SG-C         60.7114          112.8094 ; Averaged manually with SD(15.3253) and SD( 9.2780)
F3-SG-HS        0.0000           0.0000    ; Not calculated
F3-SG-2C        0.0000           0.0000    ; Not calculated
SG-F3-SG        0.0000           0.0000    ; Not calculated
O-F3-SG         0.0000           0.0000    ; Not calculated

DIHE
F3-S3-F3-S3     1  0.00000  0.00  1.00     ; Please check it manually.
S3-F3-S3-F3     1  0.00000  0.00  1.00     ; Please check it manually.
S3-F3-F3-S3     1  0.00000  0.00  1.00     ; Please check it manually.
S3-F3-F3-F3     1  0.00000  0.00  1.00     ; Please check it manually.
F3-S3-F3-F3     1  0.00000  0.00  1.00     ; Please check it manually.
F3-F3-S3-F3     1  0.00000  0.00  1.00     ; Please check it manually.
F3-F3-F3-F3     1  0.00000  0.00  1.00     ; Please check it manually.
F3-F3-F3-SG     1  0.00000  0.00  1.00     ; Please check it manually.
F3-F3-F3-O      1  0.00000  0.00  1.00     ; Please check it manually.
F3-F3-SG-2C     1  0.00000  0.00  1.00     ; Please check it manually.
F3-S3-F3-O      1  0.00000  0.00  1.00     ; Please check it manually.
F3-F3-SG-HS     1  0.00000  0.00  1.00     ; Please check it manually.
F3-S3-F3-SG     1  0.00000  0.00  1.00     ; Please check it manually.
F3-F3-O-C       1  0.00000  0.00  1.00     ; Please check it manually.
S3-F3-SG-HS     1  0.00000  0.00  1.00     ; Please check it manually.
S3-F3-F3-SG     1  0.00000  0.00  1.00     ; Please check it manually.
S3-F3-F3-O      1  0.00000  0.00  1.00     ; Please check it manually.
S3-F3-SG-2C     1  0.00000  0.00  1.00     ; Please check it manually.
S3-F3-O-C       1  0.00000  0.00  1.00     ; Please check it manually.
SG-F3-F3-O      1  0.00000  0.00  1.00     ; Please check it manually.
SG-F3-F3-SG     1  0.00000  0.00  1.00     ; Please check it manually.
```

```
HS-SG-F3-SG     1  0.00000   0.00  1.00          ; Please check it manually.
SG-F3-SG-2C     1  0.00000   0.00  1.00          ; Please check it manually.
O-F3-SG-2C      1  0.00000   0.00  1.00          ; Please check it manually.
O-F3-SG-HS      1  0.00000   0.00  1.00          ; Please check it manually.
C-O-F3-SG       1  0.00000   0.00  1.00          ; Please check it manually.


IMPR

NONB
  F3     1.4090  0.0172100000      IOD set for Fe2+ ion from Li et al. JCTC, 2013, 9, 2733
  S3     2.0000  0.2500           W. Cornell CH3SH and CH3SCH3 FEP's
```

**Table S5**. Listing of charge allocation to all the atoms interacting with the metal center (B3LYP/6-31G*)

| **Atom** | **Atomic charge** | **Atom** | **Atomic charge** | **Atom** | **Atomic** |
|---|---|---|---|---|---|
| FE1 | 0.489 | S1 | - 0.802 | GLN(156) | - 0.467 |
| FE2 | 0.725 | S2 | - 0.841 | CYS(136) | - 0.726 |
| FE3 | 0.622 | S3 | - 0.866 | CYS(130) | - 0.570 |
| FE4 | 0.614 | S4 | - 0.822 | CYS(91) | - 0.672 |
| FE5 | 0.553 | S5 | - 0.758 | CYS(82) | - 0.752 |
| FE6 | 0.705 | S6 | - 1.025 | CYS(87) | - 0.823 |
| FE7 | 0.667 | S7 | - 0.799 | CYS(140) | - 0.718 |
| FE8 | 0.638 | S8 | - 0.752 | CYS(79) | - 0.722 |

**A) Model 1**

**Table S6.** Comparison of **A** bond length, **B** internal and **C** external angles (Å) calculated with X-ray, DFT (B3LYP) and (LSDA/LANL2DZ) method for the molecular cluster model ([$Fe^{2+}{}_4S^{2-}{}_4$ (S-Cys)$_3$(S-Gln)]) 1026A of Native DPD protein.

**A**

| **Geometry** | **Bond length** *(Å)* | | | | | | |
|---|---|---|---|---|---|---|---|
| **Model System** | **$Fe^{2+}{}_4S^{2-}{}_4$(S-Cys)$_3$(O-Gln): 1026A Clusters** | | | | | | |
| **Bond** | **X-ray** | **QM** | | | | **AFTER-MD** | |
| **Bond description** | **1H7X** | **B3LYP (Model 1)** | | **LSDA/LANL2DZ (Model 2)** | | **Model 1** | **Model 2** |
| | **Bond length** *(Å)* | **Equilibrium bond length [req]** *(Å)* | **Force constant [Kr]** ***(kcal mol$^{-1}$ Å$^{-2}$)*** | **Equilibrium bond length [req]** *(Å)* | **Force constant [Kr]** *(kcal mol$^{-1}$ Å$^{-2}$)* | **Bond length** *(Å)* **Mean & SD** | **Bond length** *(Å)* **Mean & SD** |
| FE1-S2 | 2.51 | 2.28 | 56.10 | 2.22 | 89.23 | 2.28±0.16 | 2.22±0.21 |
| FE1-S3 | 2.42 | 2.20 | 55.10 | 2.22 | 89.23 | 2.20±0.17 | 2.23±0.14 |
| FE1-S4 | 2.58 | 2.22 | 66.10 | 2.22 | 89.23 | 2.24± 0.24 | 2.23± 0.25 |
| FE1-SG91 | 2.32 | 2.39 | 44.60 | 2.33 | 39.77 | 2.36±0.03 | 2.35±0.02 |
| FE2-S1 | 2.61 | 2.26 | 54.70 | 2.22 | 89.23 | 2.28±0.23 | 2.23±0.27 |
| FE2-S3 | 2.64 | 2.18 | 66.50 | 2.22 | 89.23 | 2.21±0.30 | 2.22±0.30 |
| FE2-S4 | 2.73 | 2.25 | 62.00 | 2.22 | 89.23 | 2.27±0.33 | 2.22±0.43 |
| FE2-SG136 | 2.37 | 2.33 | 53.40 | 2.33 | 39.77 | 2.36±0.01 | 2.34±0.02 |
| FE3-S1 | 2.43 | 2.26 | 49.30 | 2.22 | 89.23 | 2.25±0.13 | 2.23±0.14 |
| FE3-S2 | 2.40 | 2.30 | 57.30 | 2.22 | 89.23 | 2.30±0.07 | 2.23±0.12 |
| FE3-S4 | 2.56 | 2.24 | 76.30 | 2.22 | 89.23 | 2.23±0.23 | 2.23±0.23 |
| FE3-OE156 | 1.89 | 1.92 | 60.40 | 1.93 | 24.97 | 1.98±0.06 | 1.92±0.02 |
| FE4-S1 | 2.60 | 2.25 | 50.50 | 2.22 | 89.23 | 2.24±0.25 | 2.23±0.26 |
| FE4-S2 | 2.60 | 2.29 | 54.00 | 2.22 | 89.23 | 2.25±0.25 | 2.24±0.25 |
| FE4-S3 | 2.37 | 2.19 | 55.60 | 2.22 | 89.23 | 2.18±0.13 | 2.22±0.11 |
| FE4-SG130 | 2.36 | 2.40 | 36.40 | 2.33 | 39.77 | 2.38±0.01 | 2.34±0.01 |

[1]DFT: density functional theory, [2]B3LYP: Becke three-parameter hybrid exchange and Lee Yang Parr, [3]LSDA/LANL2DZ**:** Los Alamos double-zeta basis, [4]SD: standard deviation

**B**

| Geometry | Internal Angle ($^0$) | | | | | | |
|---|---|---|---|---|---|---|---|
| **Model System** | **$Fe^{2+}_4S^{2-}_4$(S-Cys)$_3$(O-Gln): 1026A Clusters** | | | | | | |
| **Angle** | **X-ray** | **QM** | | | | **AFTER-MD** | |
| **Angle description** | **1H7X** | **B3LYP (Model 1)** | | **LSDA/LANL2DZ (Model 2)** | | **Model 1** | **Model 2** |
| | **Average angle** ($^0$) | **Average equilibrium Angle [$\Theta_{eq}$]** ($^0$) | **Force constant [KΘ]** *(kcal mol$^{-1}$rad$^{-2}$)* | **Average equilibrium Angle [$\Theta_{eq}$]** ($^0$) | **Force constant [KΘ]** *(kcal mol$^{-1}$rad$^{-2}$)* | **Angle** $^{(0)}$ **Mean & SD** | **Angle** $^{(0)}$ **Mean & SD** |
| FE1-S2-FE3 | 68.12 | 69.98 | 23.20 | 66.28 | 26.86 | 58.20±7.01 | 69.05±0.66 |
| FE1-S2-FE4 | 65.87 | 65.65 | 53.67 | 66.28 | 26.86 | 64.32±0.29 | 69.61±2.64 |
| FE1-S3-FE2 | 64.61 | 68.39 | 75.11 | 66.28 | 26.86 | 64.33±0.20 | 68.93±3.05 |
| FE1-S3-FE4 | 65.86 | 68.59 | 29.62 | 66.28 | 26.86 | 67.08±0.86 | 67.81±1.37 |
| FE1-S4-FE2 | 68.72 | 66.76 | 80.79 | 66.28 | 26.86 | 60.34±5.93 | 70.09±0.96 |
| FE1-S4-FE3 | 67.60 | 72.04 | 26.76 | 66.28 | 26.86 | 63.80± 2.69 | 63.97± 2.56 |
| FE2-S1-FE3 | 68.15 | 66.08 | 47.73 | 66.28 | 26.86 | 67.27±0.62 | 68.49±0.24 |
| FE2-S3-FE4 | 71.15 | 68.33 | 61.07 | 66.28 | 26.86 | 65.35±4.10 | 69.07±1.47 |
| FE2-S4-FE3 | 64.60 | 66.60 | 40.49 | 66.28 | 26.86 | 61.15±2.43 | 67.96±2.38 |
| FE3-S1-FE4 | 71.92 | 65.300 | 53.87 | 66.28 | 26.86 | 62.67±6.54 | 67.05± 3.44 |
| FE3-S2-FE4 | 67.59 | 64.15 | 20.95 | 66.28 | 26.86 | 58.20±6.64 | 68.10±0.36 |
| FE4-S1-FE2 | 71.60 | 66.00 | 78.37 | 66.28 | 26.86 | 62.23±6.62 | 67.09±3.19 |
| S1-FE2-S3 | 102.67 | 106.66 | 62.28 | 109.21 | 39.52 | 114.04±8.04 | 105.95±2.31 |
| S1-FE2-S4 | 102.09 | 110.31 | 34.47 | 109.21 | 39.52 | 104.29± 1.55 | 105.23±2.22 |
| S1-FE3-S2 | 107.87 | 111.70 | 16.26 | 109.21 | 39.52 | 105.68±1.54 | 105.77±1.48 |
| S1-FE3-S4 | 107.47 | 110.45 | 22.23 | 109.21 | 39.52 | 104.41±2.16 | 108.21±0.52 |
| S1-FE4-S2 | 113.03 | 112.69 | 28.28 | 109.21 | 39.52 | 111.05±1.40 | 110.56±1.40 |
| S1-FE4-S3 | 101.97 | 106.50 | 59.25 | 109.21 | 39.52 | 105.86±2.75 | 109.33±5.20 |
| S2-FE1-S3 | 107.61 | 109.45 | 43.74 | 109.21 | 39.52 | 112.63±3.55 | 105.74±1.32 |
| S2-FE1-S4 | 102.08 | 103.16 | 18.56 | 109.21 | 39.52 | 111.25± 6.48 | 101.64±0.31 |
| S2-FE3-S4 | 107.74 | 101.98 | 18.52 | 109.21 | 39.52 | 110.49± 1.94 | 109.93±1.55 |
| S2-FE4-S3 | 106.71 | 109.42 | 60.52 | 109.21 | 39.52 | 108.75±1.44 | 108.36±1.17 |
| S3-FE1-S4 | 107.71 | 109.98 | 35.95 | 109.21 | 39.52 | 112.97±3.71 | 107.41±0.21 |
| S3-FE2-S4 | 105.38 | 109.74 | 69.35 | 109.21 | 39.52 | 109.61±2.99 | 105.77±0.27 |

[1]DFT: density functional theory, [2]B3LYP: Becke three-parameter hybrid exchange and Lee Yang Parr, [3]LSDA/LANL2DZ: Los Alamos double-zeta basis; [4]SD: standard deviation

C

| Geometry | External Angle (⁰) | | | | | | |
|---|---|---|---|---|---|---|---|
| Model System | $Fe_4S_4$(S-Cys)$_3$(O-Gln): 1026A | | | | | | |
| Bond calculations | X-ray | QM | | | | AFTER-MD | |
| Bond description | 1H7X | B3LYP (Model 1) | | GFN1-xTB (Model 2) | | Model 1 | Model 2 |
| | Angle (⁰) | Equilibrium Angle (⁰) | Force constant (kcal $mol^{-1}rad^{-2}$) | Equilibrium Angle (⁰) | Force constant (kcal $mol^{-1}rad^{-2}$) | Angle (⁰) Mean & SD | Angle (⁰) Mean & SD |
| C -OE155-FE3 | 117.29 | 130.30 | 75.86 | 115.32 | 41.23 | 115.29± 1.41 | 114.42±2.02 |
| CT-SG90-FE1 | 99.53 | 103.98 | 106.68 | 107.39 | 100.90 | 108.04±6.02 | 107.92±5.93 |
| CT-SG135-FE2 | 102.34 | 107.39 | 100.90 | 107.40 | 100.90 | 105.06±1.92 | 107.41±3.58 |
| CT-SG129-FE4 | 100.25 | 97.96 | 96.66 | 107.39 | 100.90 | 99.42±0.58 | 107.20±4.91 |
| N-C-OE155-H | 104.50 | 122.90 | 80.0 | 118.02 | 44.55 | 113.34±6.25 | 116.93±8.78 |
| SG90-CT-H | 107.85 | 109.50 | 50.0 | 104.33 | 23.56 | 100.25±5.37 | 103.85± 2.83 |
| SG135-CT-H | 108.36 | 109.50 | 50.0 | 104.33 | 23.56 | 109.45±0.77 | 103.96±3.11 |
| SG129-CT-H | 105.31 | 109.50 | 50.0 | 104.33 | 23.56 | 98.83±4.58 | 101.12±2.96 |
| SG90-FE1-S2 | 106.08 | 111.30 | 41.30 | 113.28 | 36.14 | 108.03±1.38 | 112.68±4.66 |
| SG90-FE1-S3 | 111.94 | 115.56 | 63.82 | 113.28 | 36.14 | 112.26±0.23 | 113.07±0.80 |
| SG90-FE1-S4 | 116.45 | 106.49 | 65.80 | 113.28 | 36.14 | 106.80±6.82 | 113.79±1.88 |
| SG135-FE2-S3 | 112.62 | 115.18 | 52.84 | 113.28 | 36.14 | 105.02±5.37 | 112.46±0.11 |
| SG135-FE2-S4 | 107.32 | 109.23 | 75.62 | 113.28 | 36.14 | 105.13±1.54 | 113.63± 4.46 |
| SG135-FE2-S1 | 100.34 | 105.58 | 47.43 | 113.28 | 36.14 | 108.76±5.95 | 113.69± 9.43 |
| OE155-FE3-S2 | 109.87 | 111.71 | 34.09 | 113.08 | 40.55 | 109.75±0.08 | 111.78±1.35 |
| OE155-FE3-S4 | 106.39 | 109.31 | 59.86 | 113.08 | 40.55 | 114.93±6.03 | 112.69±4.45 |
| OE155-FE3-S1 | 105.27 | 107.58 | 51.72 | 113.08 | 40.55 | 108.63±2.37 | 111.81±4.62 |
| SG129-FE4-S2 | 110.83 | 112.42 | 53.37 | 113.28 | 36.14 | 106.42±3.12 | 113.16±1.64 |
| SG129-FE4-S3 | 113.52 | 111.58 | 46.08 | 113.28 | 36.14 | 110.62±2.05 | 113.01±0.36 |
| SG129-FE4-S1 | 99.46 | 104.01 | 64.85 | 113.28 | 36.14 | 106.51±4.99 | 100.49±0.72 |

[1]DFT: density functional theory, [2]B3LYP: Becke three-parameter hybrid exchange and Lee Yang Parr, [3]GFN1-xTB DA/LANL2DZ**:** Los Alamos double-zeta basis; [4]SD: standard deviation

**Table S7.** Comparison of **A** bond length, **B** internal and **C** external angles (Å) calculated with X-ray, DFT (B3LYP) and (LSDA/LANL2DZ) method for the molecular cluster model ([$Fe_4S_4$(S-Cys)$_4$]) 1027A of native DPD protein

**A**

| **Geometry** | **Bond length (Å)** | | | | | | |
|---|---|---|---|---|---|---|---|
| **Model System** | **([$Fe_4S_4$(S-Cys)$_4$]): 1027A** | | | | | | |
| **Bond calculations** | **X-ray** | **QM** | | | | **AFTER-MD** | |
| **Bond description** | **1H7X** | **B3LYP (Model 1)** | | **LSDA/LANL2DZ (Model 2)** | | **Model 1** | **Model 2** |
| | **Bond length** *(Å)* | **Equilibrium Bond length** *(Å)* | **Force constant [Kr]** ***(kcal mol$^{-1}$ Å$^{-2}$)*** | **Equilibrium Bond length** *(Å)* | **Force constant [Kr]** ***(kcal mol$^{-1}$ Å$^{-2}$)*** | **Bond length** *(Å)* **Mean & SD** | **Bond length** *(Å)* **Mean & SD** |
| FE1_S2 | 2.63 | 2.24 | 45.10 | 2.22 | 89.23 | 2.23±0.28 | 2.22±0.29 |
| FE1_S3 | 2.43 | 2.25 | 64.30 | 2.22 | 89.23 | 2.24± 0.13 | 2.23±0.14 |
| FE1_S4 | 2.60 | 2.21 | 58.30 | 2.22 | 89.23 | 2.23±0.26 | 2.23±0.26 |
| FE1_SG140 | 2.30 | 2.40 | 36.10 | 2.33 | 39.77 | 2.39± 0.06 | 2.35±0.035 |
| FE2_S1 | 2.61 | 2.24 | 66.30 | 2.22 | 89.23 | 2.25±0.25 | 2.22± 0.27 |
| FE2_S3 | 2.55 | 2.24 | 69.00 | 2.22 | 89.23 | 2.23±0.22 | 2.22±0.23 |
| FE2_S4 | 2.25 | 2.23 | 51.20 | 2.22 | 89.23 | 2.24±0.01 | 2.23±0,01 |
| FE2_SG87 | 2.24 | 2.35 | 50.00 | 2.33 | 39.77 | 2.40±0.11 | 2.34±0.07 |
| FE3_S1 | 2.44 | 2.24 | 60.80 | 2.22 | 89.23 | 2.24± 0.14 | 2.23±0.15 |
| FE3_S2 | 2.38 | 2.23 | 45.30 | 2.22 | 89.23 | 2.32±0.04 | 2.23±0.11 |
| FE3_S4 | 2.25 | 2.19 | 64.70 | 2.22 | 89.23 | 2.19± 0.13 | 2.23±0.11 |
| FE3_SG79 | 2.30 | 2.37 | 40.20 | 2.33 | 39.77 | 2.40±0.07 | 2.35±0.04 |
| FE4_S1 | 2.55 | 2.27 | 41.50 | 2.20 | 89.23 | 2.26±0.21 | 2.22±0.23 |
| FE4_S2 | 2.25 | 2.27 | 61.80 | 2.22 | 89.23 | 2.28±0.02 | 2.23±0.01 |
| FE4_S3 | 2.60 | 2.26 | 57.00 | 2.22 | 89.23 | 2.27± 0.23 | 2.22±0.27 |
| FE4_SG82 | 2.40 | 2.41 | 37.10 | 2.33 | 39.77 | 2.37±0.02 | 2.33±0.05 |

[1]DFT: density functional theory, [2]B3LYP: Becke three-parameter hybrid exchange and Lee Yang Parr, [3]LSDA/LANL2DZ**:** Los Alamos double-zeta basis; [4]SD: standard deviation

**B**

| Geometry | Internal Angle $(^0)$ | | | | | | |
|---|---|---|---|---|---|---|---|
| **Model System** | **($[Fe_4S_4(S\text{-}Cys)_4]$): 1027A** | | | | | | |
| **Bond calculations** | **X-ray** | **QM** | | | | **MD** | |
| **Bond angle description** | **1H7X** | **B3LYP (Model 1)** | | **LSDA/LANL2DZ (Model 2)** | | **Model 1** | **Model 2** |
| | **Angle** $(^0)$ | **Equilibrium Angle** $(^0)$ | **Force constant** *(kcal $mol^{-1}rad^{-2}$)* | **Equilibrium Angle** $(^0)$ | **Force constant** *(kcal $mol^{-1}rad^{-2}$)* | **Angle** $(^0)$ **Mean & SD** | **Angle** $(^0)$ **Mean & SD** |
| FE1-S2-FE3 | 67.72 | 67.16 | 42.52 | 66.28 | 26.86 | 66.86±0.61 | 68.54±0.58 |
| FE1-S2-FE4 | 67.87 | 67.08 | 61.00 | 66.28 | 26.86 | 65.27±1.84 | 68.40±0.12 |
| FE1-S3-FE2 | 68.36 | 66.89 | 27.43 | 66.28 | 26.86 | 67.35±0.71 | 69.70±0.94 |
| FE1-S3-FE4 | 64.95 | 70.89 | 36.63 | 66.28 | 26.86 | 63.30±1.17 | 67.45±1.77 |
| FE1-S4-FE2 | 66.89 | 67.77 | 77.31 | 66.28 | 26.86 | 62.02±0.61 | 69.32±1.72 |
| FE1-S4-FE3 | 70.03 | 67.02 | 46.02 | 66.28 | 26.86 | 60.89±6.46 | 67.74±1.62 |
| FE2-S1-FE3 | 68.46 | 68.38 | 64.00 | 66.28 | 26.86 | 67.33±0.80 | 66.97±1.05 |
| FE2-S3-FE4 | 72.01 | 67.59 | 35.00 | 66.28 | 26.86 | 64.79±5.11 | 67.77±3.00 |
| FE2-S4-FE3 | 69.43 | 68.59 | 42.93 | 66.28 | 26.86 | 62.23±5.09 | 67.67±1.24 |
| FE3-S1-FE4 | 67.82 | 66.06 | 54.32 | 66.28 | 26.86 | 64.75±2.17 | 68.51±0.49 |
| FE3-S2-FE4 | 67.71 | 67.32 | 36.84 | 66.28 | 26.86 | 66.98±0.52 | 69.31±1.13 |
| FE4-S1-FE2 | 69.42 | 66.62 | 67.61 | 66.28 | 26.86 | 62.78±4.07 | 67.45±1.40 |
| S1-FE2-S3 | 107.69 | 104.82 | 49.55 | 109.21 | 39.52 | 108.31±0.43 | 109.66±1.39 |
| S1-FE2-S4 | 104.28 | 107.73 | 29.29 | 109.21 | 39.52 | 107.45±2.25 | 103.33±0.67 |
| S1-FE3-S2 | 117.43 | 111.70 | 44.49 | 109.21 | 39.52 | 117.93±0.35 | 109.85±5.36 |
| S1-FE3-S4 | 107.59 | 108.93 | 76.82 | 109.21 | 39.52 | 109.23±1.16 | 109.66±1.46 |
| S1-FE4-S2 | 106.71 | 109.34 | 42.92 | 109.21 | 39.52 | 108.31±1.13 | 110.03±2.35 |
| S1-FE4-S3 | 101.60 | 103.08 | 16.77 | 109.21 | 39.52 | 112.49±7.70 | 105.50±2.76 |
| S2-FE1-S3 | 107.70 | 110.60 | 28.91 | 109.21 | 39.52 | 109.76± 1.46 | 109.65±1.38 |
| S2-FE1-S4 | 105.51 | 105.86 | 36.64 | 109.21 | 39.52 | 111.75±4.41 | 108.32±1.99 |
| S2-FE3-S4 | 101.61 | 106.90 | 42.99 | 109.21 | 39.52 | 108.4±4.80 | 107.76±4.35 |
| S2-FE4-S3 | 106.67 | 108.94 | 20.07 | 109.21 | 39.52 | 110.93±2.24 | 108.45±0.49 |
| S3-FE1-S4 | 112.33 | 109.89 | 59.08 | 109.13 | 39.52 | 111.41±0.65 | 105.10±5.11 |
| S3-FE2-S4 | 107.37 | 109.86 | 37.10 | 109.21 | 39.52 | 104.05±2.35 | 109.54±1.53 |

[1]DFT: density functional theory, [2]B3LYP: Becke three-parameter hybrid exchange and Lee Yang Parr, [3]LSDA/LANL2DZ**:** Los Alamos double-zeta basis; [4]SD: standard deviation

**C**

| Geometry | External Angle (⁰) | | | | | | |
|---|---|---|---|---|---|---|---|
| Model System | ($[Fe_4S_4(S\text{-}Cys)_4]$): 1027A | | | | | | |
| Bond calculations | X-ray | QM | | | | AFTER-MD | |
| Bond description | 1H7X | B3LYP (Model 1) | | GFN1-xTB (Model 2) | | Model 1 | Model 2 |
| | Angle (⁰) | Equilibrium Angle (⁰) | Force constant (kcal $mol^{-1} rad^{-2}$) | Equilibrium Angle (⁰) | Force constant (kcal $mol^{-1} rad^{-2}$) | Angle (⁰) Mean & SD | Angle (⁰) Mean & SD |
| CT-SG78-FE3 | 123.43 | 108.65 | 100.02 | 115.32 | 41.23 | 108.76±10.37 | 114.21±6.52 |
| CT-SG139-FE1 | 101.35 | 105.21 | 94.60 | 107.39 | 100.90 | 105.13±2.67 | 107.30±4.21 |
| CT-SG86-FE2 | 102.43 | 107.61 | 107.61 | 107.39 | 100.90 | 110.52±6.43 | 107.14±4.04 |
| CT-SG81-FE4 | 100.25 | 103.60 | 106.65 | 107.39 | 100.90 | 105.82±3.94 | 107.56±5.17 |
| SG139-CT-H | 114.50 | 122.90 | 80.0 | 118.02 | 44.55 | 114.93 ± 0.30 | 116.43±1.36 |
| SG86-CT-H | 107.58 | 109.50 | 50.0 | 104.33 | 23.56 | 106,63±0.67 | 100.91±4.71 |
| SG78-CT-H | 108.61 | 109.50 | 50.0 | 104.33 | 23.56 | 112.28±2.60 | 103.65±3.51 |
| SG81-CT-H | 105.01 | 109.50 | 50.0 | 104.33 | 23.56 | 110.72±4.03 | 103.12±1.34 |
| SG139-FE1-S2 | 107.08 | 109.82 | 53.22 | 113.28 | 36.14 | 105.60±1.05 | 113.10±4.25 |
| SG139-FE1-S3 | 115.94 | 111.05 | 45.22 | 113.28 | 36.14 | 110.80± 3.63 | 113.16±1.97 |
| SG139-FE1-S4 | 112.45 | 109.60 | 51.92 | 113.28 | 36.14 | 112.52±0.05 | 112.64±0.13 |
| SG86-FE2-S3 | 112.62 | 109.29 | 45.25 | 113.28 | 36.14 | 106.83±4.09 | 111.18±1.01 |
| SG86-FE2-S4 | 117.32 | 116.82 | 63.88 | 113.28 | 36.14 | 112.53±3.38 | 113.37±2.80 |
| SG86-FE2-S1 | 100.34 | 107.61 | 53.36 | 113.28 | 36.14 | 99.83±0.36 | 111.84±8.13 |
| SG78-FE3-S2 | 109.78 | 113.79 | 62.02 | 113.28 | 36.14 | 110.21±0.30 | 112.49± 1.91 |
| SG78-FE3-S4 | 106.39 | 113.14 | 55.31 | 113.28 | 36.14 | 109.30±2.05 | 112.38± 4.24 |
| SG78-FE3-S1 | 105.27 | 102.35 | 62.04 | 113.28 | 36.14 | 108.77±2.47 | 113.18±5.59 |
| SG81-FE4-S2 | 110.83 | 116.54 | 47.02 | 113.28 | 36.14 | 112.78±1.38 | 113.33±1.77 |
| SG81-FE4-S3 | 113.26 | 110.21 | 51.70 | 113.28 | 36.14 | 107.39±4.15 | 112.98±0.20 |
| SG81-FE4-S4 | 110.66 | 107.95 | 53.93 | 113.28 | 36.14 | 109.50±0.82 | 111.56±0.64 |

[1]DFT: density functional theory, [2]B3LYP: Becke three-parameter hybrid exchange and Lee Yang Parr, [3]GFN1-xTB: eXtended TB; [4]SD: standard deviation

**Table S8.** Dihedral related force constants for X-ray and post-MD simulation for both models' clusters ([Fe4S4(S-Cys)$_3$(S-Gln)]) and ([$Fe_4S_4$(S-Cys)$_4$]) of Native

| $Fe_4S_4$ Clusters name | Geometry | Dihedral angles ($^0$) | | | | | | |
|---|---|---|---|---|---|---|---|---|
| | **Model System** | **$Fe_4S_4$(S-Cys)$_3$(O-Gln) and ([$Fe_4S_4$(S-Cys)$_4$]) Clusters** | | | | | | |
| | **Bond** | **X-ray** | **QM** | | | | **AFTER-MD** | |
| | **Bond** | **X-ray** | **B3YLP (Model 1)** | | **LSDA/LANL2DZ (Model 2)** | | **Model 1** | **Model 2** |
| | **Description** | **Average dihedral angle** *($^0$)* | **Equilibrium Bond angle** *($^0$)* | **Force constant** *(kcal/mol rad$^2$)* | **Equilibrium Bond angle** *($^0$)* | **Force constant (kcal/mol rad$^2$)** | **Bond angle ($^0$) Mean & SD** | **Bond angle ($^0$) Mean & SD** |
| **Cluster 1026_A** | S-FE-SG-CT | 67.43 | | | | | 74.87± 5.26 | 76.64±6.51 |
| | S-FE-OE-CT | 58.67 | | | | | 62.56± 2.75 | 63.42±3.35 |
| | CT-CT-FE-S | -112.78 | | | | | -105.36±5.25 | -105.79±4.94 |
| | CT-CT-FE-SG | -54.79 | | | | | -50.87±2.77 | -49.93±3.43 |

DPD protein

| | | | | | | | | |
|---|---|---|---|---|---|---|---|---|
| **Cluster 1027_A** | S-FE-SG-CT | 63.16 | | | | | 76.92±76.92 | 75.48± 8.71 |
| | S-FE-SG-CT | 56.23 | | | | | 63.54±5.17 | 63.74± 5.31 |
| | CT-CT-FE-S | -111.87 | | | | | -103.33±6.04 | -105.67±4.38 |
| | CT-CT-FE-SG | -56.42 | | | | | -51.76±3.30 | -51.93±3.1 |
| **Cluster 1028_B** | S-FE-SG-CT | 63.01 | | | | | 76.75±9.72 | 76.86±9.79 |
| | S-FE-SG-CT | 59.99 | | | | | 64.40±3.12 | 69.87±6.99 |
| | CT-CT-FE-S | -112.61 | | | | | -104.75±5.56 | -105.23±5.22 |
| | CT-CT-FE-SG | -55.28 | | | | | -50.87±3.12 | -50.11±3.66 |
| **Cluster 1028_B** | S-FE-SG-CT | 65.14 | | | | | 77.56±8.78 | 76.69±8.17 |
| | S-FE-SG-CT | 60.22 | | | | | 67.41±5.08 | 68.87±6.12 |
| | CT-CT-FE-S | -110.94 | | | | | -103.57± 5.21 | -105.38± 3.93 |
| | CT-CT-FE-SG | -58.23 | | | | | -54.08±2.93 | -52.39±4.13 |

[1]DFT: density functional theory, [2]B3LYP: Becke three-parameter hybrid exchange and Lee Yang Parr, [3]LSDA/LANL2DZ**:** Los Alamos double-zeta basis; [4]SD: standard deviation

**Table S9**. DPD *.pir* sequence file used for modeling human dihydropyramidine dehydrogenase structure based on pig crystal structure template and human target sequence

```
>P1;DPD_WT
sequence: DPD_WT: : : : :::0.00:0.00
MAPVLSKDSADIESILALNPRTQTHATLCSTSAKKLDKKHWKRNPDKNCFNCEKLENNFDDIKHTTL
GERGALREAMRCLKCADAPCQKSCPTNLDIKSFITSIANKNYYGAAKMIFSDNPLGLTCGMVCPTSDL
CVGGCNLYATEEGPINIGGLQQFAT
EVFKAMSIPQIRNPSLPPPEKMSEAYSAKIALFGAGPASISCASFLARLGYSDITIFEKQEYVGGLSTSEIP
QFRLPYDV
VNFEIELMKDLGVKIICGKSLSVNEMTLSTLKEKGYKAAFIGIGLPEPNKDAIFQGLTQDQGFYTSKDFL
PLVAKGSKAG
MCACHSPLPSIRGVVIVLGAGDTAFDCATSALRCGARRVFIVFRKGFVNIRAVPEEMELAKEEKCEFLP
FLSPRKVIVKG
GRIVAMQFVRTEQDETGKWNEDEDQMVHLKADVVISAFGSVLSDPKVKEALSPIKFNRWGLPEVDPE
TMQTSEAWVFAGG
DVVGLANTTVESVNDGKQASWYIHKYVQSQYGASVSAKPELPLFYTPIDLVDISVEMAGLKFINPFGL
ASATPATSTSMI
RRAFEAGWGFALTKTFSLDKDIVTNVSPRIIRGTTSGPMYGPGQSSFLNIELISEKTAAYWCQSVTELKA
DFPDNIVIAS
IMCSYNKNDWTELAKKSEDSGADALELNLSCPHGMGERGMGLACGQDPELVRNICRWVRQAVQIP
FFAKLTPNVTDIVSI
ARAAKEGGANGVTATNTVSGLMGLKSDGTPWPAVGIAKRTTYGGVSGTAIRPIALRAVTSIARALPG
FPILATGGIDSAE
SGLQFLHSGASVLQVCSAIQNQDFTVIEDYCTGLKALLYLKSIEELQDWDGQSPATVSHQKGKPVPRI
AELMDKKLPSFG
PYLEQRKKIIAENKIRLKEQNVAFSPLKRNCFIPKRPIPTIKDVIGKALQYLGTFGELSNVEQVVAMIDEE
MCINCGKCY
MTCNDSGYQAIQFDPETHLPTITDTCTGCTLCLSVCPIVDCIKMVSRTTPYEPKRGVPLSVNPVC.....*
```

```
>P1;1h7x_atm.pdb
structureX: 1h7x_atm.pdb::@::@:::0.00:0.00
-
APVLSKDVADIESILALNPRTQSHAALHSTLAKKLDKKHWKRNPDKNCFHCEKLENNFDDIKHTTLG
ERGALREAMRCL
KCADAPCQKSCPTHLDIKSFITSISNKNYYGAAKMIFSDNPLGLTCGMVCPTSDLCVGGCNLYATEEG
SINIGGLQQFAS
EVFKAMNIPQIRNPCLPSQEKMPEAYSAKIALLGAGPASISCASFLARLGYSDITIFEKQEYVGGLSTSEI
PQFRLPYDV
VNFEIELMKDLGVKIICGKSLSENEITLNTLKEEGYKAAFIGIGLPEPKTDDIFQGLTQDQGFYTSKDFLP
LVAKSSKAG
MCACHSPLPSIRGAVIVLGAGDTAFDCATSALRCGARRVFLVFRKGFVNIRAVPEEVELAKEEKCEFLP
FLSPRKVIVKG
GRIVAVQFVRTEQDETGKWNEDEDQIVHLKADVVISAFGSVLRDPKVKEALSPIKFNRWDLPEVDPET
MQTSEPWVFAGG
DIVGMANTTVESVNDGKQASWYIHKYIQAQYGASVSAKPELPLFYTPVDLVDISVEMAGLKFINPFGL
ASAAPTTSSSMI
RRAFEAGWGFALTKTFSLDKDIVTNVSPRIVRGTTSGPMYGPGQSSFLNIELISEKTAAYWCQSVTELK
ADFPDNIVIAS
IMCSYNKNDWMELSRKAEASGADALELNLSAPHGMGERGMGLACGQDPELVRNICRWVRQAVQIP
FFAKLTPNVTDIVSI
```

ARAAKEGGADGVTATNTVSGLMGLKADGTPWPAVGAGKRTTYGGVSGTAIRPIALRAVTTIARALP
GFPILATGGIDSAE
SGLQFLHSGASVLQVCSAVQNQDFTVIQDYCTGLKALLYLKSIEELQGWDGQSPGTESHQKGKPVPRI
AELMGKKLPNFG
PYLEQRKKIIAEEKMRLKEQNAAFPPLERKPFIPKKPIPAIKDVIGKALQYLGTFGELSNIEQVVAVIDEE
MCINCGKCY
MTCNDSGYQAIQFDPETHLPTVTDTCTGCTLCLSVCPIIDCIRMVSRTTPYEPKRGL--------.....*
